# Large-scale dissociations between views of objects, scenes, and reachable-scale environments in visual cortex

**DOI:** 10.1101/2020.02.20.956441

**Authors:** Emilie L. Josephs, Talia Konkle

## Abstract

Space-related processing recruits a network of brain regions separate from those recruited in object-related processing. This dissociation has largely been explored by contrasting views of navigable-scale spaces compared to close-up views of isolated objects. However, in naturalistic visual experience, we encounter spaces intermediate to these extremes, like the tops of desks and kitchen counters, which are not navigable but typically contain multiple objects. How are such reachable-scale views represented in the brain? In two functional neuroimaging experiments with human observers, we find evidence for a large-scale dissociation of reachable-scale views from both navigable scene views and close-up object views. Three brain regions were identified which showed a systematic response preference to reachable views, located in the posterior collateral sulcus, the inferior parietal sulcus, and superior parietal lobule. Subsequent analyses suggest that these three regions may be especially sensitive to the presence of multiple objects. Further, in all classic scene and object regions, reachable-scale views dissociated from both objects and scenes with an intermediate response magnitude. Taken together, these results establish that reachable-scale environments have a distinct representational signature from both scene and object views.

---

Scene-based and object-based representations form a major joint in the organization of the visual system. Scene-selective brain regions are broadly concerned with performing global perceptual analysis of a space (Epstein & Kanwisher, 1998; Kravitz, Peng & Baker, 2011; Lescroart & Gallant, 2019; Park, Brady, Greene, & Oliva, 2011), computing its navigational affordances (Bonner & Epstein, 2017; Kamps, Julian, Kubilius, Kanwisher & Dilks, 2016), and linking the present view to stored memory about the overall location (Vass & Epstein, 2017; Marchette, Vass, Ryan, & Epstein, 2015). In contrast, object-selective regions are engaged in representing bounded entities, and do so in a manner that is robust to confounding low-level contours and minor changes in size or position (Grill-Spector, Kourtzi, & Kanwisher, 2001; Grill-Spector, Kushnir, Edelman, Itzchak & Malach, 1998). Are these two systems, one for processing spatial-layout and another for bounded-objects, together sufficient to represent any view of the physical environment?

Consider views of reachable-scale environments—the countertops where we combine ingredients for a cake, or the worktables where we assemble the components of a circuit board. These views are intermediate in scale to scenes and objects, and are the locus of many everyday actions (Figure 1a). How are they represented in the visual system?

**Figure 1.**
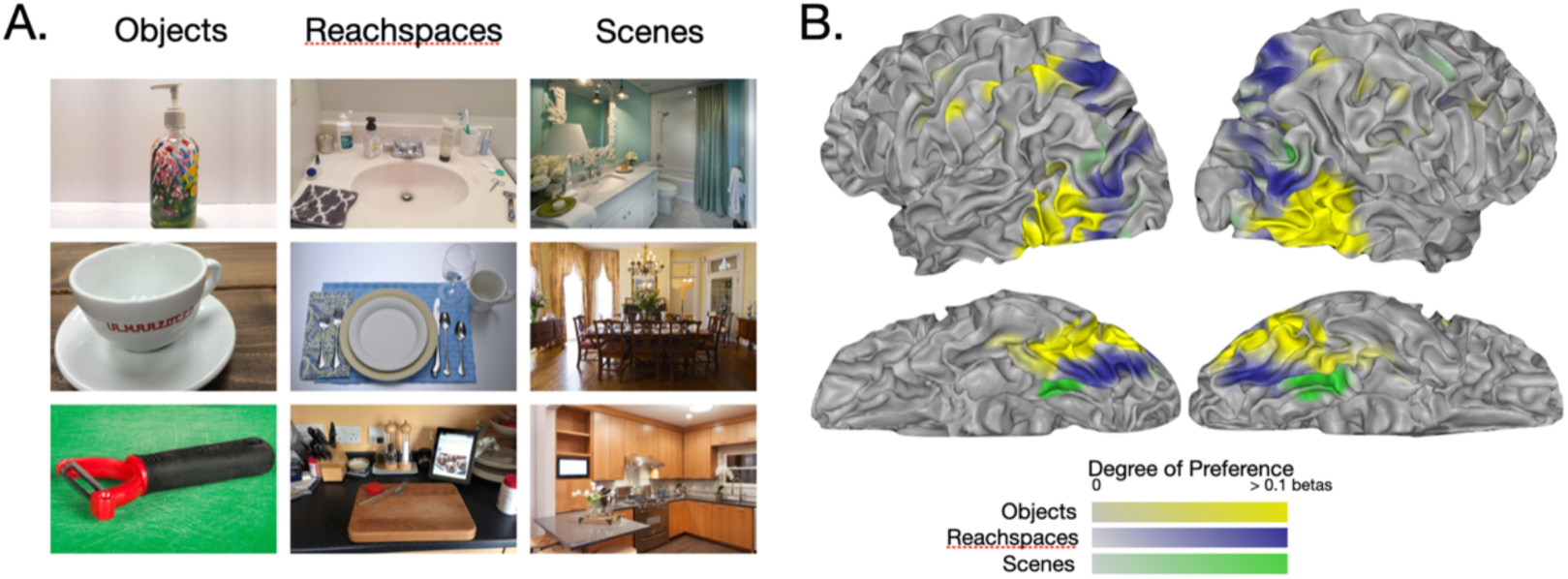
Experiment 1 stimuli and results. A) Examples of object, rcachspace and scene views. B) Preference mapping results. Colored regions have preference for either objects (yellow), reachspaces (blue) and scenes (green). Color saturation indicates the magnitude of the preference relative to the next most preferred category, operationalized as the difference between betas for that voxel.

One possibility is that reachable-scale environments are represented similarly to navigable-scale scenes, driving similar activations across the ventral and dorsal streams. Views of reachable environments are spatially extended, have 3D layout features, and need to be situated within larger environments, all of which are hypothesized functions of scene-selective regions. Indeed, this possibility is often assumed: reachable views have been used interchangeably with scene views in studies on scene processing (e.g. Epstein, Graham & Downing, 2003; Võ & Wolfe, 2013). However, everyday views of reachable-scale environments also prominently feature collections of multiple objects, and differ meaningfully from scenes by affording object-centered actions rather than navigation. Thus, a second possibility is that reachable-scale views will strongly drive object-preferring cortex.

A final, and not mutually-exclusive possibility, is that visual responses to reachable-scale environments might recruit distinct brain regions, separate from object- and scene-preferring cortex. There are both action-related and perception-related arguments for this hypothesis. First, it is clear that near-scale spaces have different behavioral demands than far-scale spaces (Previc, 1998; Grüsser, 1983, Rizzolatti & Camarda, 1987). Indeed, there are well-known motor dissociations between reach-related fronto-parietal circuits versus navigation-related medial networks (di Pellegrino & Làdavas, 2015; Graziano & Gross, 1995; Maguire, 2001). Second, statistics of visual images differ as a function of environment scale (Torralba & Oliva, 2002). We recently showed that the human perceptual system is sensitive to these differences: observers performing a visual search task were faster to find an image of a reachable environment among distractor scenes or objects than among distractor reachspaces, and vice versa (Josephs & Konkle, 2019). These results show that the scale of the depicted environment is a major factor in perceptual similarity computations.

These prior studies suggest that reachable-scale views dissociate from singleton objects views and navigable-scale scene views in *both* their input-related image statistics and output-related action requirements. Such visual input and behavioral output pressures have been proposed to be jointly essential for the large-scale functional clustering observed in visual cortex for different kinds of visual domains (e.g. faces, scenes; c.f. Op de Beeck, et al., 2019; Mahon & Caramazza, 2011; Konkle & Oliva, 2012; Arcaro & Livinsgtone, 2017; Conway, 2018). Thus, it is possible that views of reachable environments are distinct enough in form and purpose so as to require distinct visual processing regions.

In the present work, we examined how views of reachable-scale environments are represented in the human brain, using functional magnetic resonance imaging. We find clear evidence that reachspace representations dissociate from those of scenes and objects. Specifically, views of reachable environments elicited greater activity than both scenes and objects in regions of ventral and dorsal occipitoparietal cortex, in a manner that was robust to variations in luminance and global spatial frequency, and to variations in the semantic category depicted (e.g. kitchen reachspaces vs office reachspaces). Reachable-scale environments also elicited differential responses in classic object- and scene-preferring regions, generally leading to intermediate levels of activation between scene and object views. Regions preferring reachable-scale environments showed a peripheral eccentricity bias but also responded particularly strongly to images of multiple objects, a functional signature that is distinct from both scene and object regions. Taken together, these results suggest that the visual processing of near-scale environments is functionally dissociable from that of objects and scenes.

## Results

### Preferential responses to reachable-scale spaces in visual cortex

To examine the neural representation of reachable-scale environments compared to navigable-scale scenes and singleton objects, we created a stimulus set with images from each of the three environment types (see **Figure 1a**; **Supplementary Figure 1**). Object images depicted close-scale views of single objects (within 8-12 inches from the object) on their natural background. Reachable-scale images, which we will refer to as “reachspaces”, depicted near-scale environments that were approximately as deep as arm’s reach (3-4ft), and consisted of multiple small objects arrayed on a horizontal surface (Josephs & Konkle, 2019). cene images depicted views of the interior of rooms. Images were drawn from 6 different semantic categories (bar, bathroom, dining room, kitchen, office, art studio). Note that we use the term “environment scale” to refer to the distinction between conditions, but caution the reader against interpreting our results in terms of subjective distance only. Rather, differences observed here likely reflect differences across a constellation of dimensions that co-occur with scale (e.g. number of objects, number of surfaces, action affordances, and perceived reachability). Two stimulus sets were collected, with 90 images each (ImageSetA, ImageSetB; 30 images per environmental scale per set; see In-text Methods).

In Experiment 1, twelve participants viewed images of objects, reachspaces, and scenes, in a standard blocked fMRI design. All three stimulus conditions drove strong activations throughout visually-responsive cortex, particularly in early visual and posterior inferotemporal regions, with progressively weaker responses anteriorly through the ventral and dorsal stream (see **Supplementary Figure 2**). To help visualize the differences between these response topographies, voxels were colored according to the condition that most strongly activated them, with the saturation of the color reflecting the strength of the response preference (early visual regions excluded; see Supplementary Methods). This analysis revealed that different parts of cortex had reliable response preferences for each stimulus type, both at the group level (**Figure 1b**) and at the single-subject level (**Supplementary Figure 3**). Reachspace preferences (blue) were evident in three distinct zones: posterior ventral cortex, occipital-parietal cortex, and superior parietal cortex. These zones of preference lay adjacent to known object-preference zones (yellow), and scene-preference zones (green). Thus, while all three conditions extensively drive visual cortex, the activation landscapes differ in a systematic manner.

To estimate the magnitude of reachspace preferences, we defined reachspace-preferring regions of interest (ROIs) around the peaks in reachspace-preference appearing in anatomically consistent locations across subjects. Half of the data (activations from Image Set A) were submitted to a conjunction analysis to find voxels with a preference for reachspaces over objects, and reachspaces over scenes. This procedure yielded three reachspace-preferring ROIs: a ventral region (vROI), primarily located in the posterior collateral sulcus, an occipito-parietal region (opROI), variably located in the middle or superior occipital gyri, and a superior parietal region (spROI), in the anterior portion of the superior parietal lobe. Talairach (TAL) coordinates for these ROIs are given in **Supplementary Table 1**.

Next, we examined activation magnitude in the remaining half of the data (activations from Image Set B), and found that reachspace views elicited significantly higher activations than both scene and object views in all three ROIs (**Figure 2**, vROI: RS>O: t(8)=5.33, p<0.001, RS>S: t(8)=4.66, p=0.001; opROI: RS>O: t(6)=5.20, p=0.001, RS>S: t(6)=4.55, p=0.002; spROI: RS>O: t(7)=6.16, p<0.001, RS>S: t(7)=5.22, p=0.001). These results were also obtained when swapping the image set used to define the ROIs and test for activation differences: reachspaces elicited significantly greater activity than objects and scenes in all three ROIs (see **Supplementary Table 2** for all statistics). Finally, when the same data were broken out by semantic category, the reachspace categories elicited the highest overall responses (**Figure 2**).

**Figure 2:**
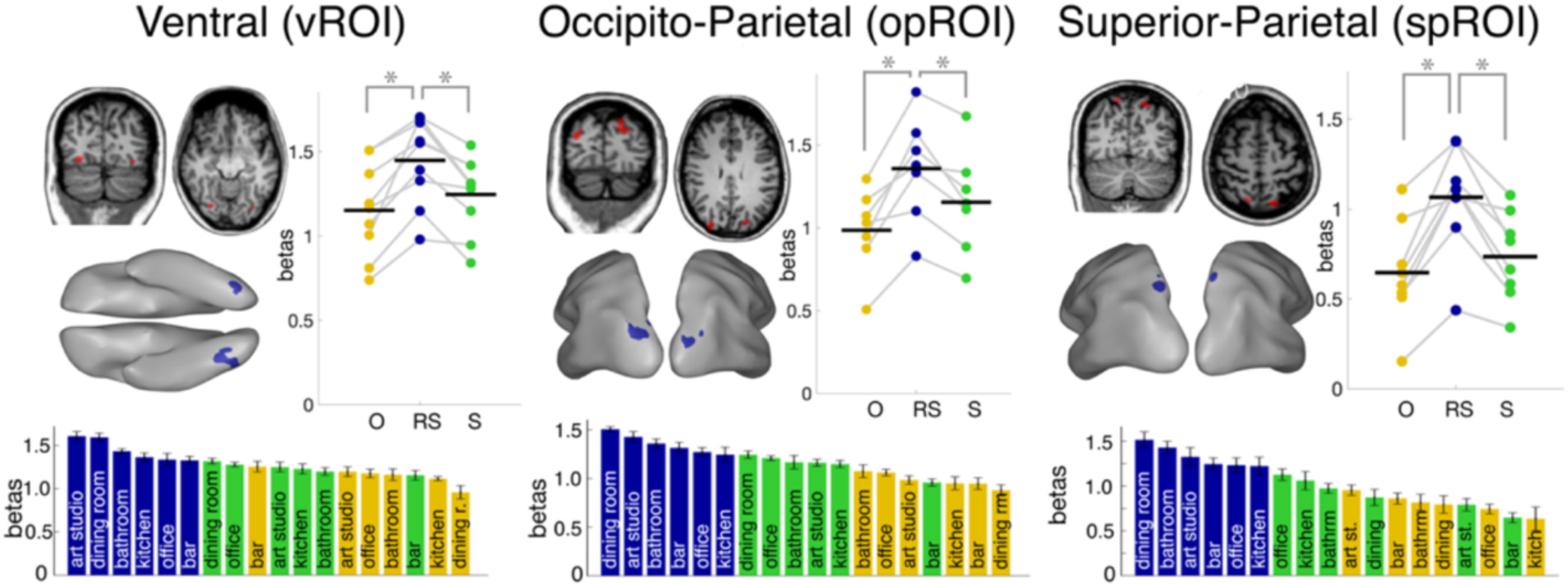
Locations and activations of reachspace-preferring ROIs. ROI locations are shown in the volume and on the inflated surface of an example subject. Bar plots show beta activations for objects, reachspaces, and scenes, averaged over semantic category (3-bar plot), or with semantic category displayed separately (18-bar plot). Error bars represent the within-subject standard error of the mean, and stars indicate statistical significance.

Taken together, these analyses show that there are portions of cortex with systematically stronger responses to images of reachable-scale environments than to the navigable-scale scenes and single object images.

### Low-level Control and Replication

In Experiment 2, we aimed to replicate the finding that reachspaces elicit greater activity than scenes and objects in some regions of cortex, and to test whether the response preferences for reachspaces are attributable to factors beyond very simple feature differences. Twelve participants (2 of whom had completed Experiment 1) viewed Image Set A (“original” images), and a version of Image Set B that was matched in mean luminance, contrast, and global spatial frequency content (“controlled” images; see **Figure 3a** and **Supplementary Figure 4** for examples).

**Figure 3:**
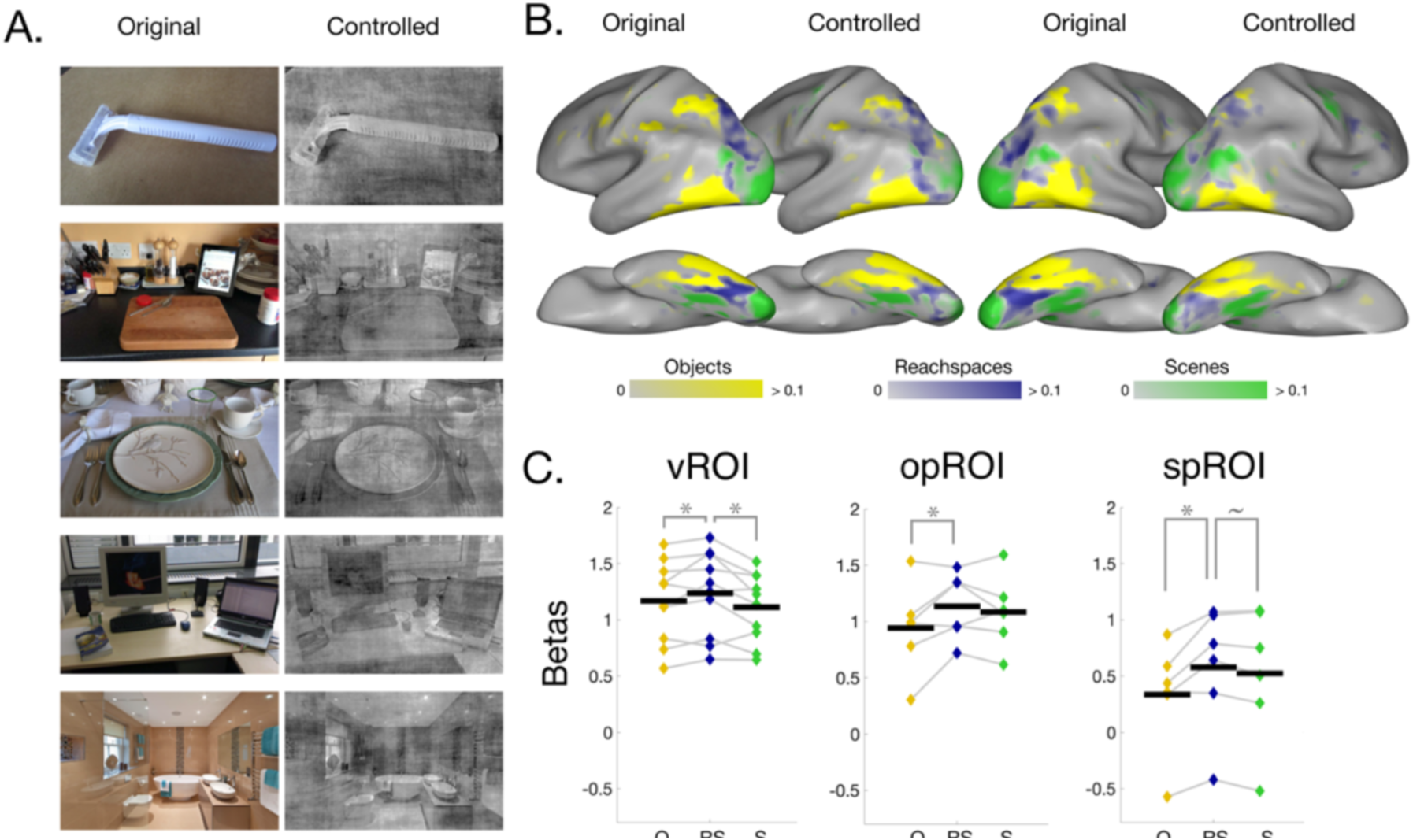
Stimuli and results for Experiment 2. A) Examples images showing effect of matching lumance. contrast, and global spatial frequency. B) A comparison of the group-average preference maps obtained for the original and controlled images, plotted on the same scale and projected onto an inflated brain. Color saturation indicates the magnitude of the preference relative to the next most preferred category. C) Activations in reachspace ROIs (defined in original images) in response to controlled images. Error bars represent the within-subject standard error of the mean, and stars indicate statistical significance.

Preference maps elicited by original and controlled images had highly similar spatial organization (**Figure 3b**; see **Supplementary Figure 5** for single-subject maps). At the group level, 69.9% of visually-responsive voxels had the same preference for either objects, reachspaces, or scene images, across original and controlled image formats (chance= 33.3%, 50.3%±1.5% match at the single subject level; see **Supplementary Figure 6**). Further, the topographies found in Experiment 2 with original images also match those found in Experiment 1 (67.4% of voxels in group-level preference maps had the same preference).

Additionally, the ROI results replicated with controlled images. Specifically, ROIs were defined in Experiment 2 subjects using original images, and activations were extracted for controlled images (**Figure 3c**). Preferential responses to reachspaces were generally maintained (vROI: RS>O: t(9)=2.08, p=0.034, RS>S: t(9)=2.72, p=0.012; opROI: RS>O: t(5)=2.38, p=0.032; spROI: RS>O: t(5)=3.61, p=0.008, RS>S: t(5)=2.02, p=0.05; RS>S in opROI was not significant: t(5)=0.79, p=0.234). Note that, in most of these ROIs, controlled images generally elicited lower overall activation magnitude than original images, and in some cases, the strength of the reachspace preference was slightly weaker than in the original image set (see **Supplementary Table 3**).

In summary, Experiment 2 found that the controlled image set elicited weaker but similar responses to object, reachspace, and scene images, indicating that these brain responses are not solely driven by stimulus differences in overall luminance, contrast, or global spatial frequency content. These results also provide a replication of Experiment 1, where the large-scale distribution of preferences for reachable-scale views was reproducible across a new sample of subjects.

### Responses to reachable-scale environments in scene- and object-preferring regions

We next evaluated reachspace-evoked activity in scene- and object-selective regions using data from both Experiment 1 (original images) and Experiment 2 (controlled images). All category-selective ROIs were defined using independent localizer runs (see Supplementary Methods).

In scene-preferring regions (parahippocampal place area, PPA; occipital place area, OPA; retrosplenial cortex, RSC), reachspaces elicited an intermediate level of activation for both original and controlled images (**Figure 4a**). That is, reachspace images evoked stronger activation than object images (original images: PPA: t(11)=11.29, p<0.001; OPA: t(10)=9.16, p>0.001, RSC: t(11)=9.15, p<0.001; controlled images: PPA: t(11)=8.43, p<0.001; OPA: t(10)=9.32, p<0.001; RSC: t(11)=5.24, p<0.001). Additionally, reachspace images evoked weaker activation than scene images, although this difference was marginal in OPA for original images (original image set: PPA: t(11)=4.50, p<0.001, OPA: t(10)=1.63, p=0.067; RSC: t(11)=6.80, p<0.001; controlled images: PPA: t(11)=9.69, p<0.001; OPA: t(10)=4.25, p=0.001, RSC: t(11)=6.48, p<0.001; see **Supplementary Table 2** for results in original images where the ROI-defining and activation-extracting runs were swapped; see **Supplementary Table 3** for comparisons of activations evoked by original and controlled images).

**Figure 4:**
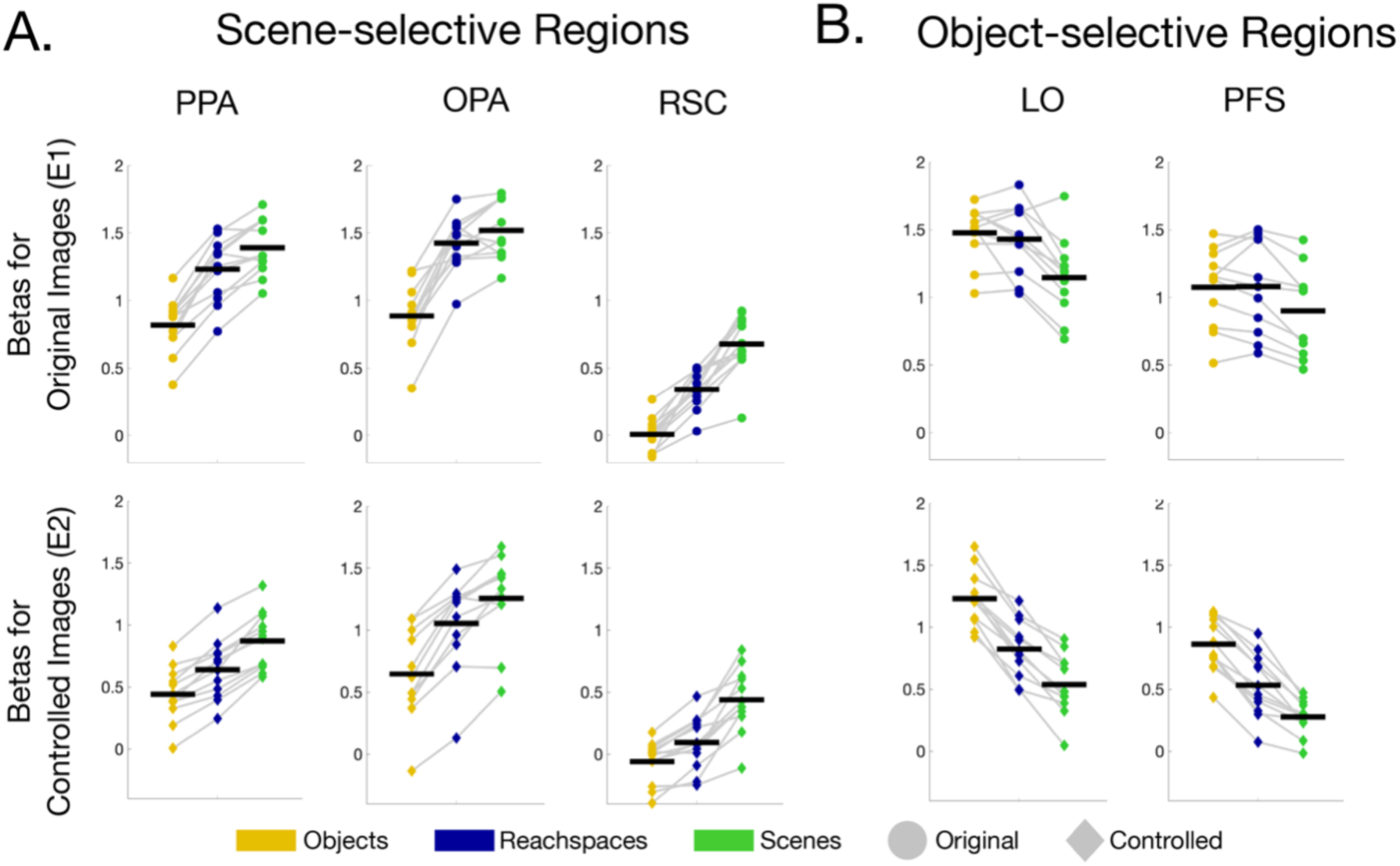
Result for classic category-selective ROIs. A) Univariate response to objects, scenes, and reachspaces in scene-selective regions, for both original and controlled images (i.e. Experiment 1 and Experiment 2 respectively). B) Same analysis for object-selective regions.

In object-preferring regions (lateral occipital, LO; and posterior fusiform sulcus, pFs), reachspaces also showed intermediate activation levels of activation in most of the comparisons (**Figure 4b**). Specifically, reachspace images elicited significantly more activity than scene images (original images: LO: t(10)=5.55, p<0.001; pFs: t(10)=4.86, p<0.001; controlled images: LO: t(11)=8.10, p<0.001; pFs: t(11)=6.04, p<0.001). Additionally, reachspace images elicited significantly weaker activation than objects for controlled images (LO: t(11)=11.20, p<0.001; pFs: t(11)=12.19, p<0.001), but showed a similar overall activation with object images in their original format (LO: t(10)=0.86, p=0.204; pFs: t(10)=-0.12, p=0.547; see **Supplementary Table 3** for all comparisons between activations to original vs. controlled images).

Taken together, these analyses show that reachspaces elicit an intermediate degree of activity in both scene- and object-preferring ROIs. These results provide further evidence that views of near-scale environments evoke different responses than both scene and objects images.

### Functional signatures of reachspace-preferring cortex

Next, we examined how object-, scene-, and reachspace-preferring ROIs differ in their broader functional signatures. We first report two opportunistic analyses from Experiment 1, which leverage stimulus conditions present in our localizer runs, then, we report data from a new Experiment 3, with planned functional signature analyses.

In our first opportunistic analysis, we examined the responses of regions with object, reachspace, and scene preferences to the eccentricity conditions present in the Experiment 1 retinotopy protocol (**Figure 5a;** see also **Supplementary Figure 7**). Reachspace-preferring regions showed a peripheral bias, which was significant at a conservative post-hoc statistical level for the ventral and occipital reachspace regions, but not in the superior parietal region (**Figure 5a**; vROI: t(8)=3.90, p=0.005; opROI: t(6)=4.82, p=0.003; spROI: t(7)=3.29, p=0.013; two tailed post-hoc paired ttest with Bonferri-corrected alpha=0.006). Similarly, scene regions were strongly peripherally-biased (PPA: t(11)=17.59, p<0.001; OPA: t(10)=9.27, p<0.001; RSC: t(11)=12.49, p<0.001). In contrast, object regions showed mixed biases, which did not reach significance after Bonferroni correction (LO: foveal bias, t(10)=2.68, p=0.023; pFs: peripheral bias, t(10)=2.26, p=0.047). These results show that regions which responded preferentially to reachspaces, like scene-selective regions, are most sensitive to peripherally presented stimuli.

**Figure 5.**
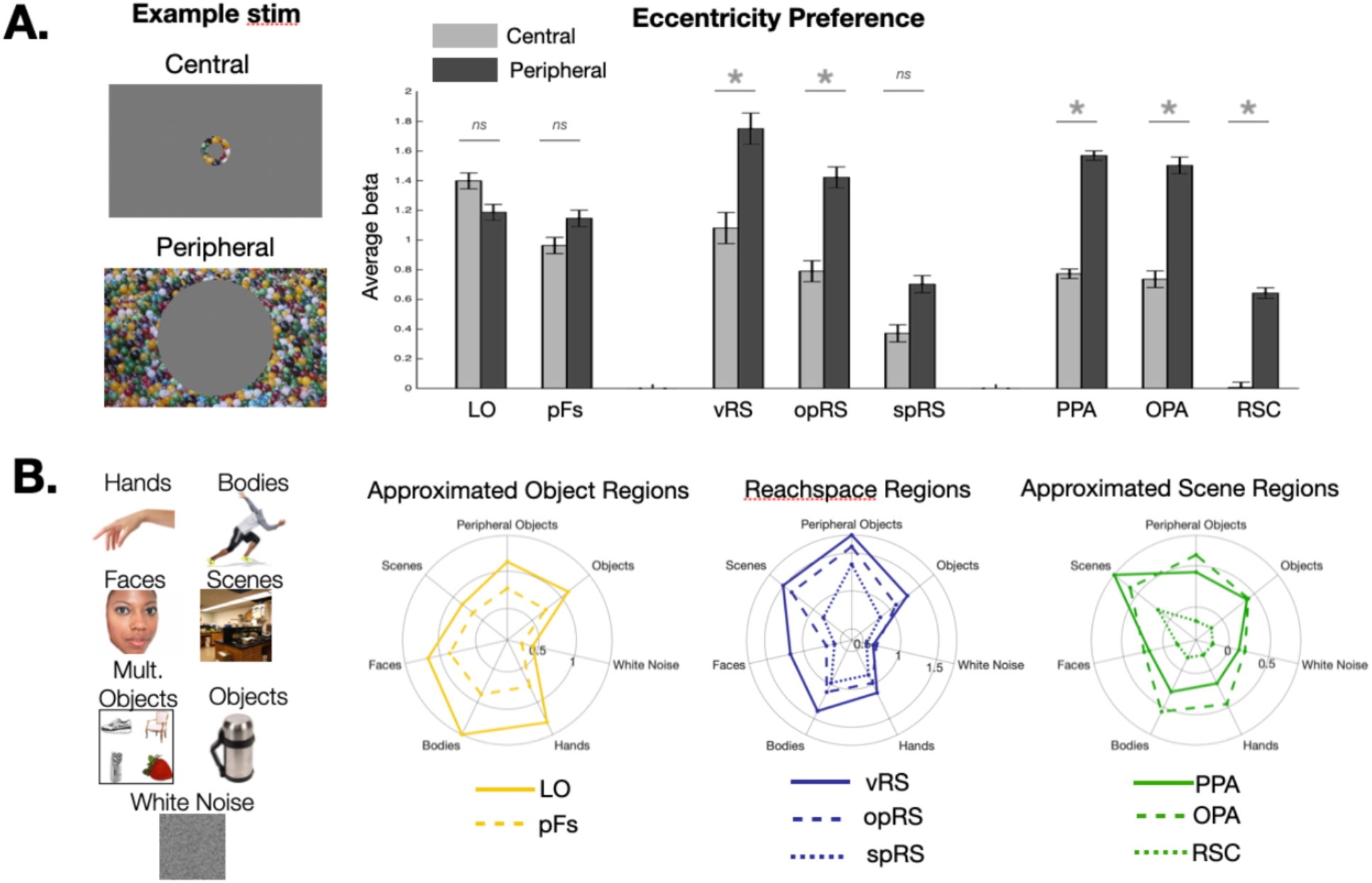
Response properties of reachspace regions, compared with scene and object regions. A) Stimuli and results for an analysis measuring the eccentricity bias of object, reachspace, and scene-preferring areas. Error bars show the within-subjects standard error of the mean, and stars indicate statistical significance. B) Stimuli and results for an analysis measuring the profile of responses across a range of categories for reachspaces regions, and regions corresponding to the anatomical locations of object- and scene-selective areas. Beta values are plotted for each condition in a polar plot; negative values were thresholded to 0 for visibility.

In our second opportunistic analysis, we investigated how ROIs differed in their response profile to a broad selection of categories present in the Experiment 1 localizer: faces, bodies, hands, objects, multiple objects, white noise, and scenes. Activations were extracted from reachspace-preferring ROIs. Since localizer runs were no longer available to define scene and object ROIs, these ROIs were approximated using a 9-mm radius sphere around TAL location that were estimated based on a literature review. Activations for all regions are plotted as fingerprint profiles in **Figure 5b**.

In all three reachspace-preferring ROIs, images of multiple objects elicited the highest activation (significant in ventral and superior parietal ROIs: vROI: Multiple Objects>Scenes, t(8)=3.49, p<0.01; spROI: Multiple Objects>Bodies, t(7)=5.54, p<0.01; marginal in opROI: Multiple Objects>Scenes t(6)=2.32, p=0.03). In contrast, scene and object preferring ROIs showed different functional signatures. The approximated PPA and RSC regions preferred scenes over all other conditions, including Multiple Objects (Scenes>Multiple objects in PPA: t(11)=12.02, p> 0.001; in RS: t(11)=7.87, p> 0.001; one-tailed paired t-test, post hoc alpha level=0.02). This difference was not significant for approximated OPA (t(11)=-0.18, p=0.57). Finally, approximated LO and pFs regions showed a maximal response to bodies, with broad tuning to hands, faces, object and multiple objects, and no differences between single objects and multiple objects in either ROI (LO: t(11)=-0.15, p=0.56; pFs: t(11)=-0.85, p=0.79). Overall, these exploratory analyses suggest that all three reachspace-preferring regions show a similar response profile with each other, in spite of their anatomically separation, and this profile is also distinct that of from scene and object preferring regions.

To test this formally, Experiment 3 probed responses in all ROIs to a broad range of conditions (**Figure 6**, stimuli in **Supplementary Figure 8**). These conditions included views of standard reachspaces, objects and scenes, as well as four different *multi-object conditions* (all depicting multiple objects with no background), and two different *minimal object conditions* (depicting near-scale spatial layouts with one or no objects). A final condition depicted *vertical* reachspaces, where the disposition of objects was vertical rather than horizontal (e.g. shelves, peg-boards). Experiment 3 was conducted in the same session as Experiment 2, and involved the same participants and functionally-defined ROIs. Activations from all conditions were extracted from each ROI and the fingerprints were compared.

**Figure 6:**
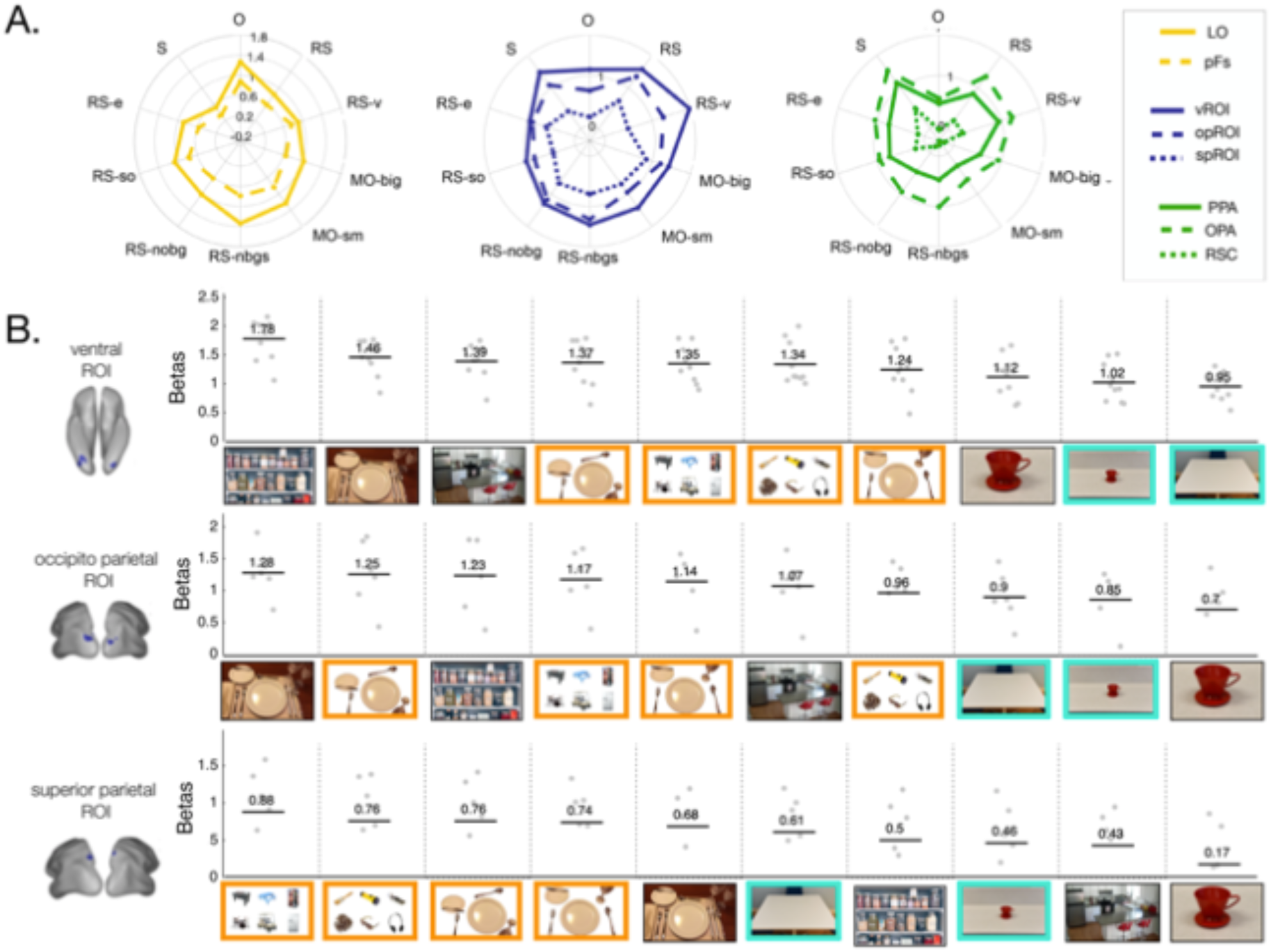
Experiment 3 results. A) Fingerprint profile of responses over all conditions in object, reachspace, and scene ROIs. O: singleton objects; RS: reachspaces; RS-v: vertical reachspaces; MO-b: multiple big objects, MO-sm: multiple small objects; RS-nbgs: reachspaces with no background, object positions scrambled; RS-nobg: Reachspace images with only objects with background removed; RS-so: reachspace images with only one object; RS-e: empty reachspace images. B) Responses in reachspace-preferring ROIs across all Experiment 3 conditions, plotted in order from highest to lowest activations. Images with orange borders indicate stimuli dominated by multiple objects, and images with teal borders highlight images of reachable space with low object content. See Supplement for all stimuli used in the experiment, in a larger format.

Across these 10 conditions, reachspace-preferring regions had a different fingerprint of activation than scene and object regions (**Figure 6a**). To test the significance of the difference in fingerprint profiles, responses across all conditions were averaged over the reachspace ROIs to create a reachspace-ROI fingerprint, then compared to the scene-ROI fingerprint (averaged over scene regions) and object-ROI fingerprint (averaged over object regions) using a 2-way ANOVA. An omnibus test of ROI-type (object, reachspace or scene) by condition revealed an ROI type-by-condition interaction (F(9, 329)=65.55, p<0.001), showing that the patterns of activations across the 10 conditions varied as a function of ROI-type. This difference held when reachspace-ROIs were compared to scene- and object-ROIs separately (interaction effect for reachspace versus scene ROIs: F(9, 219)=32.20, p<0.001; for reachspace versus object ROIs: F(9, 219)=47.89, p<0.001). These results further corroborate the conclusion that reachspace-preferring regions have a distinct representational signatures than object- and scene-preferring cortex.

Examining this response profile in more detail, in all three reachspace-preferring ROIs, responses were higher to all multi-object conditions (**Figure 7**, orange outline) than to empty reachspaces and to single-object reachspaces (blue outline). To quantify this, responses to all multi-object conditions were averaged, as were responses to empty reachspaces and single-object reachspaces, and two resulting activation levels were compared with a post hoc t-test (vROI: t(9)=7.75, p <0.01; opROI: t(5)=4.57, p <0.01; spROI: t(5)=4.50, p < 0.01). This pattern of data suggests that the presence of multiple easily-individuated objects may be particularly critical for driving the strong response to typical reachspace images relative to full-scale scenes, where object content may be less prominent than layout information. In contrast, in scene-preferring regions, the empty reachspace images generated higher responses than multiple object arrays, although this difference was marginal in OPA (**Supplementary Figure 9**; PPA: t(11)=-8.16, p < 0.01; OPA: t(10)=-1.49, p=0.08; RSC: t(11)=-7.28, p < 0.01). This result is consistent with prior work showing scene-regions strongly prefer empty full-scale rooms over multiple objects, and generally reflect responses to spatial layout (Epstein & Kanwisher, 1998).

**Figure 7.**
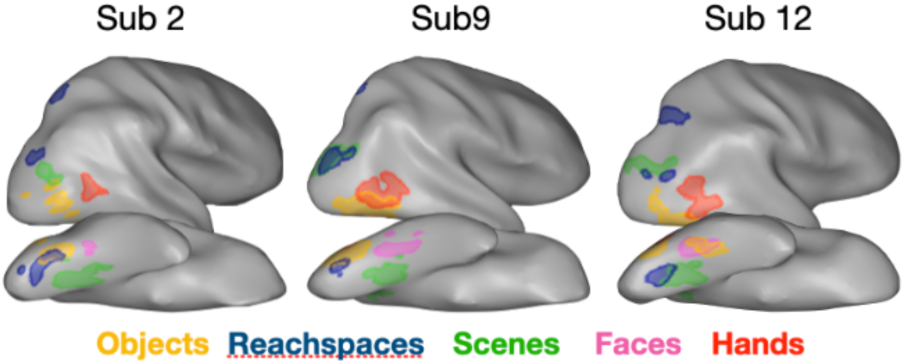
Depiction of the location of the reachspace ROIs in relation to scene-, object-, and face-preferring ROIs, shown in the right hemispheres of three examples subjects.

These activation profiles also illustrate how the stimuli used to a define a region do not allow us to directly infer what specific information is encoded there. For example, scene images depict both spatial layout and multiple objects, but scene ROIs are relatively more sensitive to the spatial layout content of the images. Analogously, reachspace images depict both spatial layout and multiple objects, but reachspaces ROIs are relatively more sensitive to the multi-object content of the images. Thus, the claim here is not that these are “reachspace-selective” regions. Rather, the claim is that these regions are responsive to some content that is relatively more present in naturalistic reachspace images than scene and object images, and we suggest that the presence of multiple individuated objects is likely to be an important factor. Future work will be required to further articulate the distinctive roles of these regions.

### New territory vs. new subdivisions of scene-preferring regions

Finally, we conducted several targeted analyses aimed at understanding whether reachspace-preferring regions are truly separate regions of cortex from scene-and object-preferring ROIs, or whether they are simply new subdivisions. First, we subdivided classically-localized PPA into anterior and posterior regions (e.g. see Bar & Aminoff, 2003; Baldassano et al., 2013), and found that neither subdivision showed a reachspace-preference (**Supplementary Figure 10**). These analyses indicate that the ventral reachspace-preferring ROI does not correspond to this known subdivision of PPA.

Next, we quantified the overlap between all ROIs, given that it was statistically possible for scene-preferring regions (defined with a standard scene>object contrast in localizer runs) to overlap with reachspace-preferring regions (defined through a preference map in experimental runs). However, we found relatively little overlap among the ROIs (e.g. for the ventral ROI, there was a 4.4 ± 1.8% overlap with PPA, 4.6 ± 2.1% with pFs, and 0.1 ± 0.1% with FFA; see **Supplementary Tables 4 and 5** for all overlap results). The relationship among these ROIs is visualized for three individual participants in **Figure 7**, and for all participants in **Supplementary Figure 11**. Overall, reachspace-preferring ROIs largely occupy different regions of cortex than object-, scene-, face-, and hand-selective cortex.

Finally, we examined whether reachspace regions could be an artifact of population mixing. For example, it is possible that the ventral reachspace-preferring region actually reflects an intermixing of object-preferring neurons (similar to nearby pFs), and scene preferring neurons (similar to nearby PPA) whose competing responses to object and scene images average out at the scale of fMRI, creating the appearance of reachspace tuning. If this were the case, then we would expect that the functional profile of the ventral reachspace region could be predicted by a weighted combination of responses in scene- and object-preferring regions. However, this was not evident in the data (**SI; Supplementary Figure 12**): for example, no mixture of pFS and PPA tuning could predict the preference for all four multi-object conditions over both single object conditions. Further, the superior parietal ROI is also informative, as this region shows both a reachspace preference and a functional fingerprint similar to other reachspace-preferring ROIs, but is anatomically far from classic scene-selective regions. Taken together, these analyses provide evidence that reachspace-preferring regions may constitute a distinct processing network from both scene-selective and object-selective networks.

## Discussion

The aim of this study was to characterize how the visual system responds to views of reachable environments relative to views of full-scale scenes and close-up singleton objects. We found that (1), reachable environments activate distinct response topographies from both scenes and objects; (2) regions with reachspace-preferences are present in consistent locations across participants, allowing us to define ROIs in the posterior collateral sulcus, in dorsal occipto-parietal cortex, and the superior parietal lobule; (3) the response topographies of reachspace preferences are maintained in an image set contolling for luminance, contrast, and global spatial frequency; (4) reachspaces elicit dissociable activity in scene and object ROIs, driving these regions to an intermediate degree; (5) reachspace-preferring regions have peripheral biases and (6) have distinctly higher response to the presence of multiple isolated objects over near-scale spatial layout with minimal object content, a combination that is unique among the ROIs explored here; (7) the reachspace-preferring regions do not appear to be a subset of the classic category-selective areas; that is, it is not the case that reachspaces revealed new divisions in regions that have long been discussed in the literature, but rather that they revealed an unexpected pattern of preference in regions adjacent to them.

### Situating reachspace-preferring cortex

Activations across a similar constellation of regions were found when participants attended to the reachability of objects versus their color or location (Bartolo, Coello, Edwards, Delepoulle, Endo & Wing, 2014), and when participants attended to a ball approaching their hand versus a more distant target (Makin, Holmes, & Zohary, 2007). In addition, the three ROIs corresponding to peaks in reachspace preference appear to overlap a subset of the parcels from Fischer, Mikhael, Tenenbaum & Kanwisher (2016), which are involved in making predictions about the physical behavior of objects. Taken together, the correspondence between these results suggest that the ROIs which preferred reachable-scale views may be generally important for reachability judgments, and suggest a potentially broader role in behaviors that rely on accurate predictions regarding objects in the physical world.

The ventral reachspace ROI lies near a swath of cortex sensitive to features of object ensembles (Cant & Xu, 2012), to the texture and surface properties of single objects (Cant, Arnott & Goodale, 2009), to regions that are sensitive to videos of actions being performed in the near space (Tarhan & Konkle, 2020), and near the posterior edge of a color-biased band running along ventral IT cortex (Lafer-Sousa, Conway, Kanwisher, 2016; Conway, 2018). The occipital reachspace ROI lies in the vicinity of inferior parietal regions associated with the maintenance of multiple objects in working memory (Todd & Marois, 2004; Xu & Chun, 2006). And, the superior parietal reachspace ROI falls in the territory thought to contain information about the reachability of an object (Gallivan, Cavina-Pratesi & Culham, 2009; Gallivan, McLean & Culham, 2011), and the type of object-directed hand movement that is planned (Gallivan, McLean, Valyear, Pettypiece & Culham, 2011). Interestingly, this ROI also appears to overlap the posterior locus of the Multiple-Demand Network, a network of fronto-parietal regions associated with the control of visual attention and the sequencing of cognitive operations (Duncan, 2010; Fedorenko, Duncan & Kanwisher, 2013). Future studies with targeted comparisons will be required to map these functions together and assess the degree to which they draw on common representations.

Finally, recently Tarhan & Konkle (2020) found that ventral and dorsal stream responses to videos of people performing actions showed tuning curves that were related to the “interaction envelope” (Bainbridge & Oliva, 2015) of the depicted action, and were sensitive to whether the actions were directed at objects in near space or far space. This result is also broadly consistent with the present results, where the scale of depicted space seems to be an important factor in the structure of responses across the entire visual system (and see Peer et al., 2019 for a related claim extending beyond the visual system).

### Implications for the visual representation of reachable space

The existence of reachspace-preferring cortex suggests that near-scale environments require some distinctive processing relative to navigable-scale scenes and close-scale objects. Part of these differences may relate to differences in scale between the views: perceived depth has been shown to affect activation strength in scene regions (Persichetti & Dilks, 2016; Lescroart, Stansbury, & Gallant, 2015; Henderson, Larson, & Zhu, 2008). However, it is clear that the ROIs which prefer reachspaces to scenes and objects do not do so on the basis of scale *alone*: environments which were near-scale, but contained one or no objects elicited low responses in these regions. Instead, the regions responded strongly to images of multi-objects arrays, suggesting a role for object-related content. Is it possible that, then, these regions are best characterized as “multiple object regions”? How important are the background spatial components, such as the desktops, and texture cues to the perceived depth of the scene? Future work will be needed to characterize the effects of scale, number of objects, and their interactions to clarify the functional contributions of these regions for the perception of reachspaces, scenes, and object views.

Finally, it is possible to extend theoretical frameworks for the large-scale organization of (isolated) object information, and apply them to the large-scale organization of objects, reachspaces, and scene views. For example, some have argued that the visual world is divided into domains linked to behavioral relevance, which are separately arrayed along the cortical sheet (e.g. Mahon & Caramazza, 2012; Conway, 2018). Consistent with this action-based perspective, objects, reachspaces and scenes differ in the kinds of high-level goals and behaviors they afford: objects afford grasping, reachspaces afford the coordinated use of multiple objects, and scenes afford locomotion and navigation. Others have argued that the large-scale organization is more of an emergent property that follows from experienced eccentricity and aggregated differences in mid-level image statistics (e.g. Konkle & Oliva, 2012; Long et al., 2018; Arcaro et al, 2017; Hasson et al., 2004, Conway, 2018). Consistent with this input-based perspective, reachspace images as a class are perceptually distinct from both scene and object images, a distinction which is also evident in the learned representations of deep neural networks (Josephs & Konkle, 2019). An intriguing possibility is that, when interacting in a reachspace, the relevant functional extent of the visual information is at an intermediate visual angle, while for navigable-scale scenes the functional extent of the visual field expands farther into the periphery. In sum, there are both action-based and image feature properties that can jointly motivate a large-scale division of objects, reachspaces, and scenes across the visual system.

## Methods

In-text methods provide details about subject, stimuli and ROI definitions. All other method details are available in the supplement.

### Subjects

Twelve participants were recruited for Experiment 1 and Experiment 2. Two participants overlapped. Experiment 3 was conducted in the same session as Experiment 2. All participants gave informed consent and were compensated for their participation. All procedures were approved by the Harvard University Human Subjects Institutional Review Board.

### Stimuli

All stimuli are available on the Open Science Framework (https://osf.io/g9aj5/). For Experiment 1, we collected views of objects, scenes, and reachable environments, each with 10 images from 6 semantic categories (bar, bathroom, dining room, kitchen, office, art studio), yielding 60 images per scale. These images were divided into 2 equal sets—Image Set A and Image Set B. Object images depicted close-scale views (within 8-12 inches from the object) on their natural background, e.g.: a view of a sponge with a small amount of granite countertop visible beyond it. Reachspace images depicted near-scale environments that were approximately as deep as arm’s reach (3-4ft), and consisted of multiple small objects arrayed on a horizontal surface, e.g.: a knife, cutting board and an onion arrayed on kitchen counter. Scene images depicted views of the interior of rooms, e.g.: a view of a home office.

For Experiment 2, we created a controlled version of Image Set B where all images were grayscaled, matched in average luminance, contrast, and global spatial frequency content using the SHINE toolbox (Willenbockel, et al, 2010).

Experiment 3 included 10 stimulus conditions: (1) reachspaces images with the background removed in photoshop, yielding images of multiple objects in realistic spatial arrangements; (2) reachspaces images with background removed and the remaining objects scrambled, where the objects from the previous condition were moved around the image to disrupt the realistic spatial arrangement; (3) 6 objects with large real-world size, e.g. trampoline, dresser, arranged in a 3×2 grid on a white background; (4) 6 objects with small real world size, e.g. mug, watch, arranged in a 3×2 grid on a white background, and presented at the same visual size as the previous image condition; (5) reachable environments with all objects removed except the support surface; (6) reachspaces containing only a single object on the support surface; (7) vertical reachspaces, where the disposition of objects was vertical rather than horizontal, e.g. shelves, peg-boards; (8) regular reachspaces, i.e. horizontal, as in earlier experiments (9) objects, i.e. close-up views of single objects on their natural background; and (10) scenes, i.e. navigable scale environments. Further details on stimulus selection and controls are available in the supplement.

### Defining ROIs with reachspace preferences

For Experiment 1, Reachspace-preferring ROIs were defined manually in Brain Voyager by applying the conjunction contrast RS>O & RS>S, using four experimental runs with the same image set. We decided a priori to define all reachspace ROIs using Image Set A runs, and extract all activations for further analysis from Image Set B runs. These results are reported in the paper, but we also validated all analyses by reversing which image set was used to localize vs extract activations, and these results are reported in the supplement.

For the ROIs used in Experiments 2 and 3 (run in the same session), we designed an automatic ROI-selection algorithm, guided by the anatomical locations of these regions in Experiment 1. This method allowed for the precise location of ROIs to vary over individuals, while still requiring them to fall within anatomically-constrained zones. The algorithm located the largest patch in the vicinity of the average location of the ROIs from E1 where the univariate preference for reachspaces over the next-most-preferred category exceeded 0.2 beta (more details in supplement). This automated procedure was developed using a separate pilot data set and all parameters were decided a priori (but see Supplementary Figure 8 for a visualization of the consequences of this parameter choice).

## Acknowledgements

We would like to thank Leyla Tarhan, Dan Janini, Ruosi Wang, John Mark Taylor, Grace Edwards and Tatiana Lau for their help scanning participants. This work was carried out at the Harvard Center for Brain Science and is supported by an NIH Shared Instrumentation Grant to the Center for Brain Science (S10OD020039). We acknowledge the University of Minnesota Center for Magnetic Resonance Research for use of the multiband-EPI pulse sequences.

## Author Contributions

E.J. and T.K. designed the research, E.J. performed the research, E.J. and T.K. wrote the paper

## Supplementary Information

### Supplementary Methods

#### Subjects

Twelve participants were recruited each for Experiment 1 and Experiment 2. Experiment 3 was conducted in the same session as Experiment 2, and thus represents the same participants. Two people participated in both E1 and E2, and author EJ participated in E1. Participants were between the ages of 20 and 31, and 13 out of 22 participants were female. One additional person participated in E1, but was excluded prior to analysis for falling asleep in the scanner. All participants gave informed consent and were compensated for their participation. All procedures were approved by the Harvard University Human Subjects Institutional Review Board.

#### Acquisition and Pre-processing

All neuroimaging data were collected at the Harvard Center for Brain Sciences using a 32-channel phased-array head coil with a 3T Siemens Magnetom Prisma fMRI Scanner (Siemens Healthcare, Erlangen, Germany). The Siemens Auto-Align tool was used to ensure reproducible placement of image fields of view. High-resolution anatomical images were collected with a T1-weighted magnetization-prepared rapid gradient multi-echo sequence (multi-echo MPRAGE [1], 176 sagittal slices, TR=2530 ms, TEs=1.69, 3.55, 5.41, and 7.27 ms, TI=1100 ms, flip angle=7°, 1 mm3 voxels, FOV=256 mm, GRAPPA acceleration=2). For functional runs, blood oxygenation level-dependent (BOLD) data were collected via a T2*-weighted echo-planar imaging (EPI) pulse sequence that employed multiband RF pulses and Simultaneous Multi-Slice (SMS) acquisition [2-5]. For the task runs, the EPI parameters were: 69 interleaved axial-oblique slices (25 degrees toward coronal from ACPC alignment), TR=1500 ms, TE=30 ms, flip angle=75°, 2.0 mm isotropic voxels, FOV=208 mm, in-plane acceleration factor (GRAPPA)=2, SMS factor=3). The SMS-EPI acquisition used the CMRR-MB pulse sequence from the University of Minnesota.

Functional data were preprocessed using Brain Voyager QX software with MATLAB scripting. Preprocessing included slice-time correction (ascending trilinear interpolation), 3D motion correction (sinc interpolation), linear trend removal, temporal high-pass filtering (0.0078 Hz cutoff), spatial smoothing (4 mm FWHM kernel), AC-PC alignment and transformation into Talairach (TAL) coordinates. Three dimensional models of each subject’s cortical surface were generated from the high-resolution T1-weighted anatomical scan using the default segmentation procedures in FreeSurfer. For visualizing activations on inflated brains, surfaces were imported into Brain Voyager and inflated using the BV surface module. Gray matter masks were defined in the volume based on the Freesurfer cortex segmentations.

#### E1 and E2 Stimuli

All stimuli are available for download on the Open Science Framework (https://osf.io/g9aj5/). We collected views of objects, scenes, and reachable environments, each with 10 images from 6 semantic categories (bar, bathroom, dining room, kitchen, office, art studio), yielding 60 images per scale. These images were divided into equal 2 sets—Image Set A and Image Set B. Object images depicted close-scale views (within 8-12 inches from the object) on their natural background, e.g.: a view of a sponge with a small amount of granite countertop visible beyond it. Reachspace images depicted near-scale environments that were approximately as deep as arm’s reach (3-4ft), and consisted of multiple small objects arrayed on a horizontal surface, e.g.: a knife, cutting board and onion arrayed on kitchen counter. Scene images depicted views of the interior of rooms, e.g.: a view of a home office.

Additionally, using Image Set B we created a controlled image set, where all images were grayscaled, matched in average luminance, contrast, and global spatial frequency content using the SHINE toolbox (Willenbockel, et al, 2010). Images were spatial frequency-matched using the specMatch function, then luminance-matched using the histMatch function, both with default parameters.

#### Experiment 1 reachspace preference analyses

The main experimental protocol for Experiment 1 consisted of a blocked design with 18 image conditions, depicting three scales of space (object, reachspace, scene views), drawn from six different semantic categories (bar, bathroom, dining room, kitchen, office, studio). Each run contained two blocks per condition, with blocks lasting 6s and consisting of 5 unique images and 1 repeated image. Within a block, each image was presented in isolation on a uniform gray background for 800ms followed by a 200ms blank. There were twelve 10s fixation blocks interleaved throughout the experiment, and each run started and ended with a 10s fixation block. A single run lasted 5.93 min (178 volumes). Participants viewed eight runs of the experimental protocol. Four runs were completed with Image Set A and four with Image Set B (see Stimuli), presented in alternating order over the course of the scan session. Participants’ task was to detect a image repeated back-to-back, which happened once per block.

General linear models (GLMs) were computed using Brain Voyager software. In Experiment 1, for each participant, separate GLMs were fit for runs containing Image Set A and Image Set B, and a third GLM was fit to all experimental runs together (this combined GLM was only used for Experiment 1 preference map analysis). Data were modeled first with 3 condition regressors (object, reachspace, scene), and then again with 18 condition regressors (3 scales of space x 6 semantic category) for the finer-grained analyses by category and the searchlight analysis. The regressors were constructed based on boxcar functions for each condition, convolved with a canonical hemodynamic response function, and were used to fit voxel-wise time course data with percent signal change normalization and correction for serial correlations. The beta parameter estimates from the GLM were used as measures of activation to each condition for all subsequent analyses.

##### Preference Mapping

Group-level preference maps were computed by extracting responses to objects, reachspaces and scenes in each voxel from single-subject GLMs, then averaging over subjects. The preferred condition for each voxel was identified in the group-average, and the degree of preference was computed as the activation differences (in betas) between the most preferred condition and the next-most-preferred condition. Responses were visualized for visually-responsive voxels only, which were defined as those that were active in an All vs Rest contrast at a threshold of t>2.0 in at least 30% of the participants. Early visual regions (V1-V3) were defined by hand on inflated brain, guided by the contrast of horizontal vs. vertical meridians from a retinotopy run (see below for run details). Group average V1-V3 was obtained by generating single subject early visual cortex maps, and selecting voxels that fell within V1-V3 in at least 30% of the participants. These voxels were removed from the visualization. Preference maps were visualized by projecting these voxels’ preferred condition (indicated by color hue) and the degree of preference (indicated by color intensity) onto the cortical surface of a sample participant. For Experiment 1, preference maps were computed from a GLM modeled with data from all 8 experimental runs.

##### RS ROI definition

For Experiment 1, three reachspace-preferring ROIs were defined manually in Brain Voyager by applying the conjunction contrast RS>O & RS>S, using four experimental runs with the same image set. Conjunction contrasts reveal voxels that show both a preference for reachspaces over scenes and reachspace over objects (assigning them the statistical value corresponding to the less robust of those contrasts). We had decided a priori to define all reachspace ROIs using Image Set A runs, and extract all activations for further analysis from Image Set B runs. These results are reported in the paper, but we also validated all analyses by reversing which image set was used to localize vs extract activations, and these results are reported in this supplement.

##### Region-of-Interest Analysis

For ROI-based analyses, univariate activations were obtained by taking the average beta for each condition in each ROI, then averaging over subjects to obtain the group-level activations. Reachspace-preferring ROIs were defined from 4 runs of the experimental protocol, and activation were extracted from the remaining 4 runs. Experiment 1 activations were examined at two levels, with separate GLMs generated for each. At the environment-scale level we examined the activations to 3 conditions: objects, reachspaces and scenes. In each ROI, we tested whether the preferred condition activated the ROI significantly more than the other conditions, using a priori paired one-sided t-tests. We also extracted responses to objects, scenes and reachspaces at the more granular scale-by-category level (18 conditions: 6 semantic categories represented at each of 3 scales). These data were visualized in a bar graph, where the bars are ordered by the strength of the activation.

#### Experiment 2 reachspace preference analyses

The main experimental protocol for Experiment 2 was the same as Experiment 1. Four runs used original images, specifically Image Set A from Experiment 1, and four runs used controlled images, specifically Image Set B with the low level controls described above. Experiment 2 GLMs were computed as above, and the data were modeled with 3 condition regressors (object, reachspace, scene).

##### Preference mapping

Experiment 2 preference mapping used the same procedure as Experiment 1, with the difference that V1-V3 were not removed from the visualization and subsequent quantification, since these regions were not localized in E2. For Experiment 2, separate preference maps were computed from original- and controlled-image runs, each estimate from a GLM with 4 runs.

To quantitively compare the similarity of preference maps elicited by original and controlled images, we assessed the proportion of voxels that showed the same preference across image sets. For the group-level preference map, we first extracted the preferred category and the strength of that preference for each voxel within the group-level mask, for original and controlled images separately (as described above). Next, we extracted the number of voxels that showed the same preference in the two maps (original vs controlled), then divided this by the total number of voxels in the visually-evoked mask, to obtain the proportion of voxels with consistent preference over the image sets. For group-level comparisons, we performed the analyses above on single-subject data, then averaged these values over all subjects.

This method was additionally used to compute the replicability of the original image activations between Experiment 1 and Experiment 2. To do so, the preference map for Experiment 1 was generated from Image Set B only, so that the preference maps being compared were generated from GLM parameter estimates made using the same amount of data (4 runs). Additionally, a common activity mask for the two preference maps was defined by taking the voxels that showed an All-vs-Rest preference of t>2.0 betas in 60% of all of the subject included (i.e. E1 and E2 subjects). Since this analysis was between subjects and no within-subject comparisons were available, the match in preference maps across experiments was only computed at the group-level.

##### RS ROI definition

For Experiment 2, we designed an automatic ROI-selection algorithm, guided by the locations of these regions in Experiment 1. ROIs were defined separately for each participant using the following procedure. First, a spherical proto-ROI was defined around the average central locations of each ROI from E1. The size of the proto-ROI was set to a radius of 6 voxels (18 mm) for the ventral and superior parietal patches, and 9 voxels (27mm) for the occipital patch, to account for different amounts of variation in the expected ROI locations. Then, the reachspace conjunction map with RS>O & RS>S was computed and spatially smoothed (5-mm gaussian kernel, sigma=1). Next, the single voxel with the highest reachspace-preference falling in each proto-ROI was selected and used as the center of 6mm spherical ROI.

Finally, the voxels within this sphere with the most statistically robust preference for reachspaces were retained for the final ROI, using the following procedure. Low-preference voxels were iteratively dropped from the ROI until the region’s univariate preference for reachspaces over the next-most-preferred category exceeded 0.2 beta. This method allowed us to define the largest ROI that still showed a relatively high reachspace bias. This automatic ROI-selection regime was developed in a separate pilot data set before being applied here, and all parameters were decided a priori, but see **Supplementary Figure 13** for visualization of how parameter variation affected significance of the final analysis. Activations within these ROIs were always assessed from independent data sets.

##### Region-of-Interest Analysis

ROI analysis of reachspace regions used the same procedure as Experiment 1, with the exception that this analysis was only performed at the environment scale level (i.e. betas were extracted for objects, reachspaces, and scenes separately, pooling over semantic category). ROIs were defined with the 4 runs depicting original images, and activations were extracted from the 4 controlled-image runs, and compared using a priori paired one-sided t-test.

#### Experiment 1 classic category-selective ROI analysis

Classic category-selective ROIs were defined in Experiment 1 using a standard localizer protocol. Stimuli included images of bodies, faces, hands, objects, multiple objects, scenes, and white noise. Body images showed clothed bodies with the head erased in photoshop, in a variety of poses. Face images were cropped from the chin to the top of the head, and depicted a variety of facial expressions from humans of different ages, races, and genders. Object images showed single objects on a white background. Multi-object images showed four randomly-selected unique objects occupying four quadrants around the center fixation location, presented over a white background. Scene images showed indoor and outdoor images of navigable-scale spaces.

The localizer protocol contained 8 blocks per image condition, with blocks lasting 6s and consisting of 5 unique images and 1 repeated image. All images were presented in isolation on a uniform gray background for 800ms followed by a 200ms blank. There were eight 8s fixation blocks interleaved throughout the experiment, and each run started and ended with an additional 8s fixation block. A single run lasted 6.9 min (208 volumes), and participants viewed four runs of the localizer protocol. Participants’ task was to detect an image repeated back-to-back, which happened once per block.

ROIs were defined using standard contrasts, and ROI activations were extracted from 4 runs of the main experimental protocol using the same univariate approach described above. Activations were extracted separately for Image Set A runs and Image Set B runs. The latter are reported in the paper, as explained above, and the former appear in the supplement. All stats were a priori paired one-sided t-test.

##### Reachspace ROI overlap analysis

In order to quantify whether the reachspace ROIs consistently overlapped any of the classic ROIs, we first divided them into ventral (PPA, pFs, vRS), and lateral-dorsal ROIs (OPA, LO, hand-preferring, opRS, spRS). Next, we assessed the overlap between the RS ROIs in a given division with each of the other ROIs in that division. For each subject, whole-brain masks were created for each ROI in the pair under comparison, and the number of voxels appearing in both masks was extracted. Then, the number of overlapping voxels was divided by the total number of voxels in the reachspace region, to obtain the percentage of the reachspace ROI voxels that overlapped the comparison ROI. With this definition, overlap estimates of 100% indicate that the reachspace-preferring regions fall fully into existing known regions; estimates of 0% indicate complete separation. This was computed separately for each hemisphere, and for RS ROIs created from each image set (Image Set A vs Image Set B).

#### Experiment 2 classic ROI analysis

Scene- and object-selective regions were defined in Experiment 2 from the main experimental protocol runs with the original images: LO and pFs were defined as objects>scenes; PPA, OPA and RSC were defined as scenes>objects. Activations from all regions were extracted from the 4 experimental runs depicting controlled images, and the analysis was otherwise carried out as described in Experiment 1.

#### Original vs Controlled comparisons

Differences in the patterns obtained in ROI responses between original images (Experiment 1) and controlled images (Experiment 2) were assessed in two ways. First, the overall difference in activations between the images sets was assessed by averaging all the activations (object, reachspace, and scenes) within an ROI to obtain it’s mean response. This was compared between the experiments using between-subject ttest. Statistical threshold were set using Bonferroni corrections, where the number of comparisons was taken as the number of ROIs of each type (i.e. reachspace preferring, object-preferring, and scene-preferring). Second, we examined whether the magnitude of the differences between conditions was different for the two image sets. For this, we calculated the difference between reachspaces and scenes for a given ROI across all subjects, then averaged over subjects. The size of this difference was then compared between original and controlled images. The same was then performed for objects. Comparisons used between-subject t-tests, and statistical threshold were set using Bonferroni corrections, where the number of comparisons was taken as the number of ROIs of each type multiplied by 2 (since there are two comparisons per ROI: RS vs S and RS vs O).

#### Experiment 1 and Experiment 2 PPA subdivision

For Experiment 1 and 2, we additionally subdivided PPA. For each subject, we separately split the PPA from the left and right hemisphere at the midpoint along the anterior to posterior axis. Anterior and posterior PPA were then submitted to the same ROI analysis described above. Statistical tests were completed using Bonferroni-corrected paired one-sided t-test with alpha 0.0125 (i.e. 0.05/4, reflecting the four comparisons being performed in each image set).

#### Eccentricity profile analysis

Data for the eccentricity analysis were collected in the same run as Experiment 1, and thus represent the same subjects and reachspace ROIs. The retinotopy protocol consisted of 4 stimulus conditions: horizontal bands, vertical bands, central stimulation, and peripheral stimulation. Vertical and horizontal bands (subtending ∼1.7° × 22° and ∼22° × 1.7° respectively) showed checkerboards which cycled between states of black-and white, white-and-black, and randomly colored at 6hz. Central and peripheral rings (radius ∼1.2° to 2.4° and radius ∼9.3° to the edges of the screen, respectively) showed photograph fragments which cycled between patterns of object ensembles (e.g. beads, berries, buttons) and scene fragments (c.f. Cant & Xu, 2012; Zeidman, Silson, Schwarzkopf, Baker & Penny, 2018). Each run contained 5 blocks per condition, with blocks lasting 12 seconds. There were four 12s fixation blocks interleaved throughout the experiment, and each run started and ended with an additional 12s fixation block. Each run lasted 4.4 min (162 volumes), and participants viewed two runs of the retinotopy protocol. Participants’ task was to maintain fixation and press a button when the fixation dot turned red, which happened at a random time once per block.

##### ROI analysis

We explored the eccentricity preference of object, reachspace, and scene ROIs (defined as described for Experiment 1 above), and for ROIs corresponding to scenes- and object ROIs. Average betas were extracted for two eccentricity conditions: central stimulation, and peripheral stimulation. Activations in the two conditions were compared using a paired one sided t-test with a Bonferroni corrected alpha value of 0.0125 (i.e. 0.05/8, reflecting the 8 ROIs where we tested for a difference between these conditions).

#### Post-hoc fingerprint profile analysis

To examine the broader tuning of object, scene and reachspace ROIs, we performed a post-hoc analysis, using activations extracted from the Experiment 1 localizer. The localizer runs included bodies, faces, hands, objects, multiple objects, scenes, and white noise). We extracted responses in Experiment 1 reachspace-preferring ROIs to these 8 conditions for each subject, and averaged the activations over subjects. First, we visualized these responses in a polar plot. Next, we noted what the most preferred condition was, and tested whether this was significantly different than the next-most preferred condition using one-tailed pair t-tests with Bonferroni-corrected alpha 0.0167 (0.05/3, reflecting the 3 reachspace-preferring ROIs where we tested for this difference).

We also examined activations to these conditions in object- and scene-processing cortex. Since the localizer runs were used to extract activations, we couldn’t use them to define ROIs. Instead, ROIs were defined as a spherical ROI with a 9-mm radius centered on the typical anatomical location of each region based on an internal meta-analysis (for left/right hemisphere, ROI centers were as follows: PPA: -25 -41 -6/ 25 -42 -7; OPA: -25 -76 25/ 28 -81 26; RSC: -16 -51 9/ 18 -49 8; LO: -39 -71 -4/ 41 -68 -4; pFs: -38 -53 -13/ 38 -50 -14).The difference between the preferred and next-most-preferred condition was assess using one-tailed pair t-tests with Bonferroni-corrected alpha 0.025, and 0.0167 respectively for areas corresponding to the anatomical location of object and scenes ROIs.

#### Experiment 3 fingerprint profile analysis

Experiment 3 stimuli contained 10 conditions intended to further probe the response profile of the reachspace regions. The conditions were the following: 1) reachspaces images with the background removed in photoshop, yielding images of multiple objects in realistic spatial arrangements; 2) reachspaces images with background removed and the remaining objects scrambled, where the objects from the previous condition were moved around the image to disrupt the realistic spatial arrangement; 3) 6 objects with large real-world size (e.g. trampoline, dresser) arranged in a 3×2 grid one a white background; 4) 6 objects with small real world size (e.g. mug, watch) arranged in a 3×2 grid a white background (presented at the same visual size as the previous image condition); 5) reachable environments with all objects removed except the support surface; 6) reachspaces containing only a single object on the support surface; 7) vertical reachspaces, where the disposition of objects was vertical rather than horizontal (e.g. shelves, peg-boards); 8) regular (i.e. horizontal) reachspaces; 9) objects (i.e. close-up views of single objects on their natural background); and 10) scenes (i.e. navigable scale environments).

Images from conditions 1 and 2 above (reachspace with background removed, and reachspace with background removed and remaining objects scrambled) were generated from the same original images. First, condition 1 images were generated by selecting high-quality reachspace images, and erasing all image content except the 6 salient objects which conveyed the identity and layout of the space. Then, condition 2 images were generated by scrambling the arrangement of the 6 remining objects in the image, and occasionally rotating objects, to break the sense of spatial congruity among them. We ensured that the average placement of objects across all the images (i.e. the heatmap of object locations) was equivalent between condition 1 and condition 2. Images in conditions 5, 6 and 9 (empty reachspaces, reachspaces with single objects, and close up view of single objects) were taken by the experimenter, and represented the same environments. Specifically, a suitable reachspace was selected by the experimenter and cleared of all objects for condition 5, and an images was taken with a camera on a tripod. Then a single salient object was placed in the center of the reachspace for condition 6, at which point a second picture was taken without moving the tripod. Finally, condition 9, the singleton object view, was generated by closely cropping the condition 6 image in Photoshop. Images in conditions 3 and 4 (large and small objects respectively) were programmatically generated by randomly 6 objects drawing from a database of large and small objects, and placing them in 3-across by 2-down grid. Images for condition 7 (vertical reachspaces) were selecting by finding reachable environments where the spatial layout of the objects was primarily on a vertical, rather than horizontal plane. This ranged from spaces with no horizontal extent (e.g. pegboard organization) to spaces with minimal horizontal extent (e.g. shelves). Finally, condition 8 and 10 images (regular reachspaces and scenes) were selected according to the same criteria as E1.

The main experimental protocol for Experiment 3 consisted of a blocked design with the 10 image conditions described above. Each run contained 4 blocks per condition, with blocks lasting 8s and consisting of 7 unique images and 1 repeated image. Within a block, each image was presented in isolation on a uniform gray background for 800ms followed by a 200ms blank. There were eight 10s fixation blocks interleaved throughout the experiment, and each run started and ended with a 10s fixation block. A single run lasted 7 min (210 volumes). Participants viewed four runs of the experimental protocol. Participants’ task was to detect a image repeated back-to-back, which happened once per block.

##### ROI definition

Experiment 3 used the same subjects and ROIs as Experiment 2.

##### Analysis

Responses across the 10 conditions were extracted from all object, scene, and reachspace ROIs. These responses were first visualized in a fingerprint profile. Next, to assess whether object, scene and reachspace ROIs had significantly different response profiles, we performed an analysis of variance to compare ROI types. Responses across the 10 conditions were averaged for all reachspace ROIs, scene ROIs and objects ROIs. These three response profiles were then submitted to a 2-way, condition-by-ROI type ANOVA.

## Supplementary Figures

**Supplementary Figure 1:**
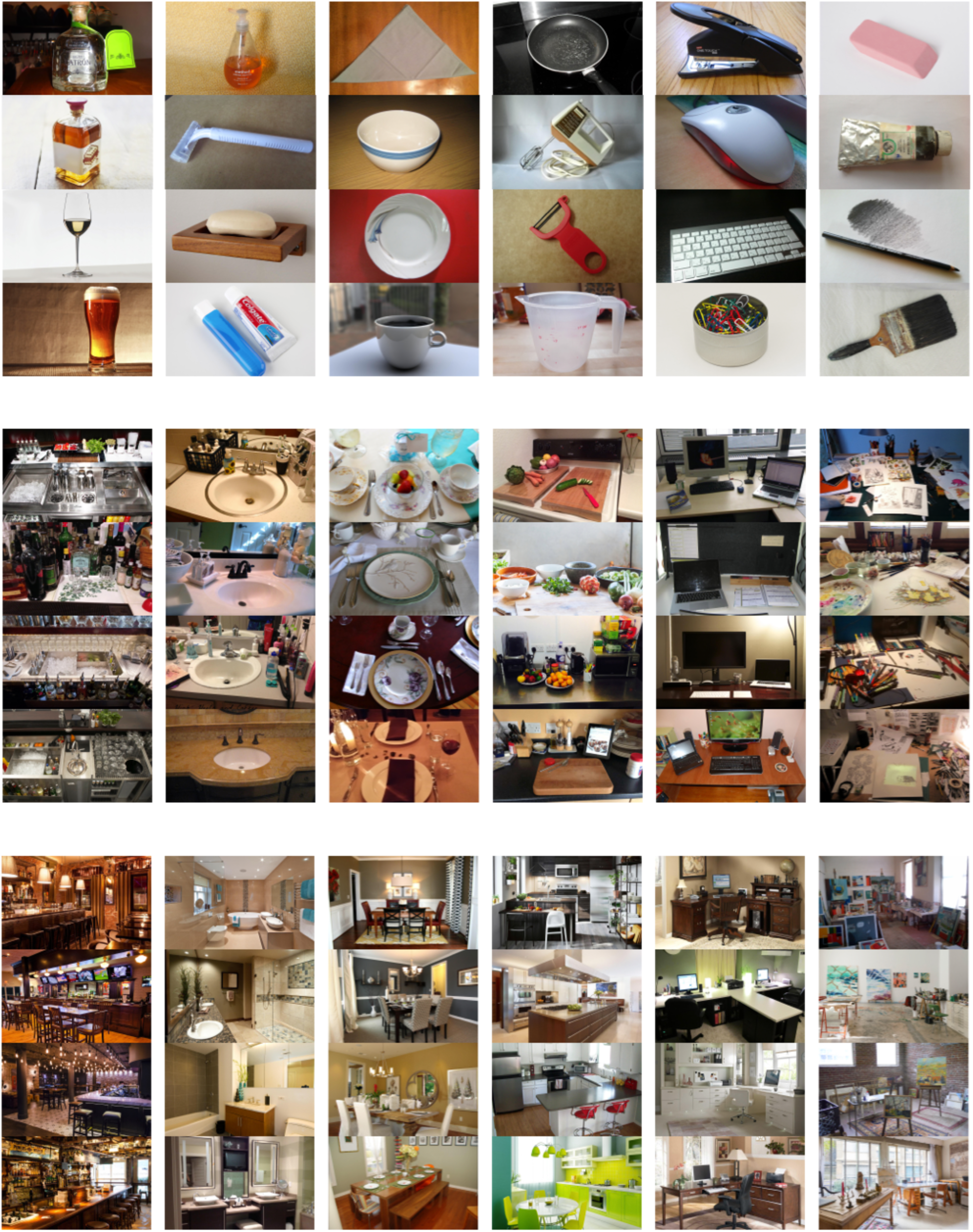
Additional examples of the object (top), reachspace (middle), and scenes (bottom) stimuli used in Experiment 1

**Supplementary Figure 2:**
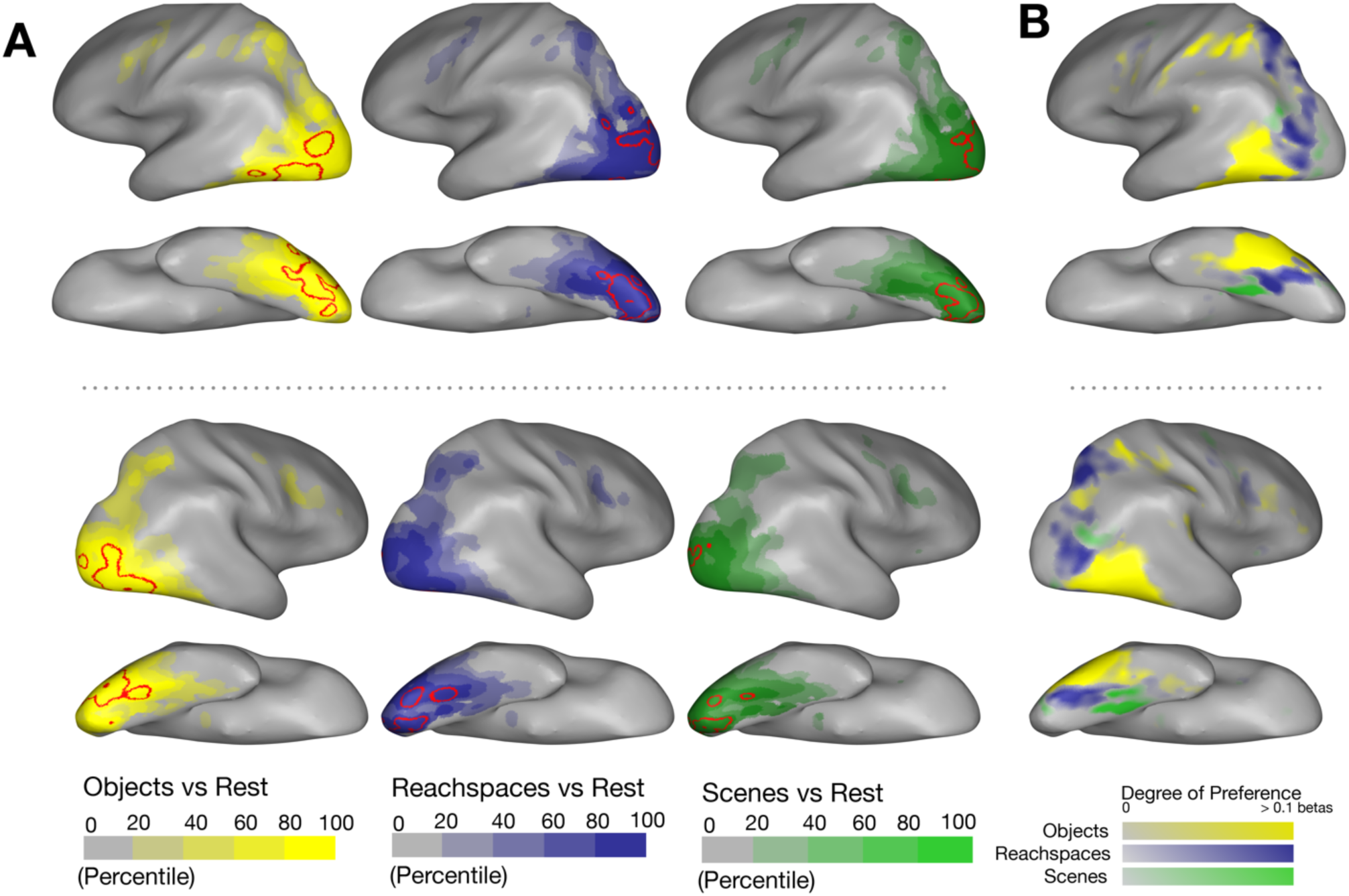
(A) Overall activations are plotted for each stimulus condition (e.g. objects>rest), for group-level data from Experiment 1. Voxels are colored based on the percentile of activation, where voxels in the 95^th^ percentile are outlined in red. Voxels were included if they exceeded T>2.0 for the contrast of all conditions > rest. Note that these activation percentile can reflect not only the strength of the response, but also differences in signal to noise (e.g. different parts of cortex are closer or farther to the coils and ear-canal artifacts), and should be interpreted with this in mind. (B) The group-level preference map is replotted (data from Figure 1) for comparison. This visualization highlights the fact that the strongest overall activation (e.g. top 5% activations) is in partial but not in perfect correspondence with the regions where objects, reachspaces, and scene images show systematically higher responses.

**Supplementary Figure 3:**
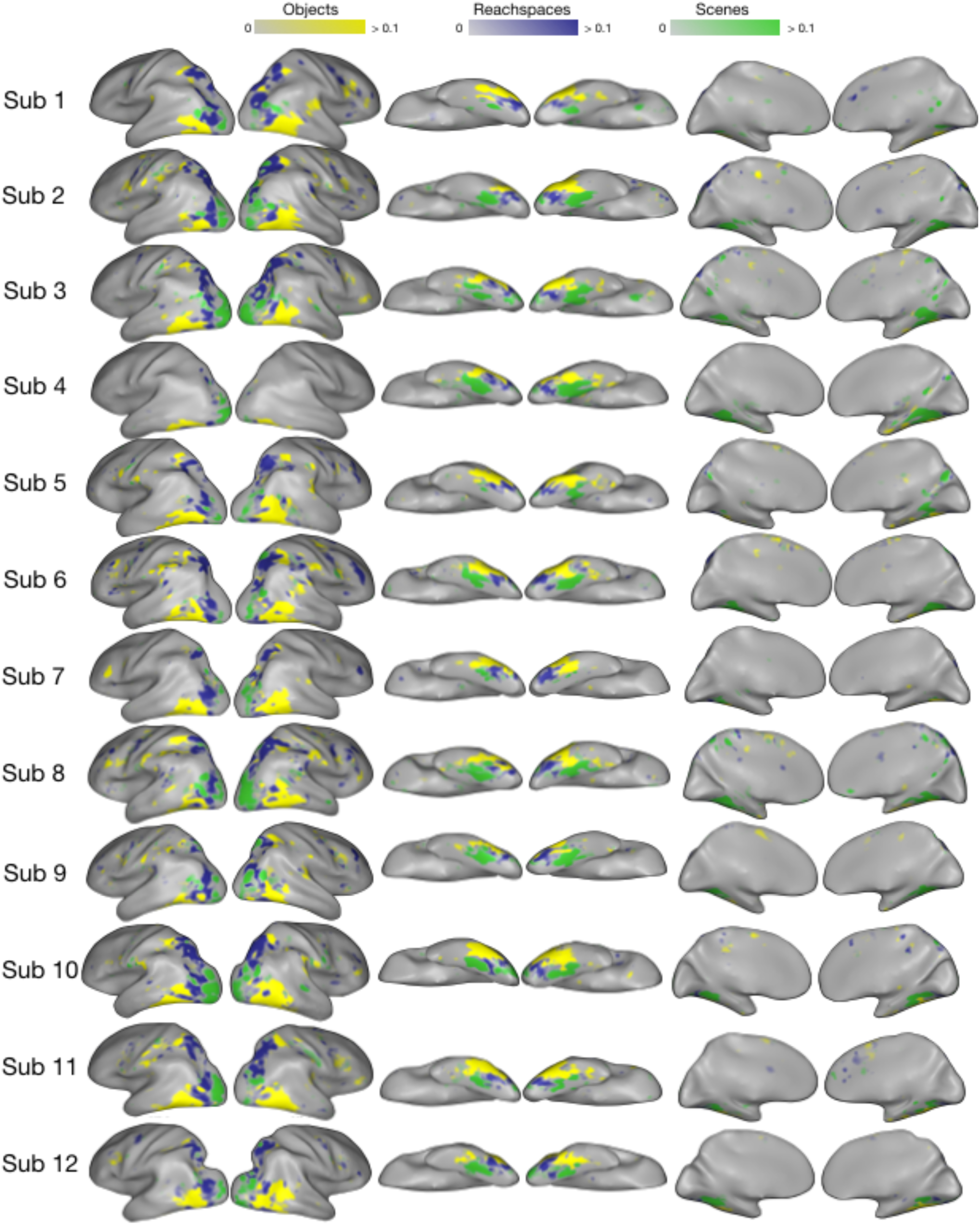
Single-subject preference maps from Experiment 1.

**Supplementary Figure 4:**
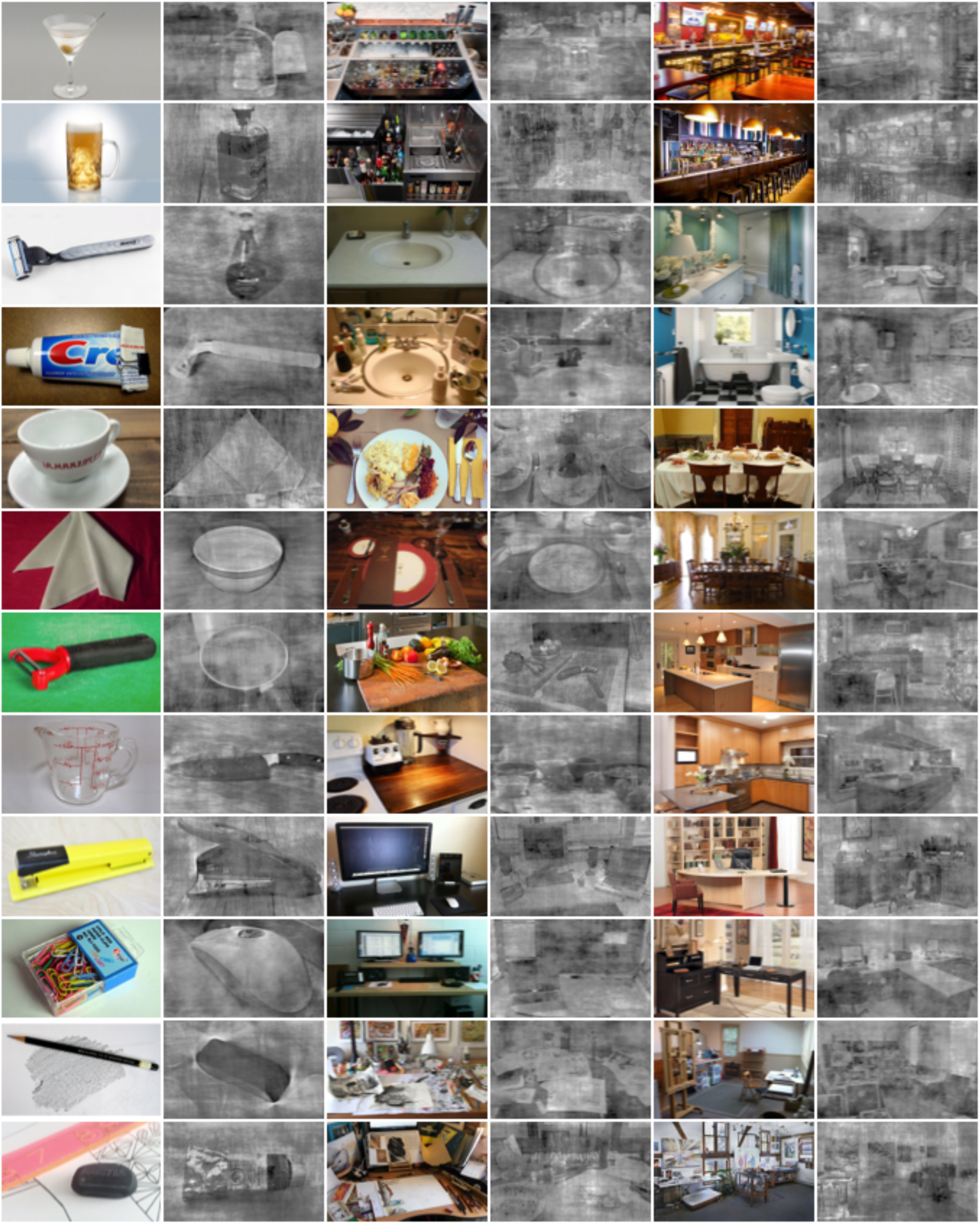
Additional examples of stimuli for Experiment 2.

**Supplemental Figure 5:**
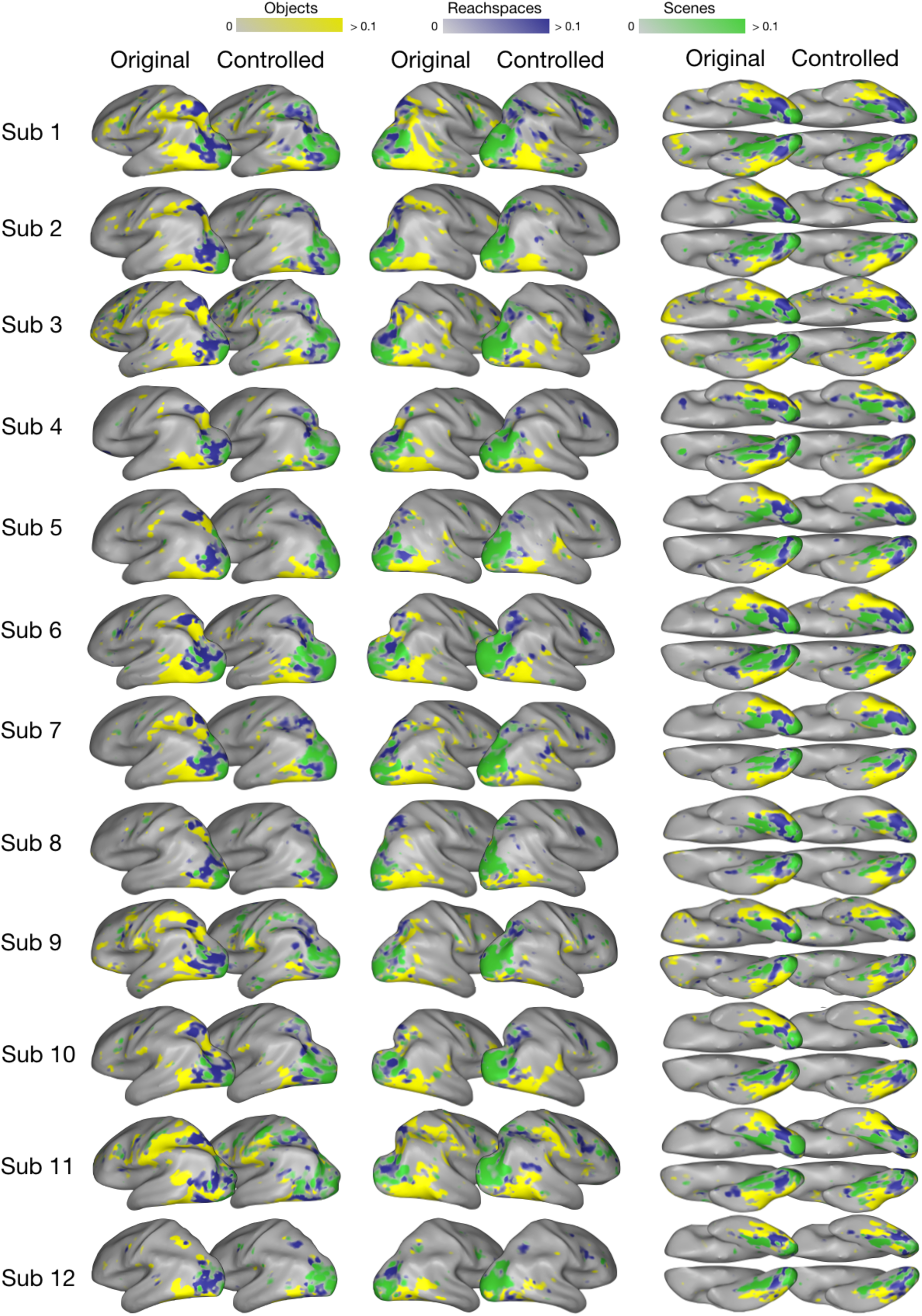
Single-subject preference maps from Experiment 2, obtained from original and controlled images (same color scale used for both original and controlled)

**Supplemental Figure 6:**
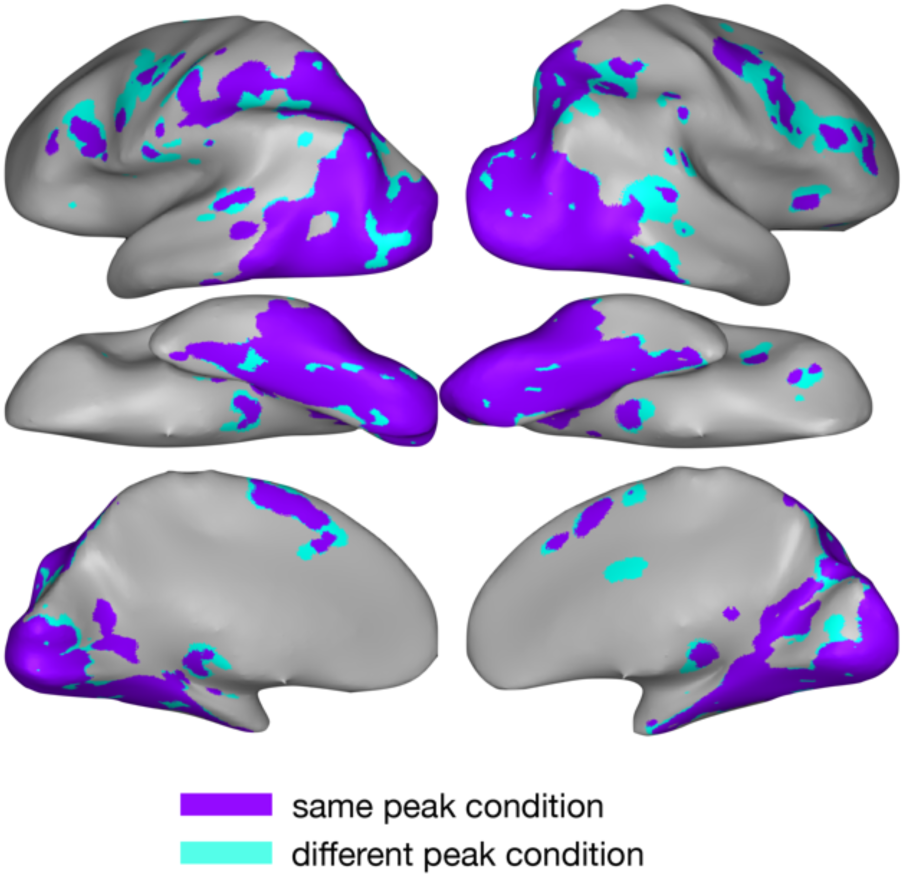
Comparison between original and controlled group-level preference maps from Experiment 2. Cortex colored in purple showed the same preference for objects, reachspaces, or scene images in both Original and Controlled image sets, while cortex colored in cyan had different peak conditions.

**Supplementary Figure 7.**
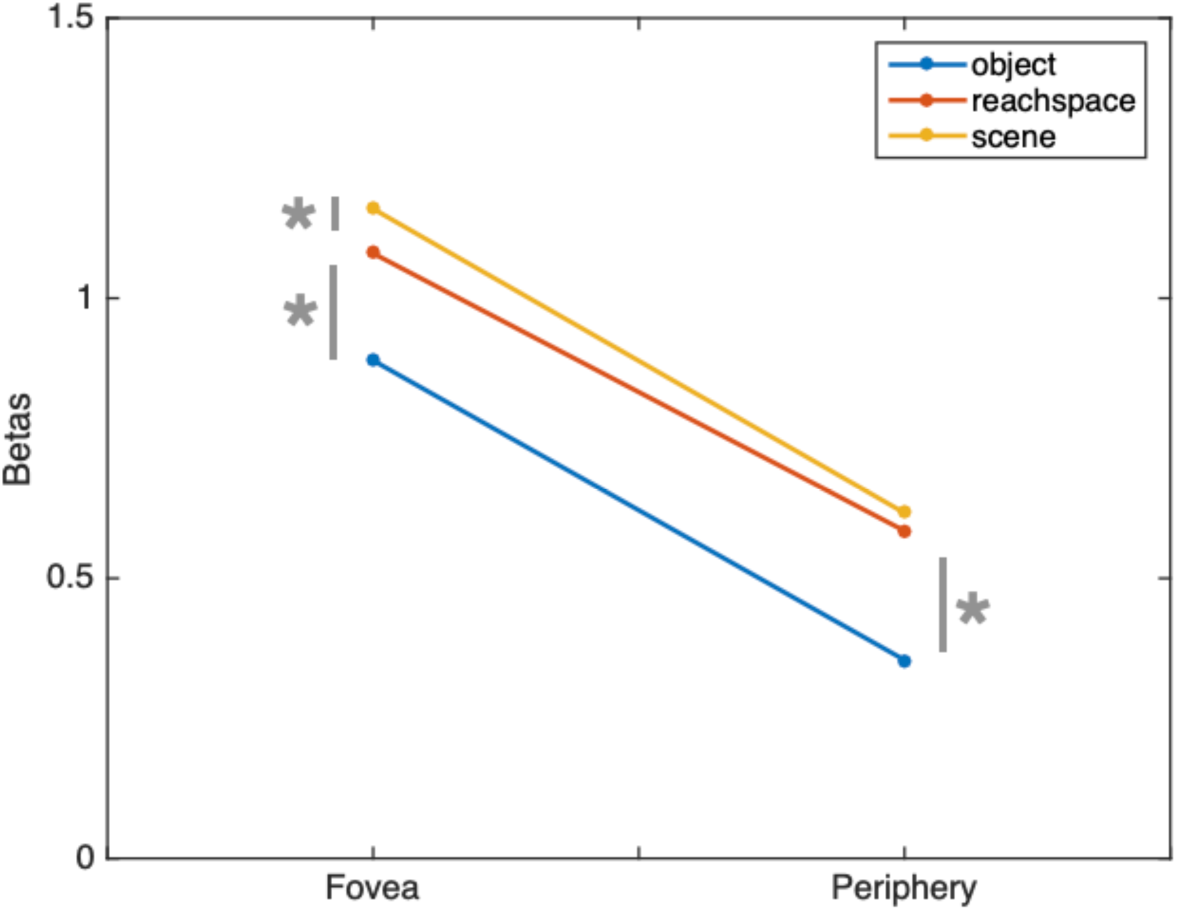
Responses to objects, reachspaces and scenes in foveal and peripheral regions of early visual cortex (V1-V3). Early visual cortex (EVC) was defined using vertical and horizontal meridians from Experiment 1 eccentricity mapping runs, and then divided it into foveal-preferring and peripheral-preferring regions, based on contrasting the Central vs Peripheral conditions. The overall response (average beta) to objects, reachspaces, and scene images from Experiment 1 was computed. The y-axis plots overall response for both foveal and peripheral regions (x-axis), for the three different stimulus conditions. Post-hoc paired t-tests indicated that in foveal cortex, scenes elicited the most activity, followed by reachspaces images and then object objects. In peripheral cortex, there was no statistically significant difference between scenes and reachspaces, but both these conditions showed relatively greater activation than object images.

**Supplementary Figure 8.**
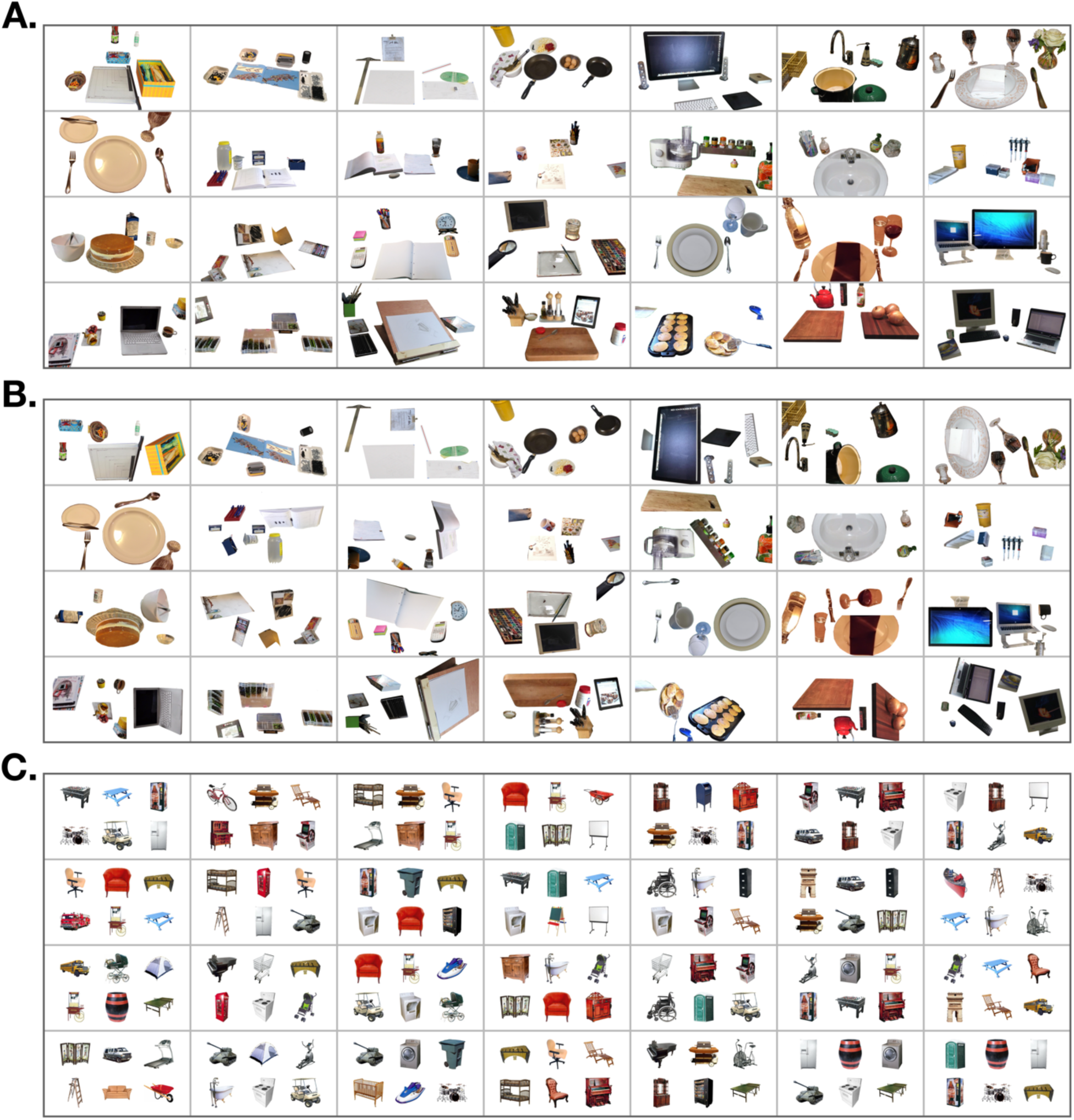
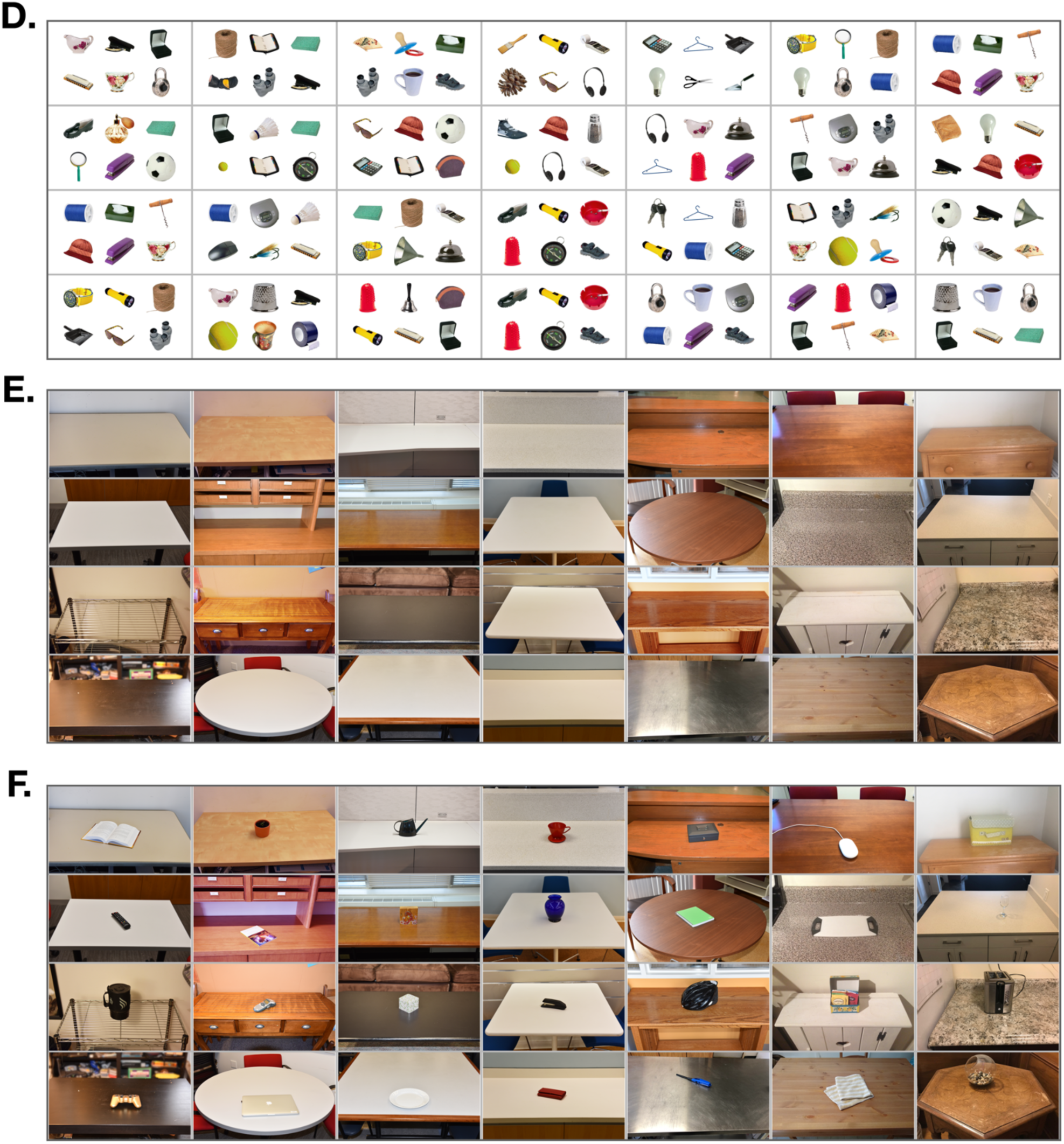
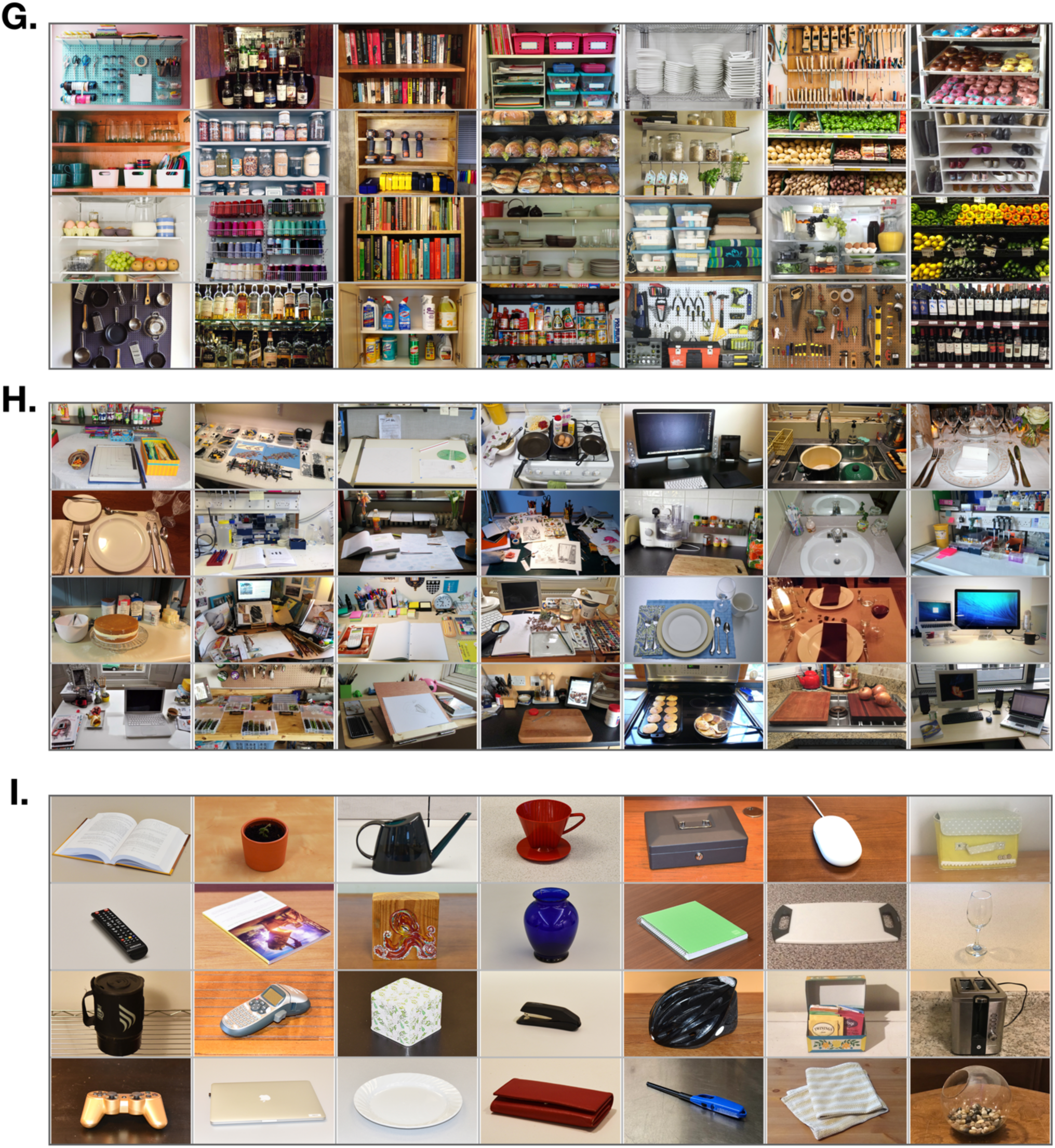
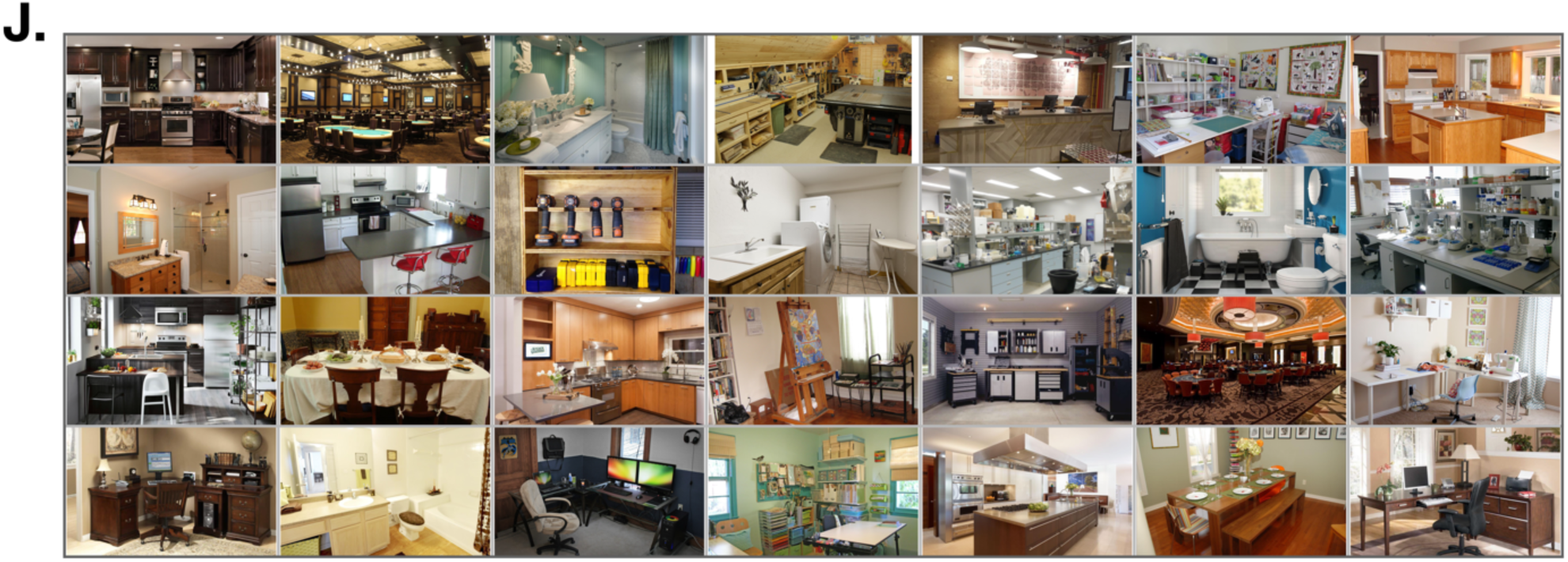
Experiment 3 stimuli. A) reachspaces images with the background removed in Photoshop, yielding images of multiple objects in realistic spatial arrangements; B) reachspaces images with background removed and the remaining objects scrambled to disrupt their spatial arrangement; C) 6 objects with large real-world size (e.g. trampoline, dresser) arranged in a 3×2 grid one a white background. D) 6 objects with small real world size (e.g. mug, watch) arranged in a 3×2 grid a white background (presented at the same visual size as the large object condition); E) reachable environments with all objects removed except the support surface; F) reachspaces containing only a single object on the support surface. G) vertical reachspaces, where the disposition of objects was vertical rather than horizontal (e.g. shelves, peg-boards); H) regular (i.e. horizontal) reachspaces; I) objects (i.e. close-up views of single objects on their natural background). J) Scene images.

**Supplementary Figure 9.**
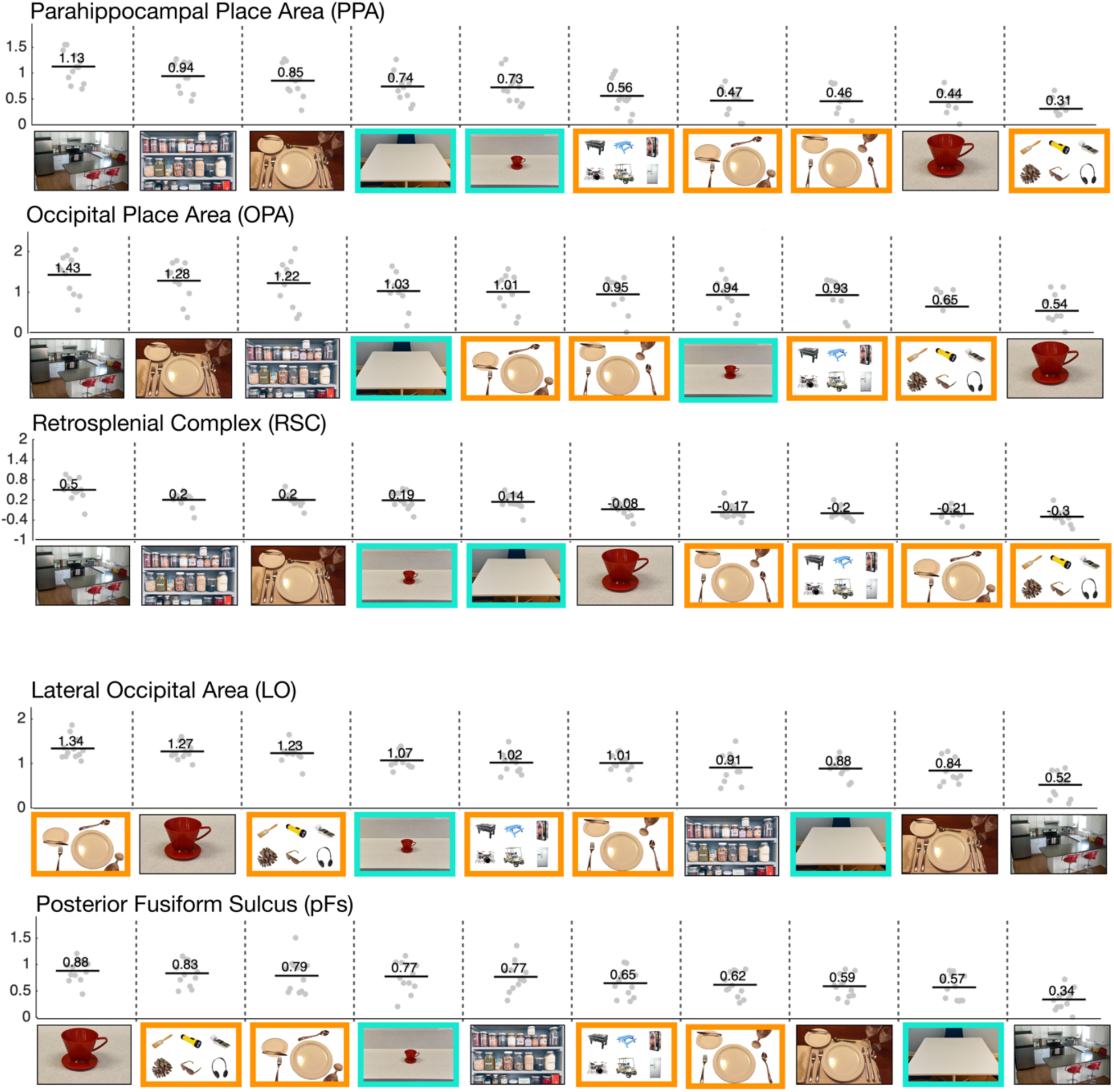
Experiment 3 results for scene- and object-selective ROIs. Responses in scene and object-preferring ROIs across all stimulus conditions are shown, with conditions plotted in order from highest to lowest activations. Images with orange borders indicate stimuli dominated by multiple objects, and images with teal borders highlight images of reachable space with low object content. The mean activation is indicated with a black horizontal bar; gray points indicate single participant data.

**Supplementary Figure 10:**
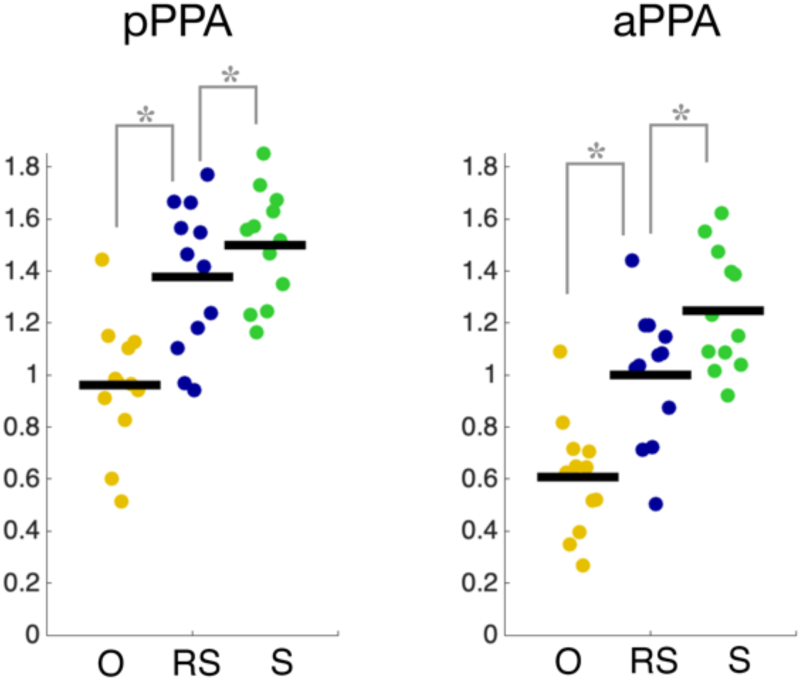
Responses to objects, reachspaces and scenes in PPA subdivisions. In each subject, PPA was divided in half along its anterior-posterior axis, and responses were assessed in each half. While overall activations were higher in posterior PPA (pPPA) than anterior PPA (aPPA), both showed the same pattern of results: scenes elicited greater activation than reachspaces (aPPA: t(11) = 6.22, p<0.01 0.00; pPPA: t(11) = 2.49, p = 0.015), and reachspaces elicited greater activation than objects (aPPA: t(11) = 11.51, p < 0.01; pPPA: t(11) = 9.71, p <0.01),

**Supplementary Figure 11.**
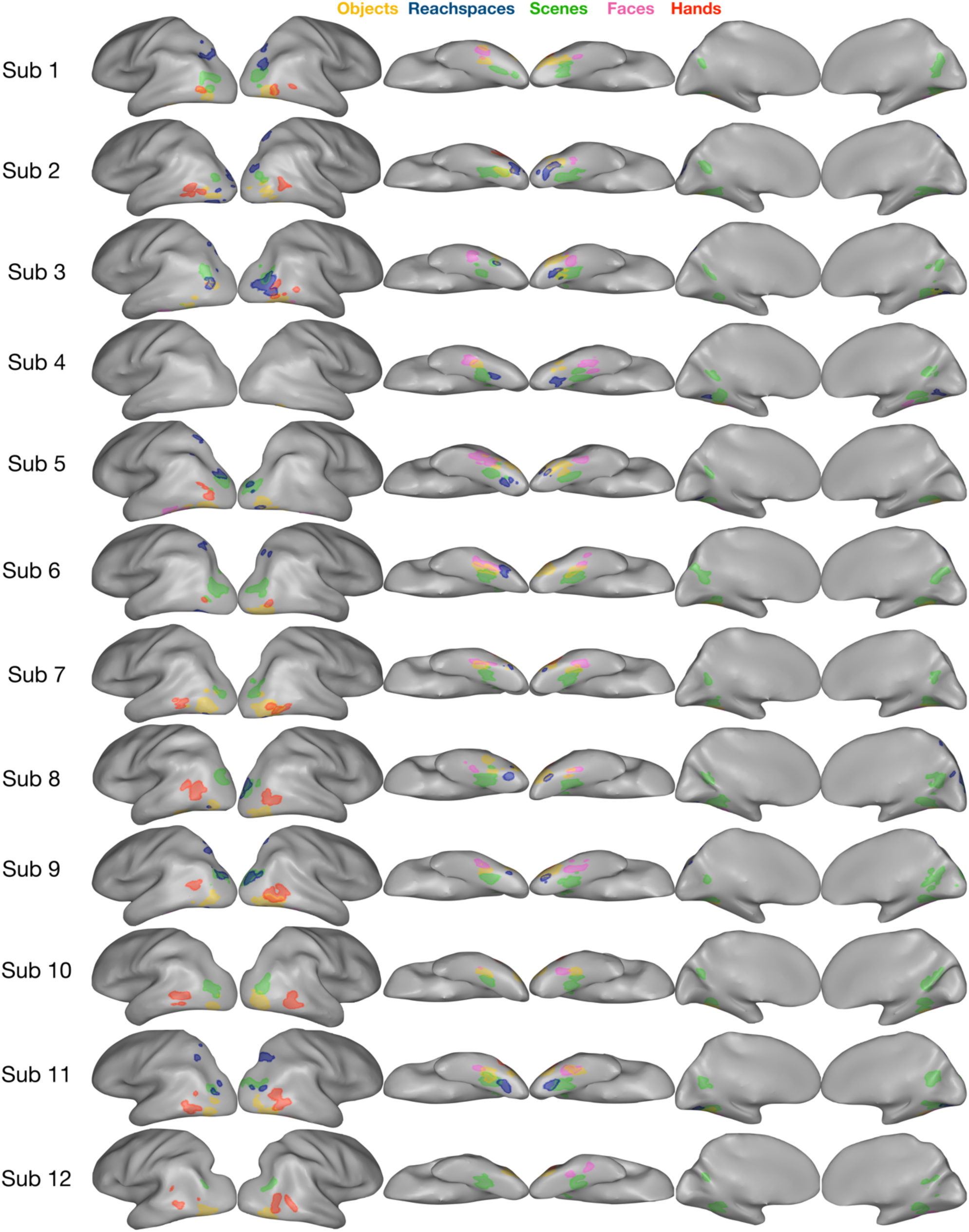
All Experiment 1 ROIs viewed in single-subjects. The color legend is as follows: yellow for objects, blue for reachspaces, green for scenes, pink for faces, orange for hands.

### Supplementary Analysis: Population Mixing

One potential concern is that a voxel’s preferences for reachspace might reflect the mixing of scene and object neurons, rather than the outright existence of any reachspace preferring neurons. For example, it is possible that the observed cluster with a preference for reachspaces on the ventral surface (vROI) may be an artifact of some combination of scene and object responses from nearby PPA and pFs. In this scenario, the voxels’ response to scene images would be artificially depressed (reflecting the average between the PPA-like high response and the pFS-like low response) as would the response to object images (reflecting the pFS-like high response and the PPA-like low response). Response to reachspaces, reflecting the average of the intermediate reachspace response in individual neurons, might remain unchanged. In this way, voxels may show the highest response to reachspace images, even if the individual neural populations within it all responded to reachspaces in an intermediate way.

To examine this possibility, we attempted to predict the 10-condition response profile in the ventral reachspace region using different weighted combinations of PPA and pFs response profile (10%-90%, 20%-80%, 30%-70%, 40-60%, 50%-50%, 60%-40%, 70%-30%, 80%-20-%, and 90%-10%). Predicted and actual responses were compared using Peason’s correlations. The results of this analysis are shown below in **Supplementary Figure 11**.

With this measure, the PPA-pFs weighting that most predicted vROI responses was an even 50-50 split, with r = 0.42 (all correlations below). For comparison, PPA and OPA correlated at r = 0.88, PPA and RSC correlated at r = 0.96, while LO and PFS correlated at r = 0.90. Thus, the correlation between predicted and actual results, for even the most optimal PPA-pFs combination, was half as strong as the correlation to between ROIs with similar category preference.

Importantly, even the best-fitting PPA-pFs weighting (50%-50%) could not predict several crucial aspects of the vROI’s response across the 10 conditions and Experiment 3. First, vROI responded more strongly to all four multi-object conditions (orange bars: multiple large objects in an array, multiple small objects in an array, objects cut from a reachspace image, objects cut from a reachspace image then moved around) than to a close-up view of a single object. The PPA-pFs weighting predicted the opposite. Second, the actual responses for these four conditions were higher than responses to an empty reachspace (orange bars higher than first cyan bar), while the best weighted combination predicted that they should be similar. Finally, responses to these four conditions were higher than responses to a reachspace with only a single object (orange bars higher than the second cyan bar), while the average would have predicted the reverse. Thus, some of the signature elements of the vROI response were not captured by combining PPA and pFs responses.

Finally, to confirm that these conclusions were not an artifact of the measurement that we used, we repeated the analysis using Spearman’s rank ordered correlation. Here we found that the optimal combination was 90% -10%, as well 80%-20%, for PPA and pFs respectively, with r =0.48. For comparison, PPA and OPA correlated at r = 0.88, PPA and RSC correlated at r = 0.88, while LO and PFS correlated at r = 0.92. Altogether, these analyses provide evidence against the possibility that the reachspace preference in vROI voxels simply reflected some combination of scene and object neurons.

**Supplementary Figure 12.**
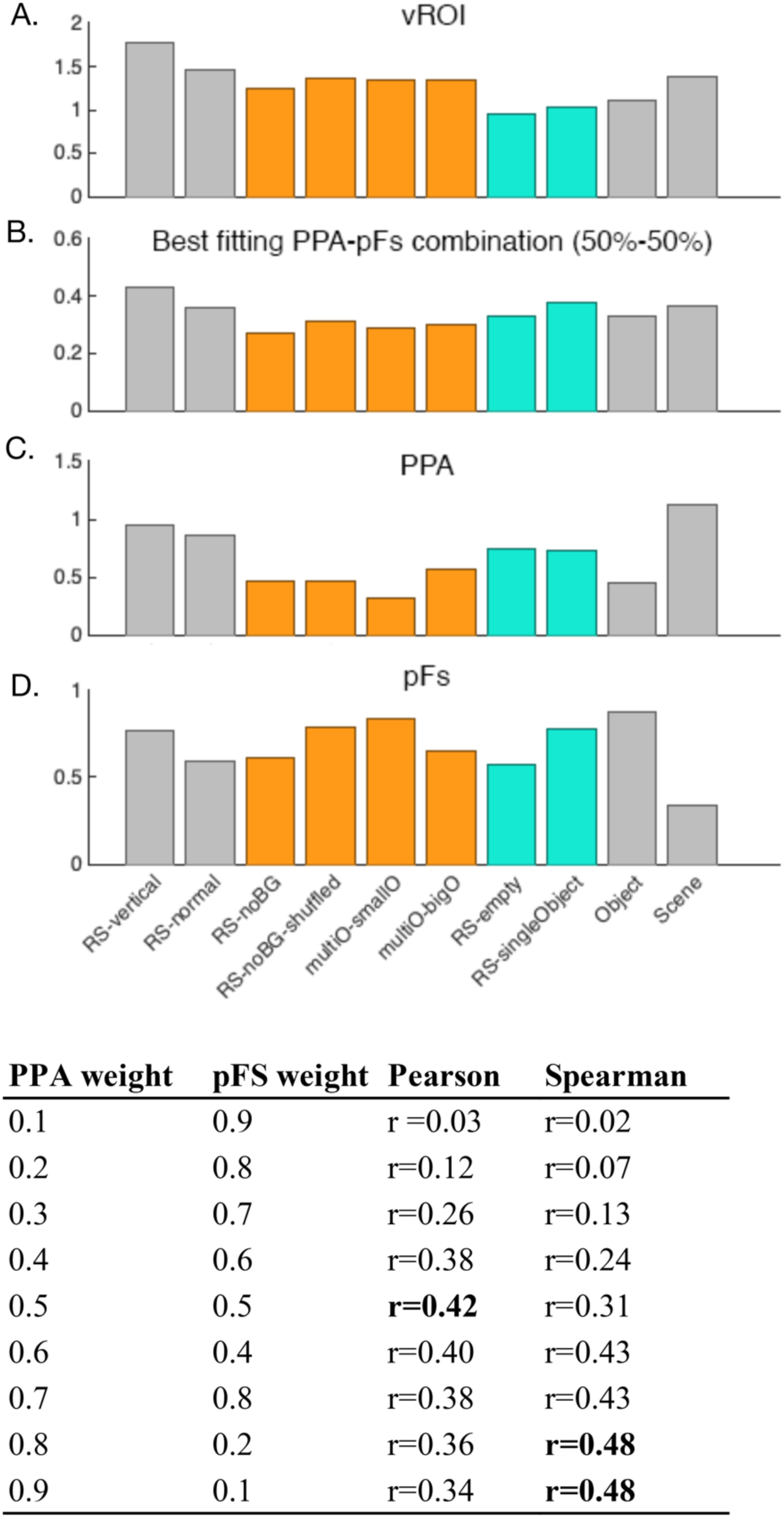
(A): The actual 10-condition response profile of the ventral reachspace-preferring ROI is shown (y-axis: betas; x-axis, conditions). (B) The best fitting PPA-pFS profile combination is shown. (C,D). The profiles for PPA and pFS are also shown. Table shows Pearson and Spearman correlations for all tested weightings of PPA and pFs with vROI.

**Supplemental Figure 13:**
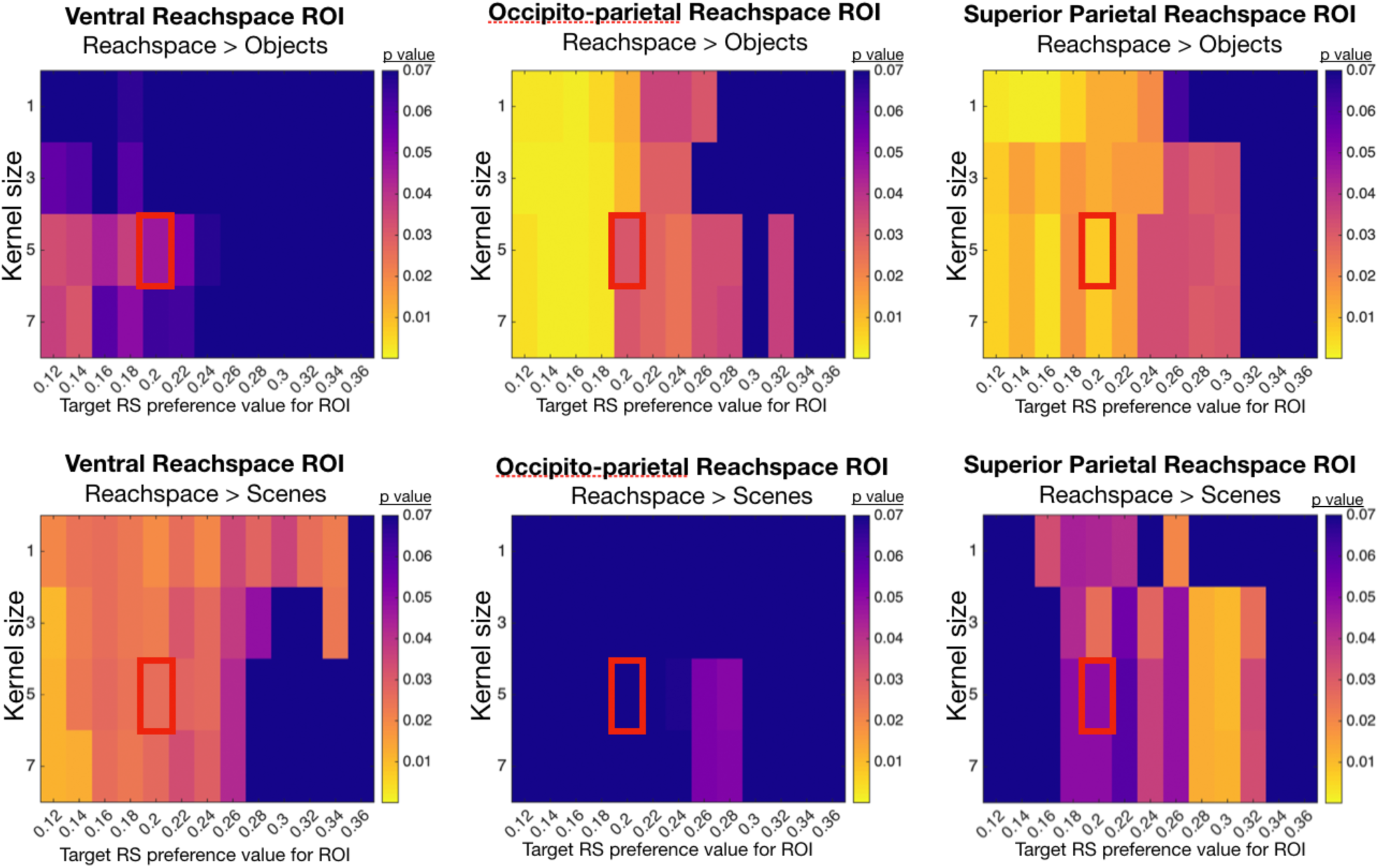
Visualization of how Experiment 1 results vary as the automatic ROI-selection parameters are varied. For Experiment 2, reachspace-preferring ROIs were selecting using a semi-automatic procedure (see Methods). Parameters such as the size of the smoothing kernel and the reachspace-preference threshold value were determined a priori, based on analyses run in a separate set of data. Results in the main text were extracted from ROIs defined using a 5-voxel smoothing kernel and requiring a reachspace preference of 0.2 betas. Here, we display how the statistics significance of the preference for reachspaces over objects (top row) and scenes (bottom row) would have changed with different parameters. In each graph, the rows vary the size of the smoothing kernels applied to the statistical maps computed from the conjunction contrast RS>O & RS>S, from 1 to 7 voxels. The columns vary the threshold for the beta value of reachspace preference we required of the final ROI, from 0.12 to 0.36. The color in each cell shows the statistical significance of the comparison indicated in the title. The red square shows the cell corresponding to the parameters used in the main text.

### Supplementary Tables

**Supplementary Table 1:**
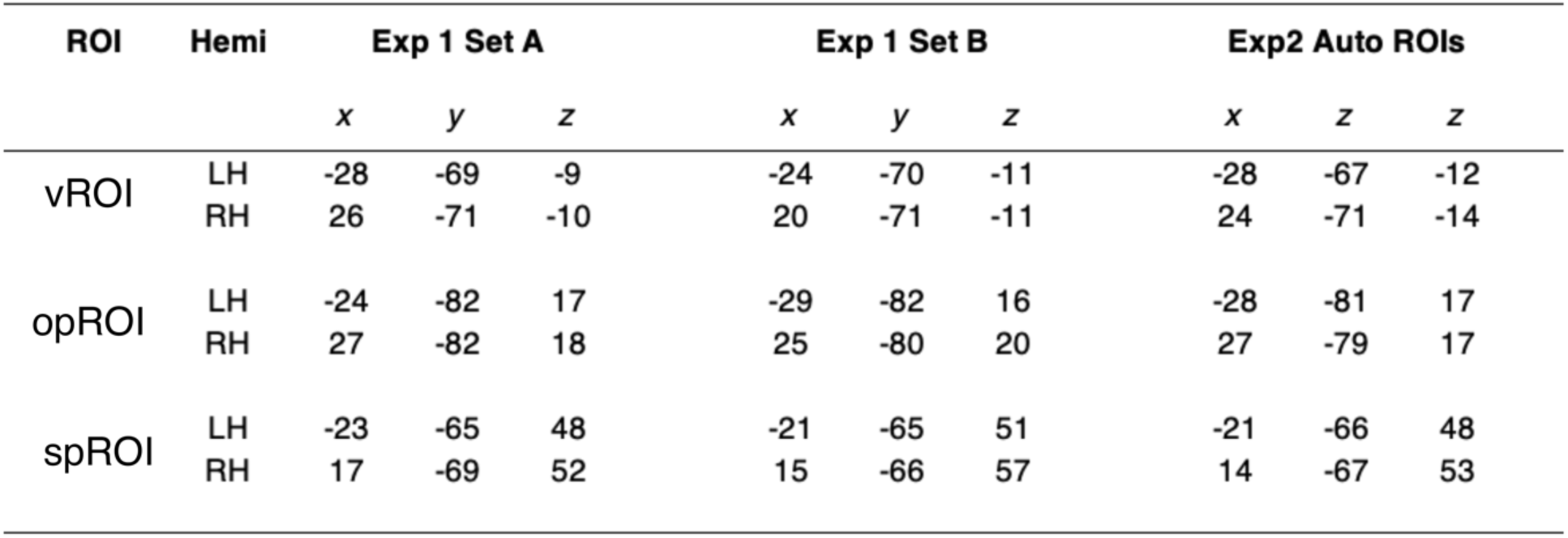
Average TAL coordinates for reachspace-preferring ROIs in all experiments.

**Supplementary Table 2:**
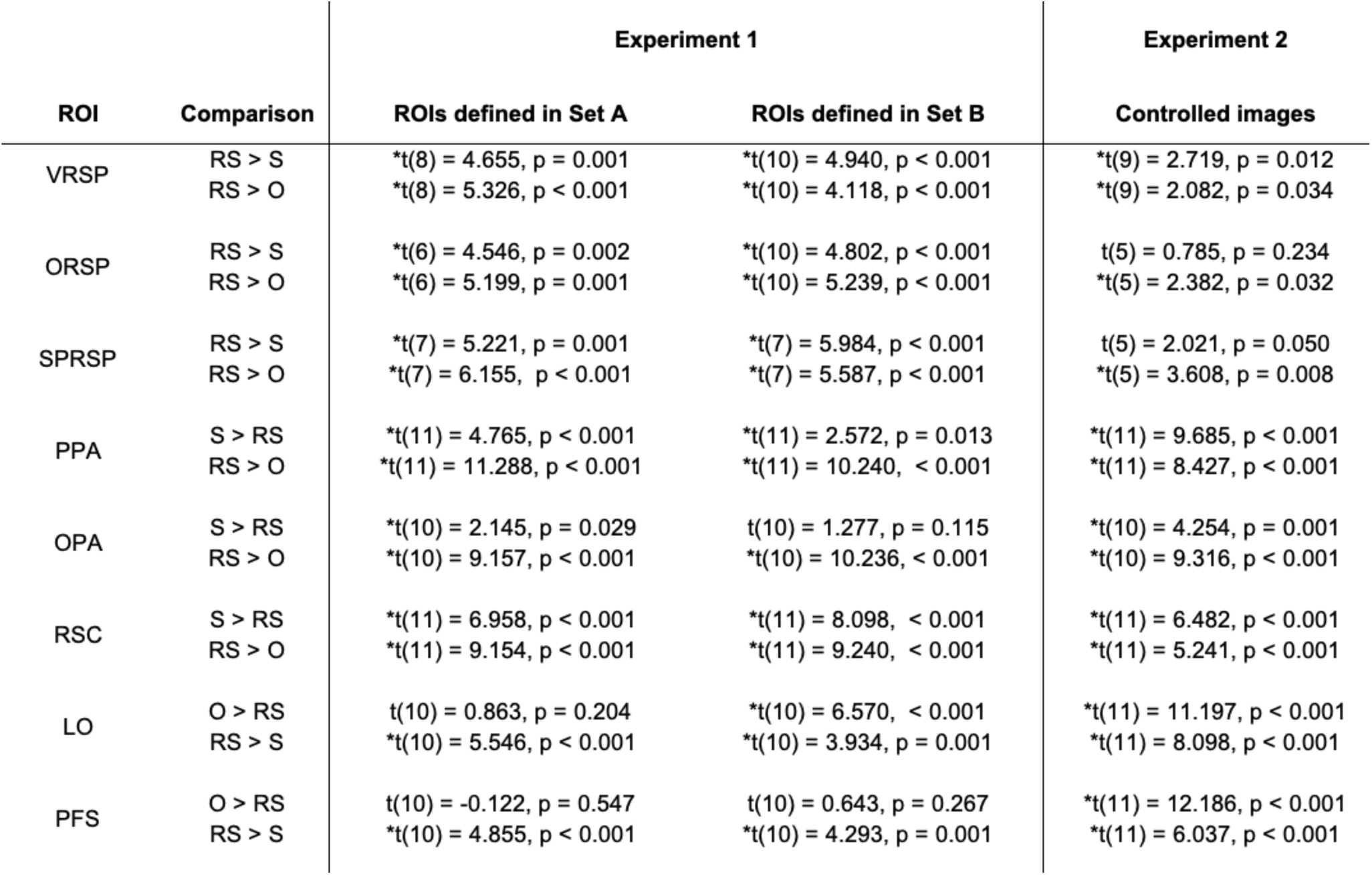
Statistical tests for all pairwise conditions comparisons in all ROIs in Experiments 1 and 2. In Experiment 1, we defined ROIs in Set A and extracted activations from them using Set B. Those statistical test are reported in the main text. Additionally, to confirm that the results were stable, we defined ROIs in Set B and extracted activations from Set B. These statistical results are reported here.

**Supplementary Table 3:**
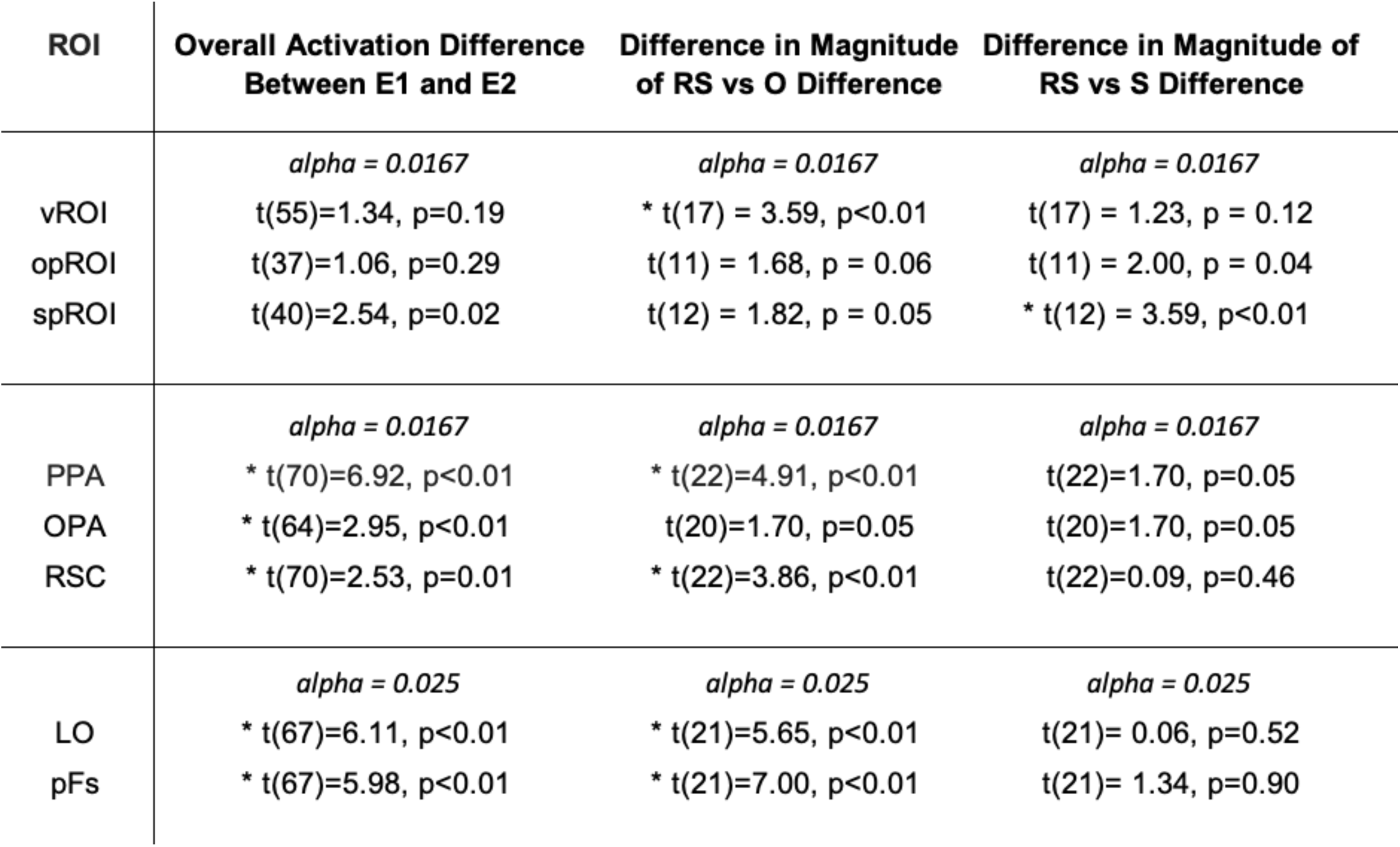
Comparison between activations from original images (E1) vs controlled image sets (E2). **Rows:** reachspace-, scene- and object-preferring ROIs are shown in the three row sections. **First column:** The difference in overall magnitude between original and controlled images is report. These statistics were computed by averaging the overall responses to object, scenes and reachspace conditions, and then comparing these original and controlled image overall activation levels with a t-test. Controlled images elicited numerically smaller responses in all ROIs, though this difference was only significant (at the indicated post-hoc significance threshold) for scene and object ROIs. **Second column:** This column reportss whether the difference between between reachspaces and object conditions changed between original and controlled image sets. Betas for the object condition were subtracted from betas for the reachspace condition for both original (E1) and controlled images (E2), and the results were compared with a t-test. Overall, vROI, PPA and OPA saw a significant decrease in this difference for controlled images, while LO and pFs both saw a significant increase. **Third Column**. This column reports the same analysis as in the 2^nd^ column, instead focusining on the difference between reachspace and scene activations. Overall, only spROI showed a significant difference, with a smaller difference between reachspaces and scenes in controlled images.

**Supplementary Table 4:**
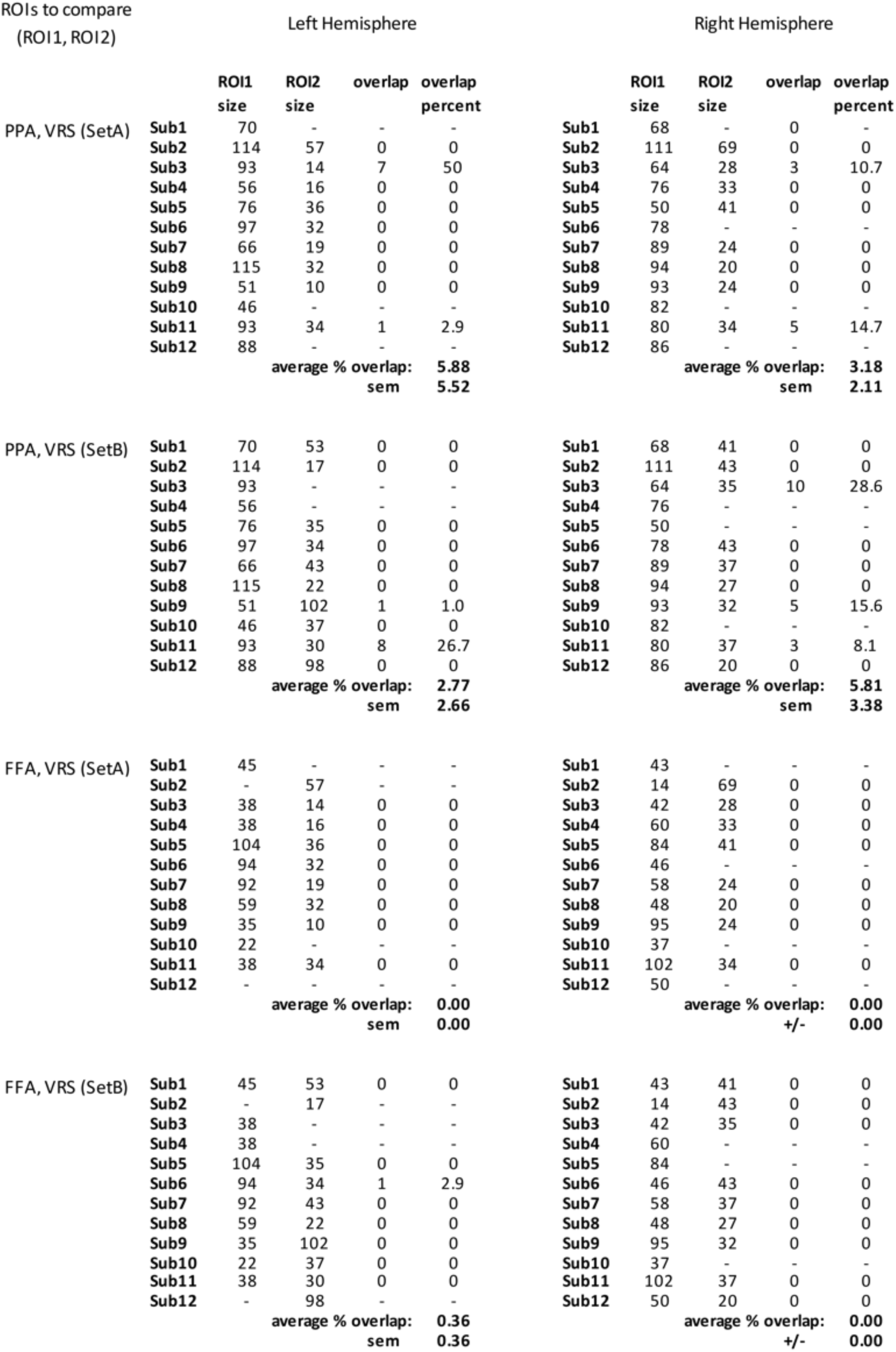
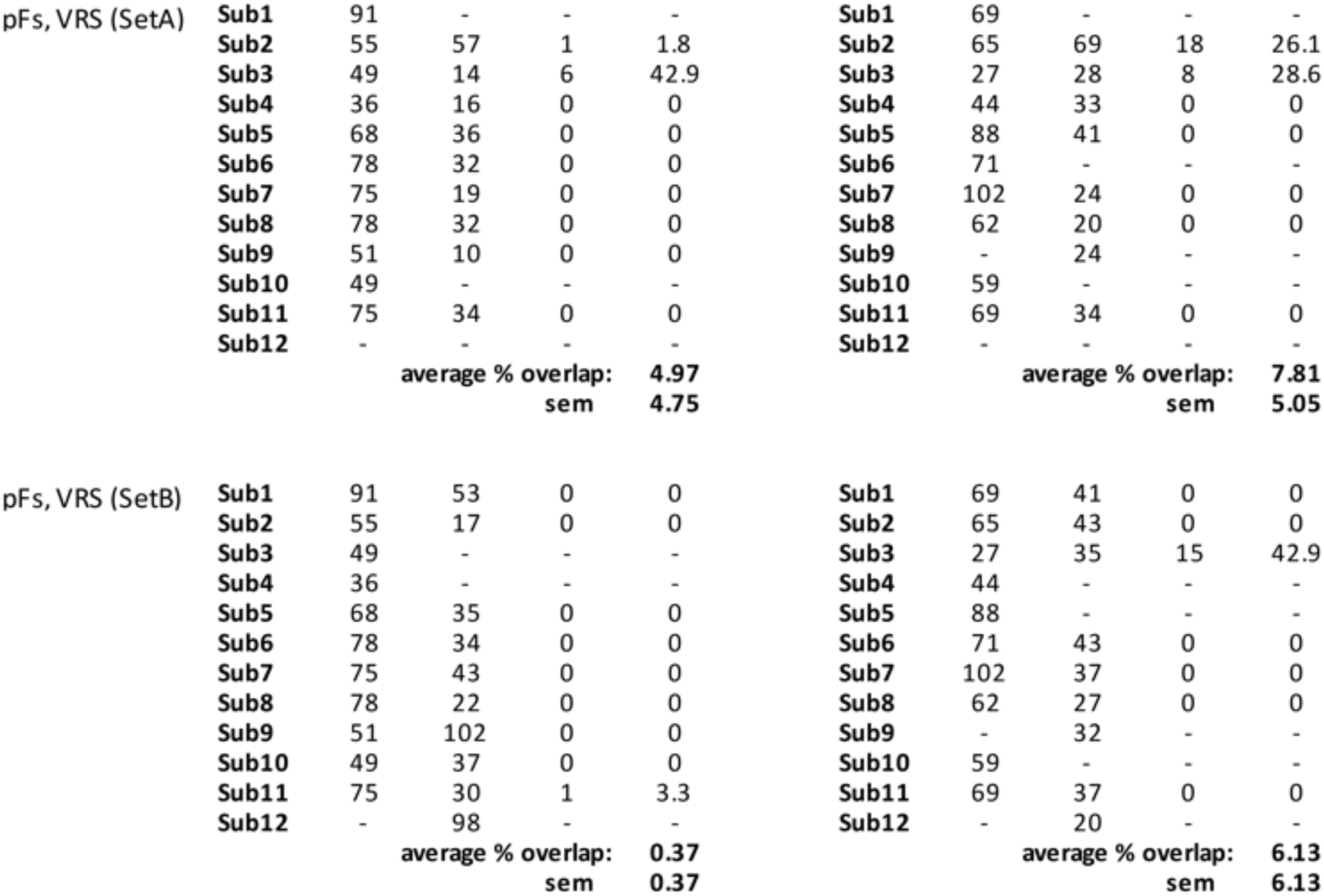
Analysis of voxel overlap between the ventral reachspace-preferring region and other classic ventral ROIs.

**Supplementary Table 5:**
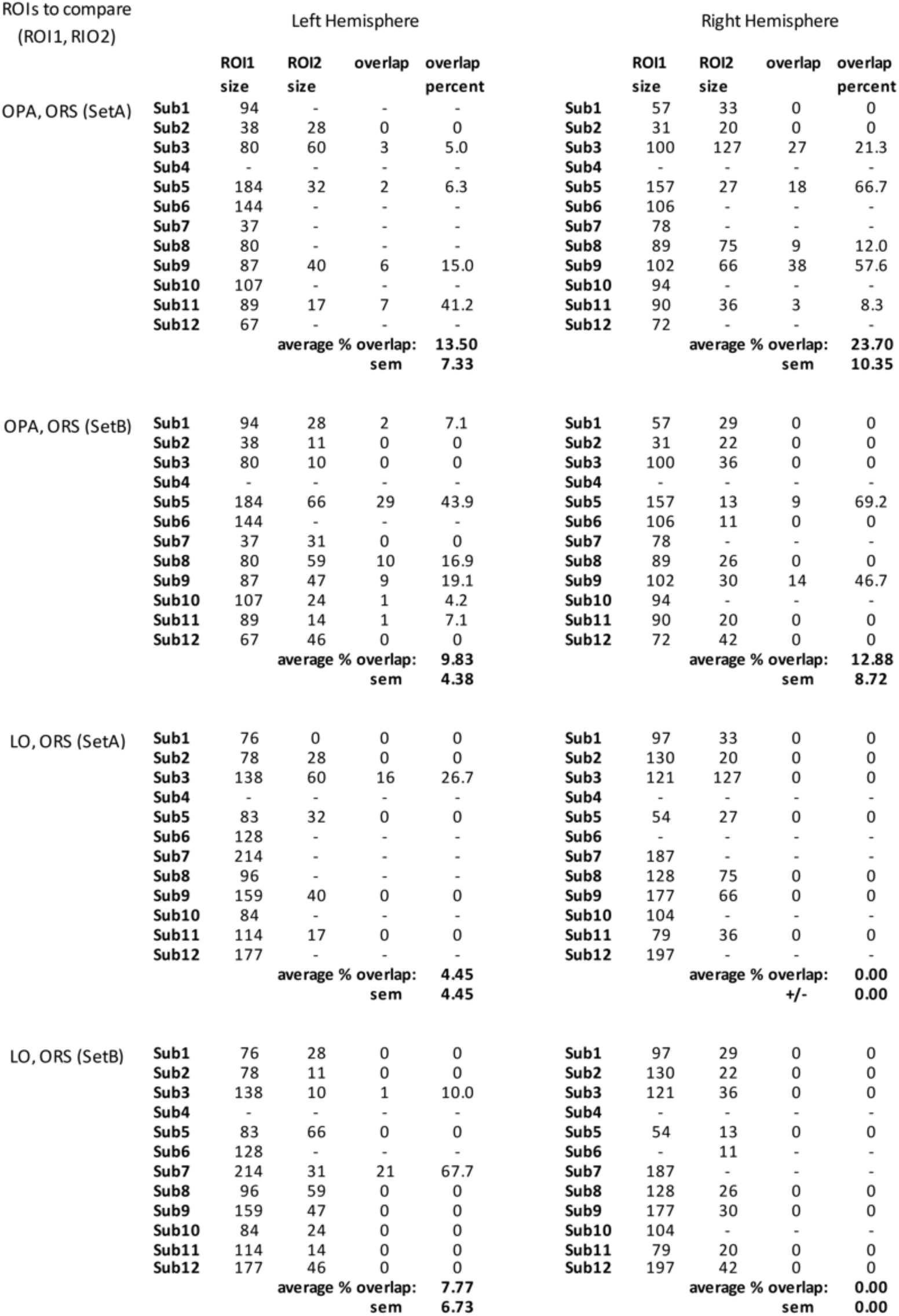
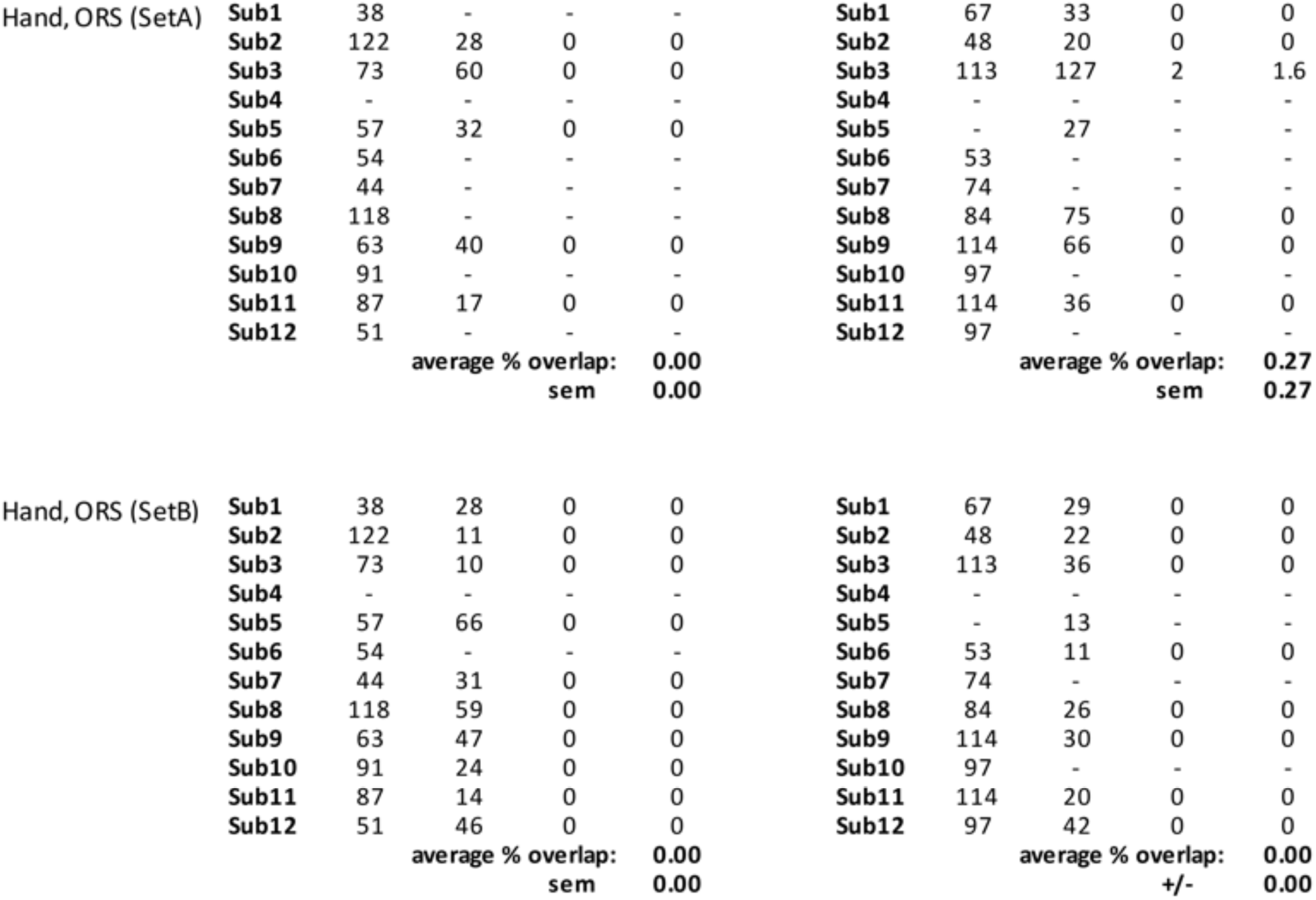
Analysis of voxel overlap between the occipital-partietal reachspace-preferring region and other classic lateral-dorsal ROIs. The superior parietal reachspace region is not shown, as it did not overlap with any ROIs.

