## Supplement for "Large-scale dissociations between views of objects, scenes, and reachable-scale environments in visual cortex"

### Supplementary Figures

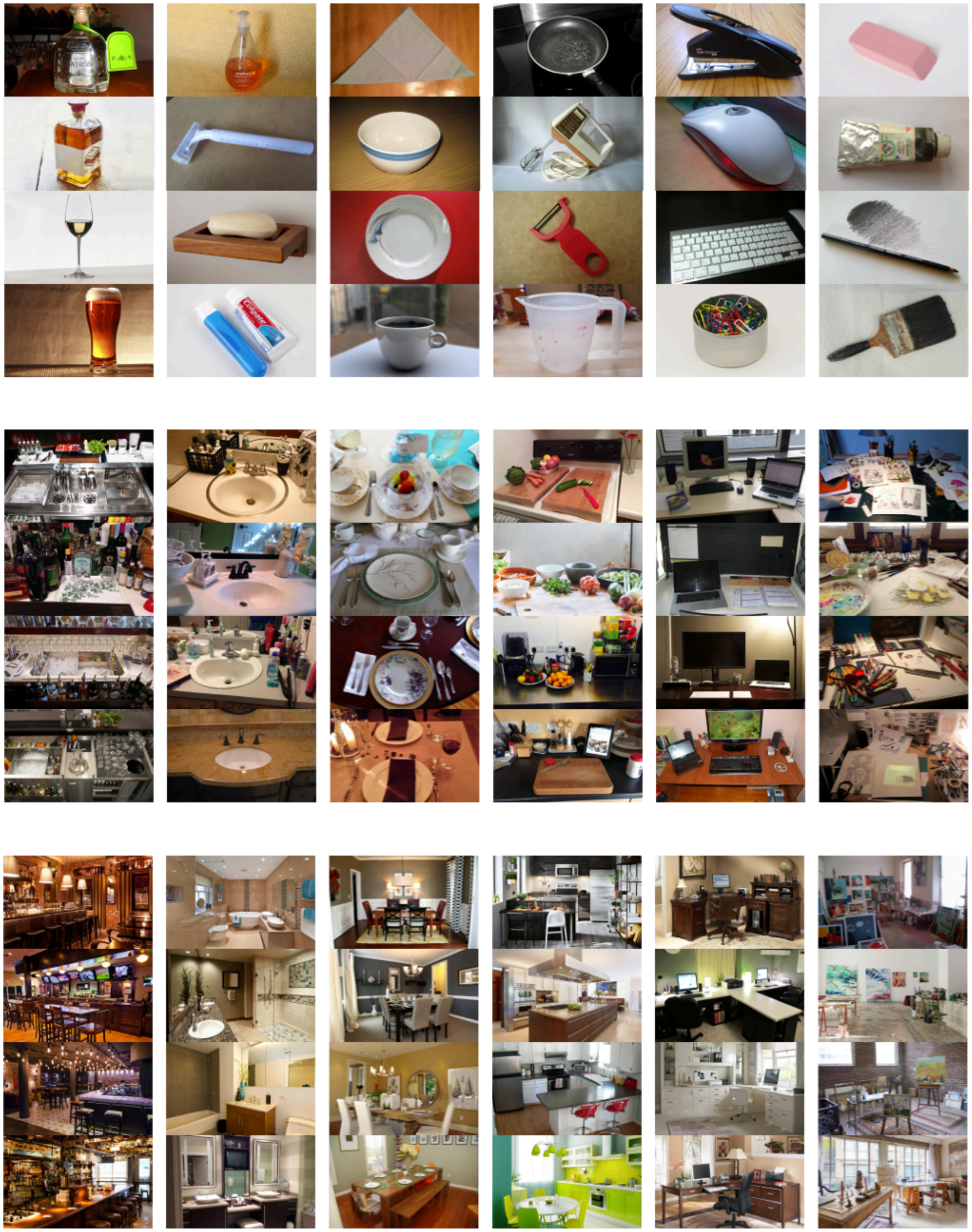

**Supplementary Figure 1:** Additional examples of the object (top), reachspace (middle), and scenes (bottom) stimuli used in Experiment 1

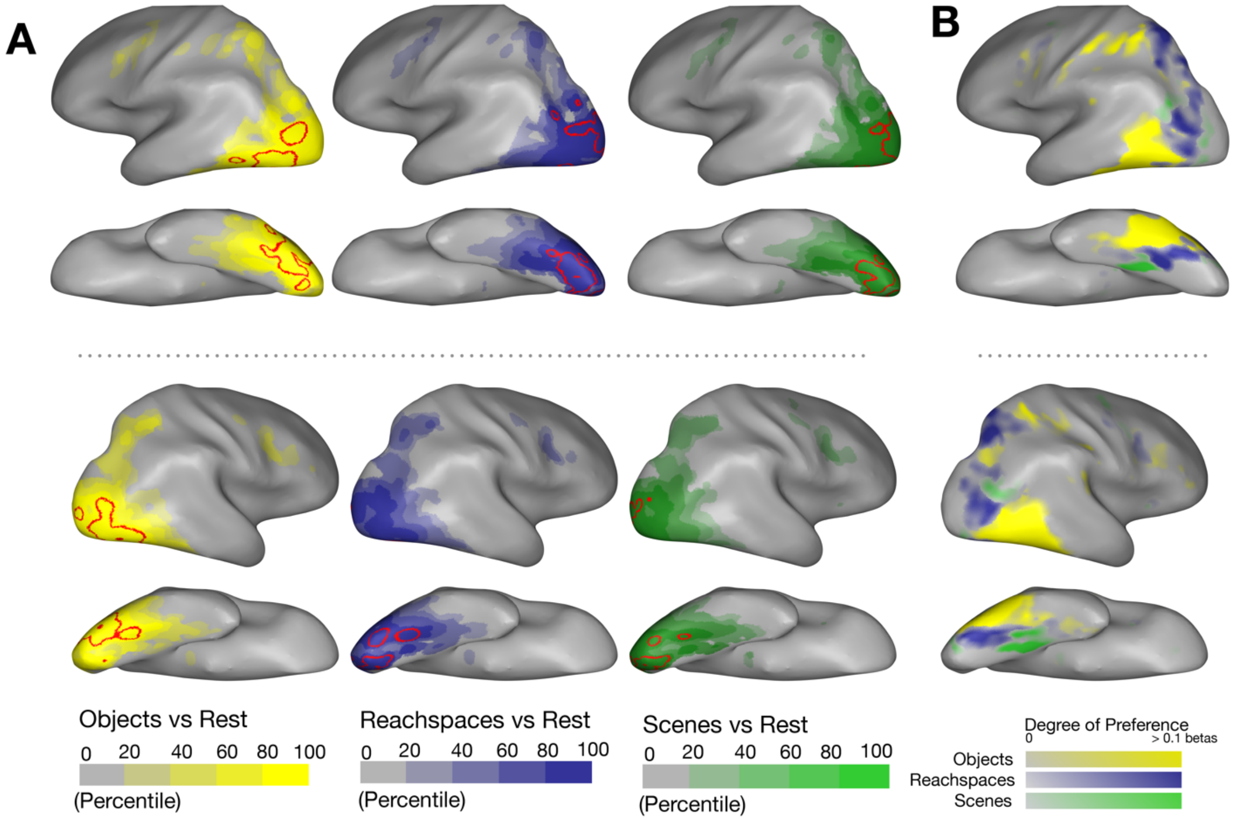


**Supplementary Figure 2:** (A) Overall activations are plotted for each stimulus condition (e.g. objects>rest), for group-level data from Experiment 1. Voxels are colored based on the percentile of activation, where voxels in the 95$^{th}$ percentile are outlined in red. Voxels were included if they exceeded T>2.0 for the contrast of all conditions > rest. Note that these activation percentile can reflect not only the strength of the response, but also differences in signal to noise (e.g. different parts of cortex are closer or farther to the coils and ear-canal artifacts), and should be interpreted with this in mind. (B) The group-level preference map is replotted (data from Figure 1) for comparison. This visualization highlights the fact that the strongest overall activation (e.g. top 5% activations) is in partial but not in perfect correspondence with the regions where objects, reachspaces, and scene images show systematically higher responses.

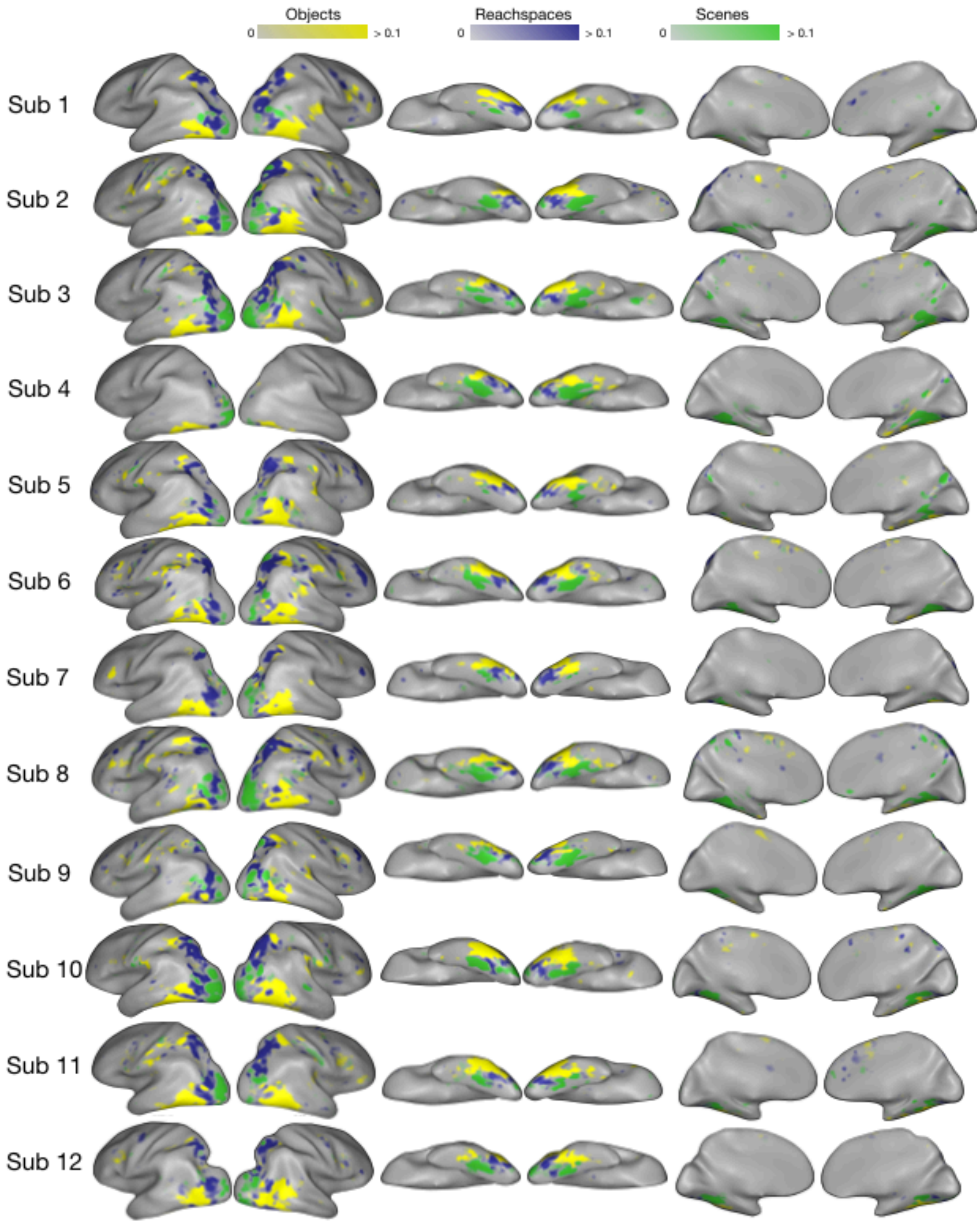


**Supplementary Figure 3**: Single-subject preference maps from Experiment 1.

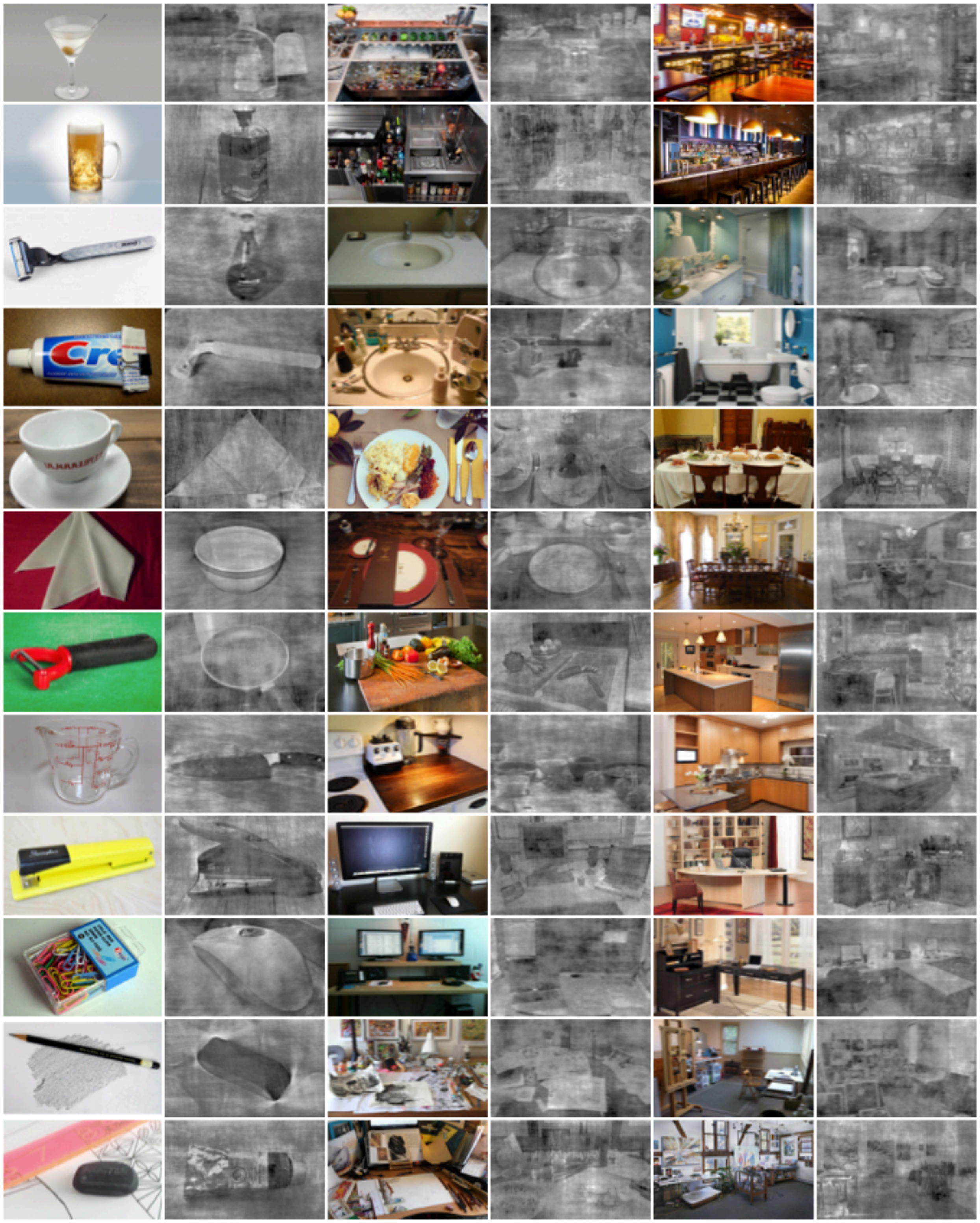

**Supplementary Figure 4:** Additional examples of stimuli for Experiment 2.

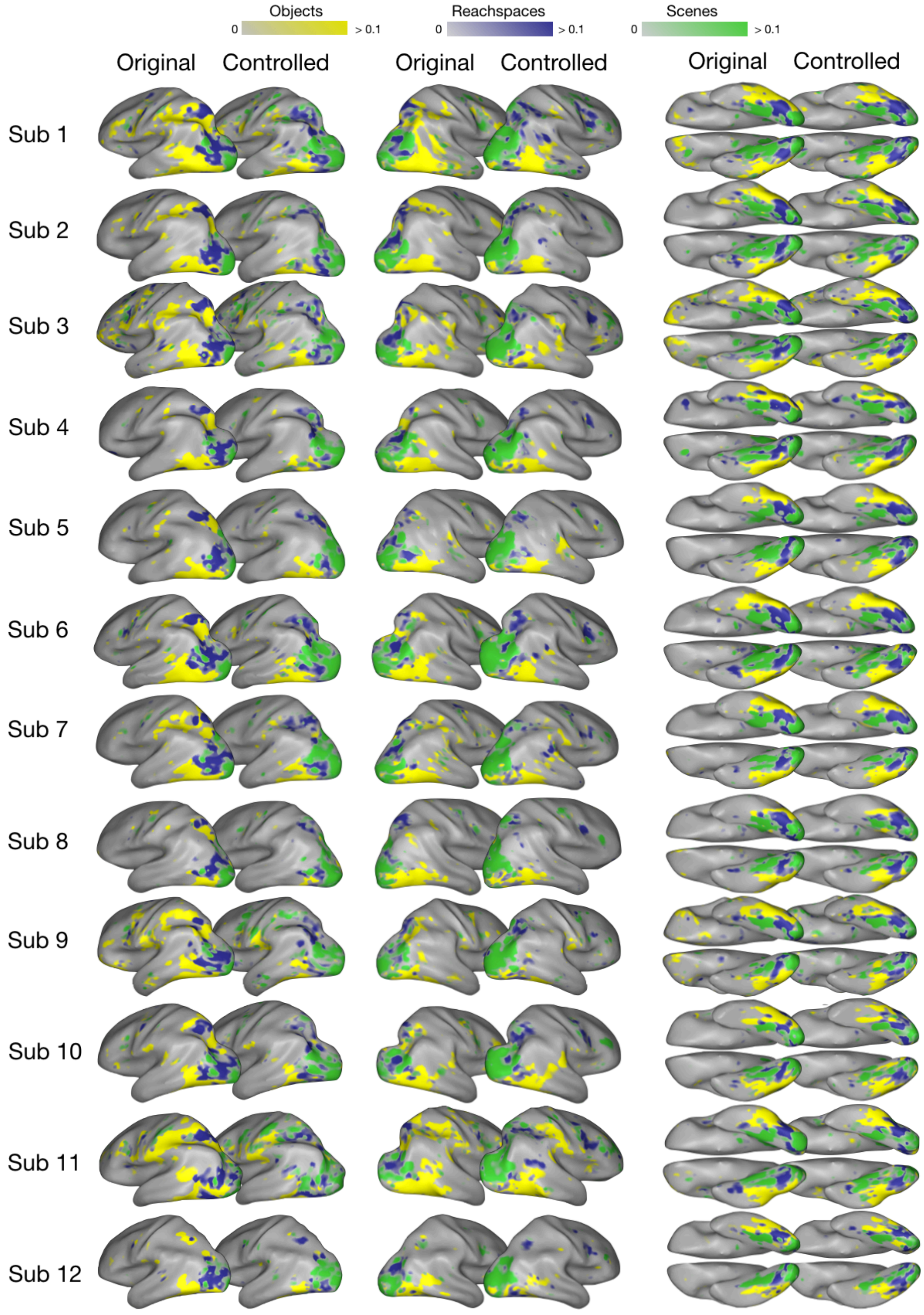


**Supplemental Figure 5:** Single-subject preference maps from Experiment 2, obtained from original and controlled images (same color scale used for both original and controlled)

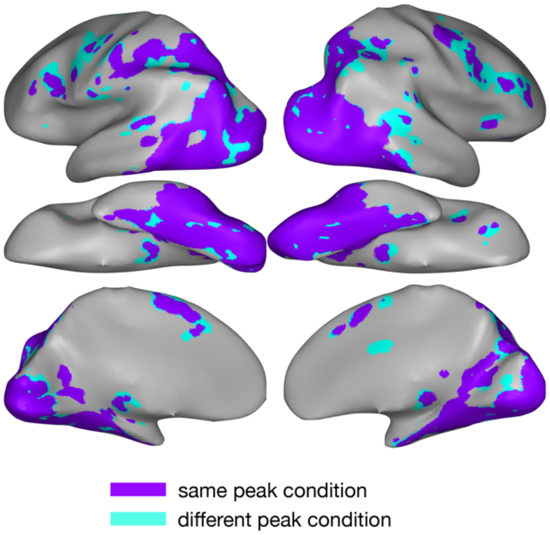


**Supplemental Figure 6:** Comparison between original and controlled group-level preference maps from Experiment 2. Cortex colored in purple showed the same preference for objects, reachspaces, or scene images in both Original and Controlled image sets, while cortex colored in cyan had different peak conditions.

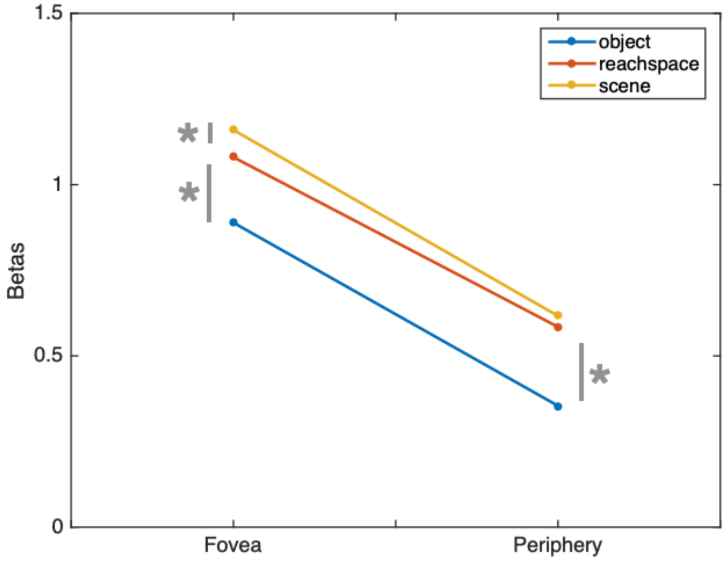


**Supplementary Figure 7.** Responses to objects, reachspaces and scenes in foveal and peripheral regions of early visual cortex (V1-V3). Early visual cortex (EVC) was defined using vertical and horizontal meridians from Experiment 1 eccentricity mapping runs, and then divided it into foveal-preferring and peripheral-preferring regions, based on contrasting the Central vs Peripheral conditions. The overall response (average beta) to objects, reachspaces, and scene images from Experiment 1 was computed. The y-axis plots overall response for both foveal and peripheral regions (x-axis), for the three different stimulus conditions. Post-hoc paired t-tests indicated that in foveal cortex, scenes elicited the most activity, followed by reachspaces images and then object objects. In peripheral cortex, there was no statistically significant difference between scenes and reachspaces, but both these conditions showed relatively greater activation than object images.

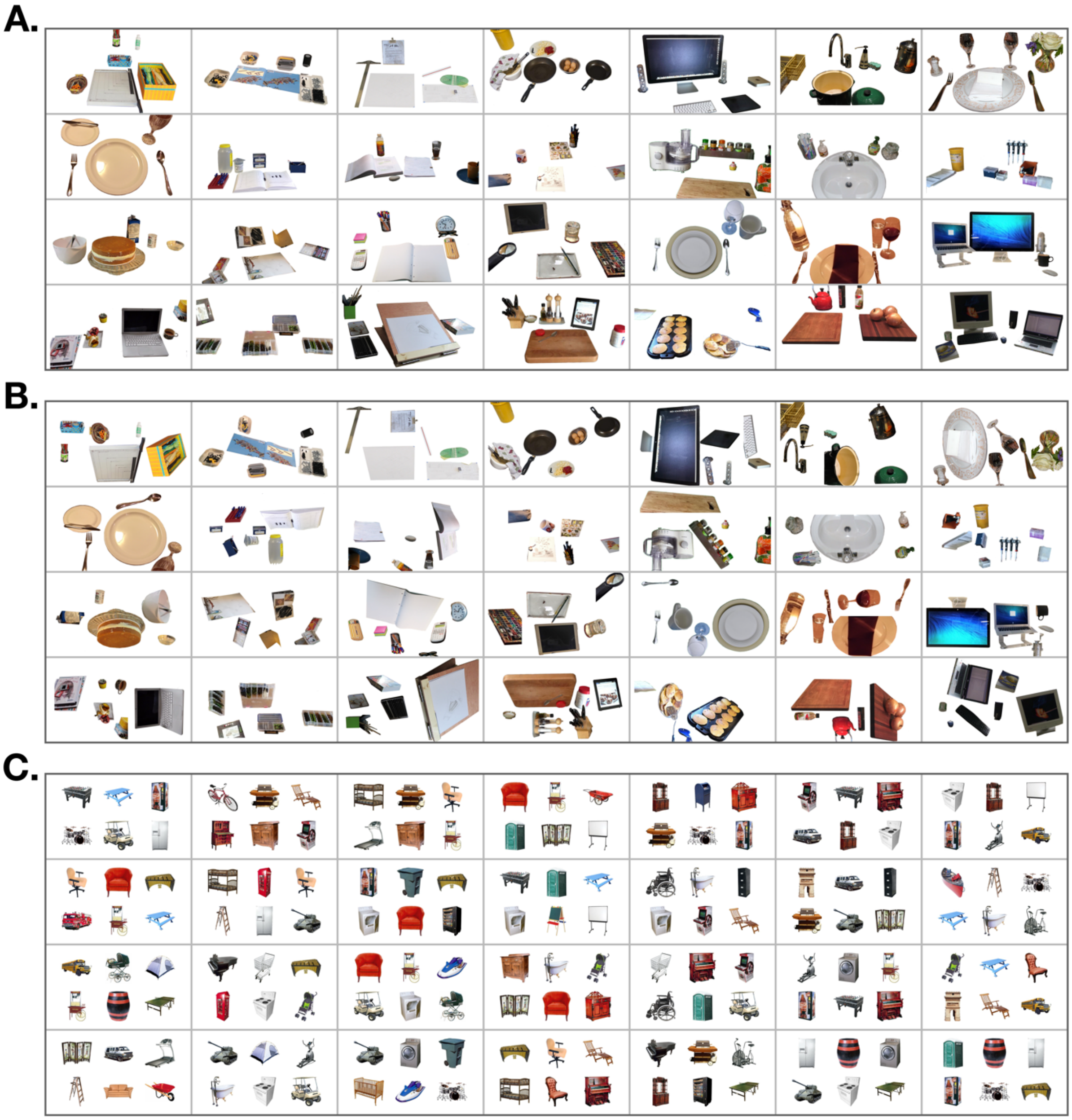

**Supplementary Figure 8**. Experiment 3 stimuli. A) reachspaces images with the background removed in Photoshop, yielding images of multiple objects in realistic spatial arrangements; B) reachspaces images with background removed and the remaining objects scrambled to disrupt their spatial arrangement; C) 6 objects with large real-world size (e.g. trampoline, dresser) arranged in a 3x2 grid one a white background.

D.

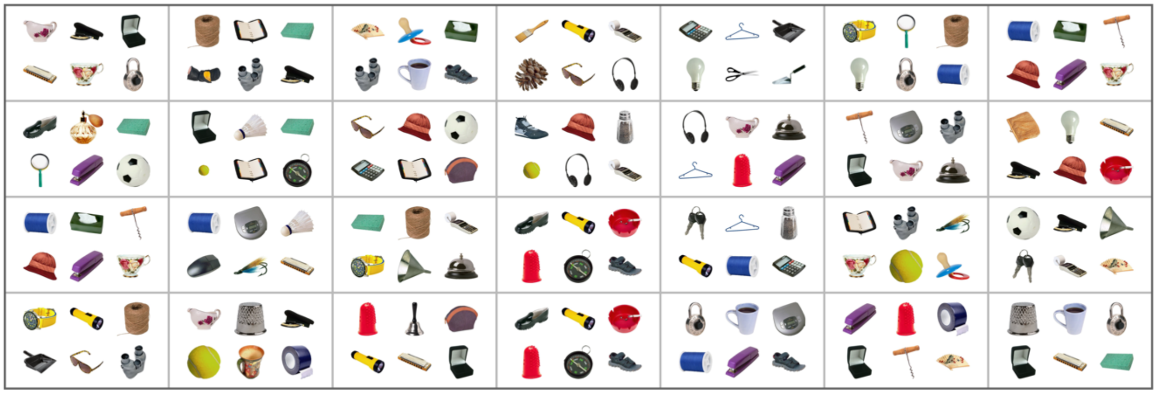

E.

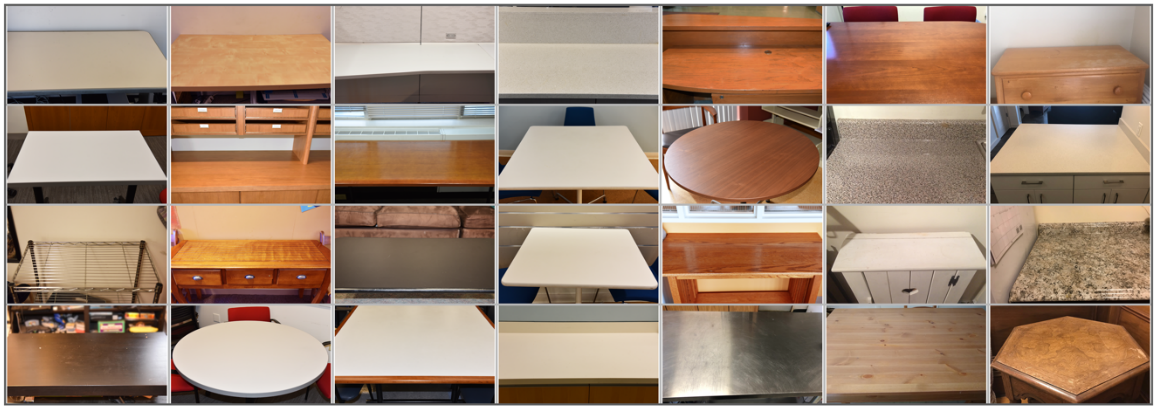

F.

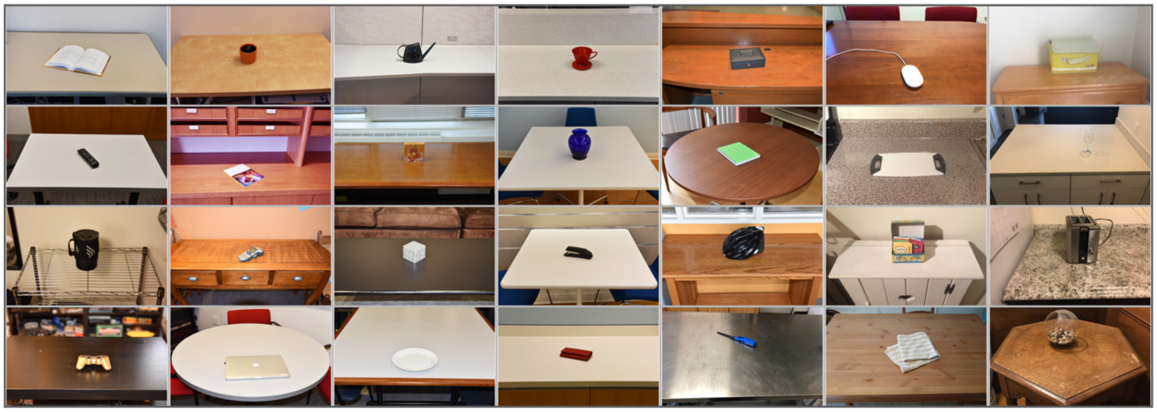

**Supplementary Figure 8 (continued)**. Experiment 3 stimuli. D) 6 objects with small real world size (e.g. mug, watch) arranged in a 3x2 grid a white background (presented at the same visual size as the large object condition); E) reachable environments with all objects removed except the support surface; F) reachspaces containing only a single object on the support surface.

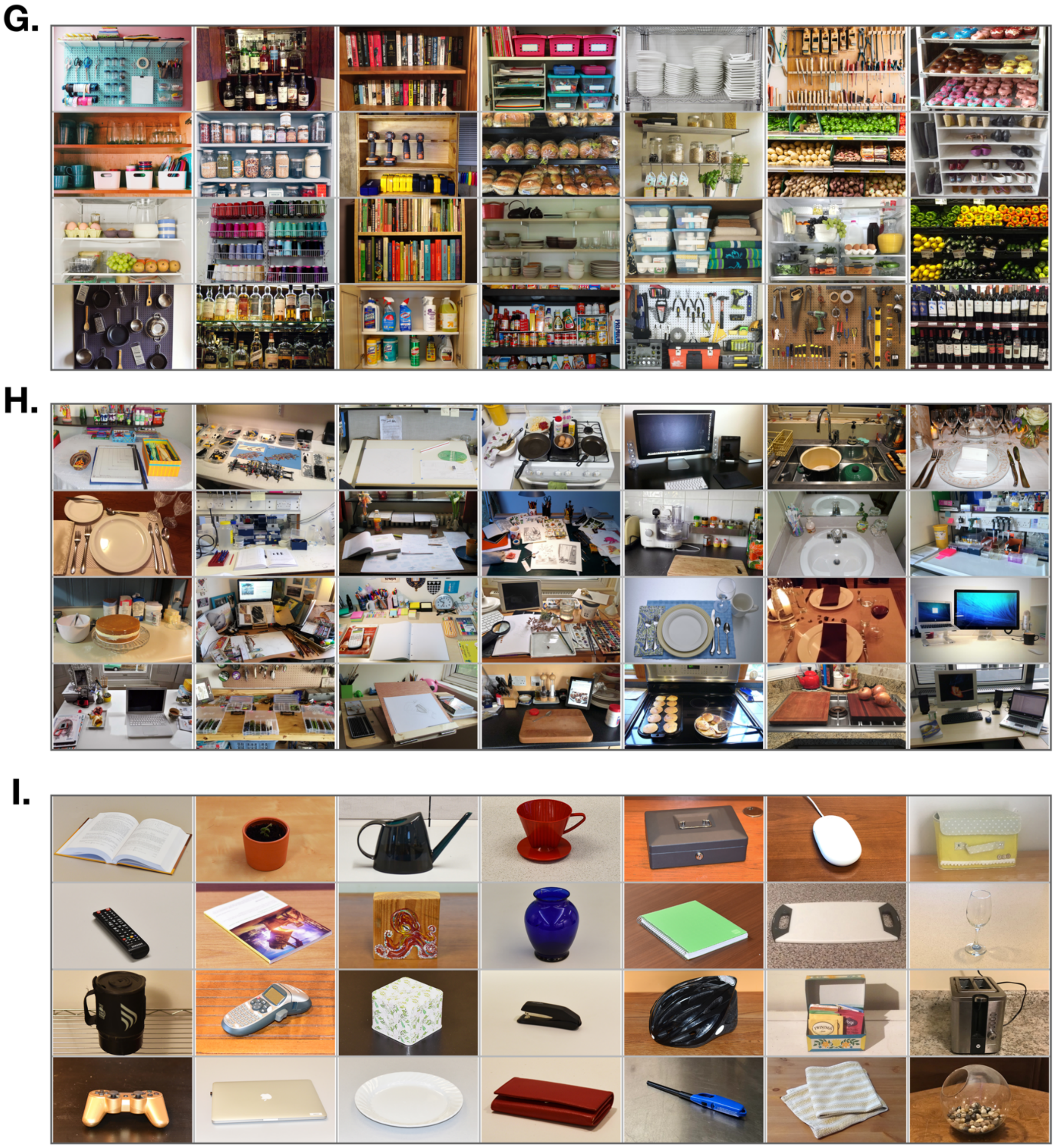

**Supplementary Figure 8 (continued)**. Experiment 3 stimuli. G) vertical reachspaces, where the disposition of objects was vertical rather than horizontal ( e.g. shelves, peg-boards); H) regular (i.e. horizontal) reachspaces; I) objects (i.e. close-up views of single objects on their natural background).

J.

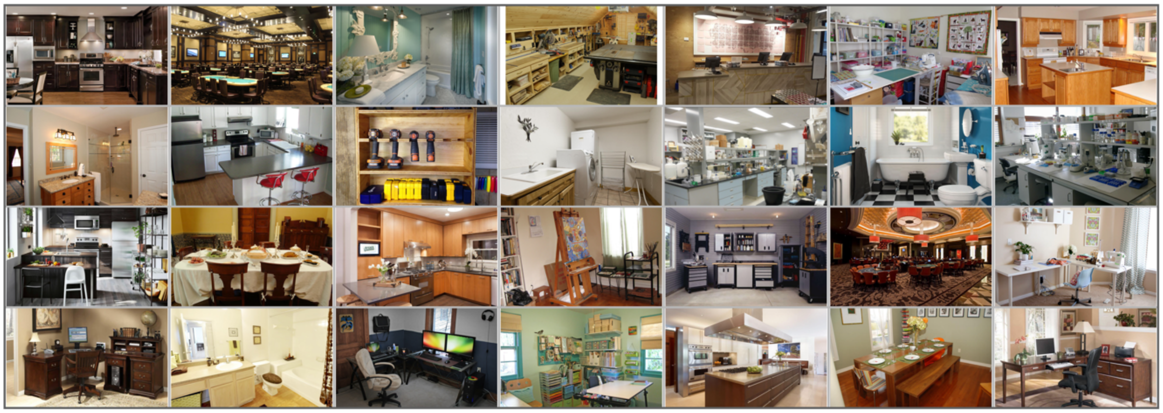

**Supplementary Figure 8 (continued)**. Experiment 3 stimuli, continued. J) Scene images.

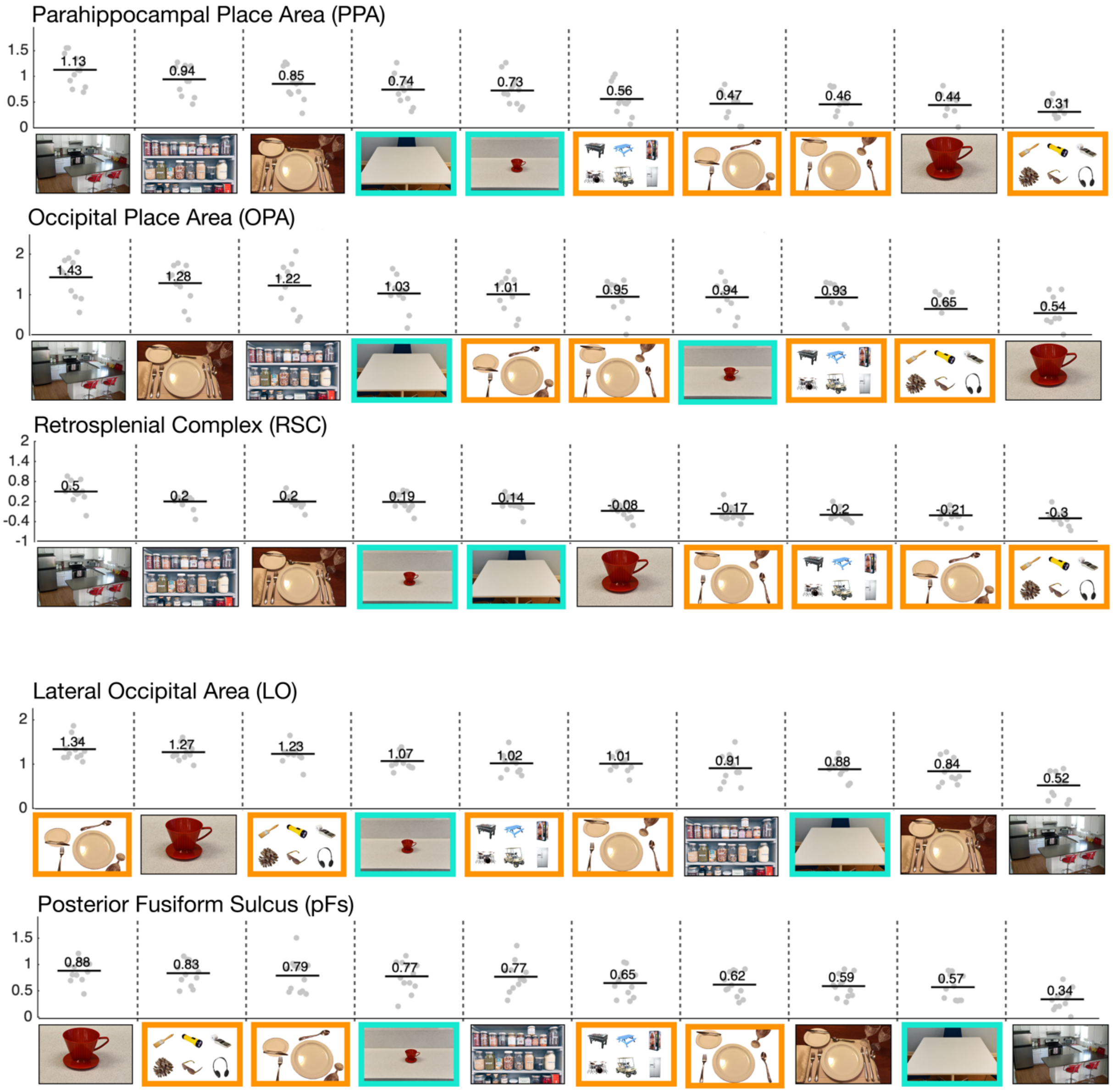


**Supplementary Figure 9**. Experiment 3 results for scene- and object-selective ROIs. Responses in scene and object-preferring ROIs across all stimulus conditions are shown, with conditions plotted in order from highest to lowest activations. Images with orange borders indicate stimuli dominated by multiple objects, and images with teal borders highlight images of reachable space with low object content. The mean activation is indicated with a black horizontal bar; gray points indicate single participant data.

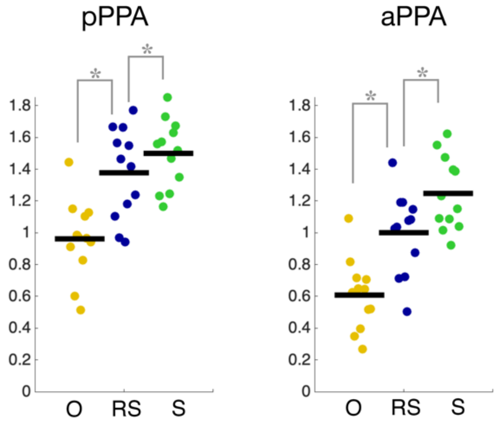


**Supplementary Figure 10:** Responses to objects, reachspaces and scenes in PPA subdivisions. In each subject, PPA was divided in half along its anterior-posterior axis, and responses were assessed in each half. While overall activations were higher in posterior PPA (pPPA) than anterior PPA (aPPA), both showed the same pattern of results: scenes elicited greater activation than reachspaces (aPPA: $t(11) = 6.22$, $p<0.01$ 0.00; pPPA: $t(11) = 2.49$, $p = 0.015$ ), and reachspaces elicited greater activation than objects (aPPA: $t(11) = 11.51$, $p < 0.01$; pPPA: $t(11) = 9.71$, $p <0.01$ ),

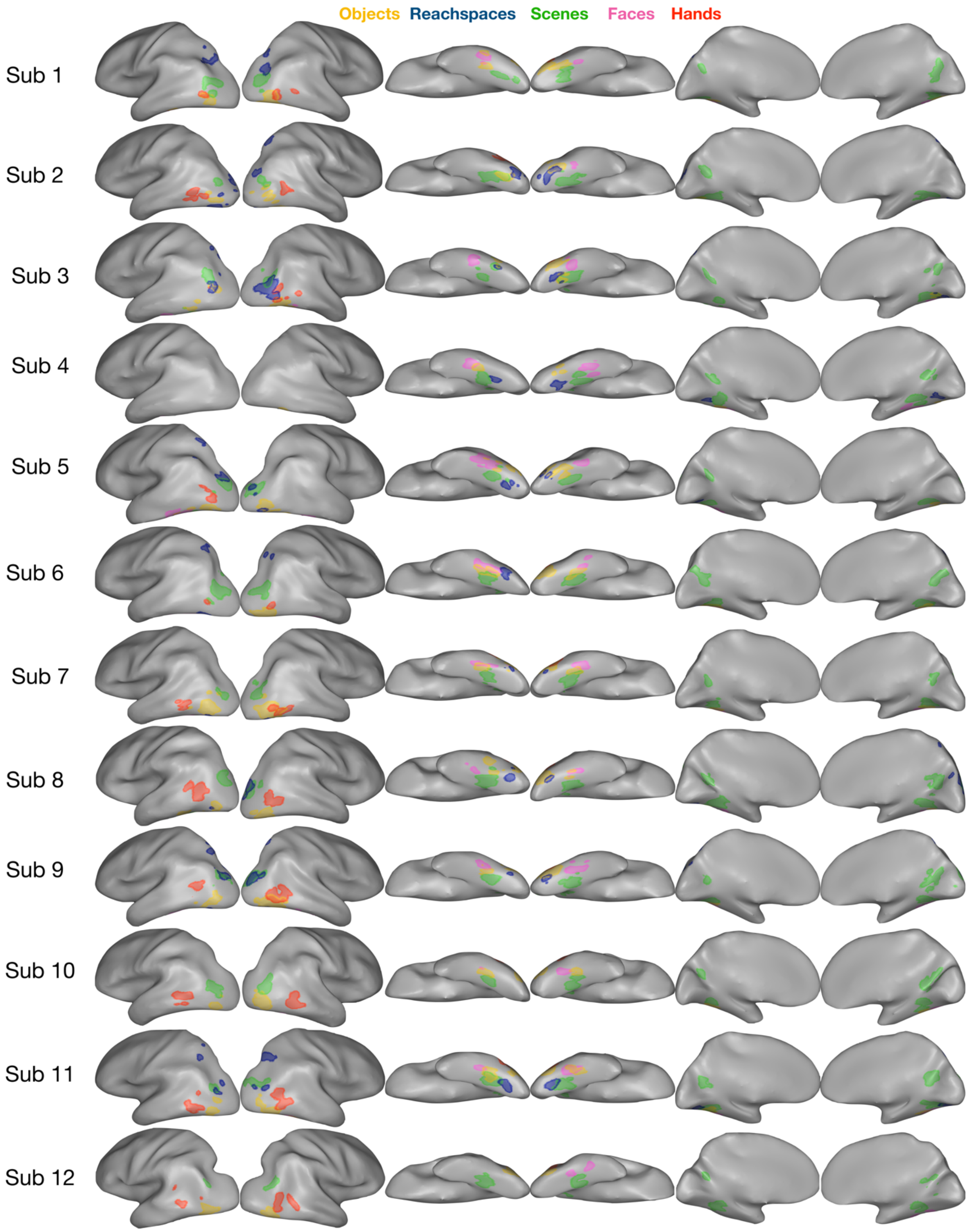


**Supplementary Figure 11.** All Experiment 1 ROIs viewed in single-subjects. The color legend is as follows: yellow for objects, blue for reachspaces, green for scenes, pink for faces, orange for hands.

#### Supplementary Analysis: Population Mixing

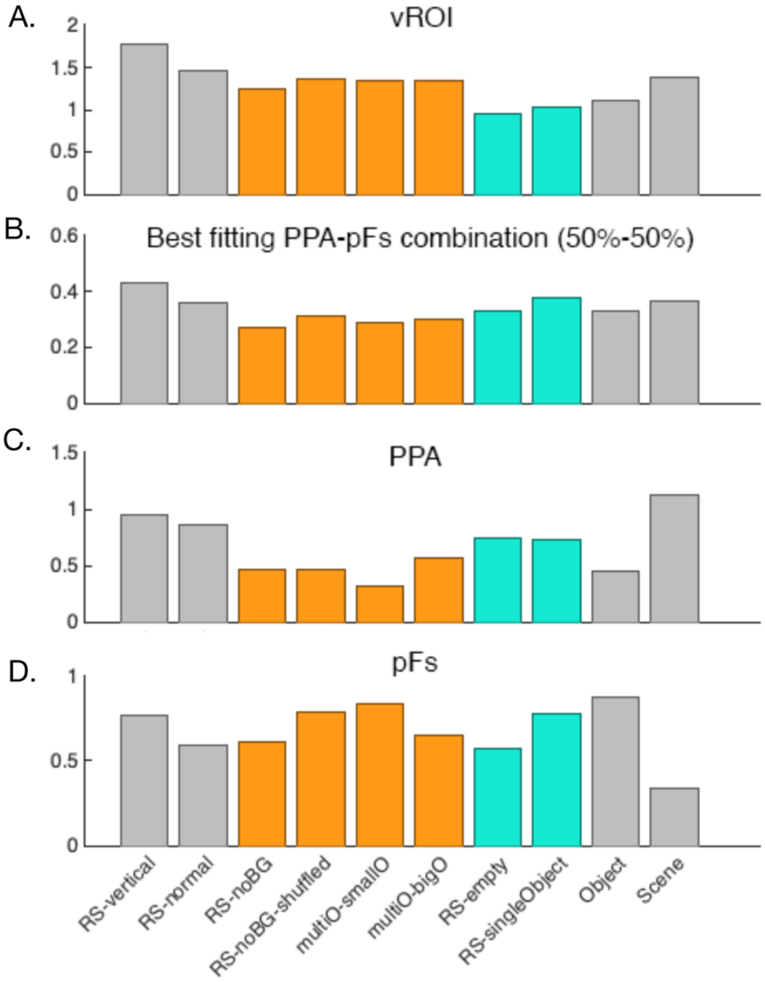


| PPA weight | pFS weight | Pearson | Spearman |
|---|---|---|---|
| 0.1 | 0.9 | r =0.03 | r=0.02 |
| 0.2 | 0.8 | r=0.12 | r=0.07 |
| 0.3 | 0.7 | r=0.26 | r=0.13 |
| 0.4 | 0.6 | r=0.38 | r=0.24 |
| 0.5 | 0.5 | **r=0.42** | r=0.31 |
| 0.6 | 0.4 | r=0.40 | r=0.43 |
| 0.7 | 0.8 | r=0.38 | r=0.43 |
| 0.8 | 0.2 | r=0.36 | **r=0.48** |
| 0.9 | 0.1 | r=0.34 | **r=0.48** |

**Supplementary Figure 12.** (A): The actual 10-condition response profile of the ventral reachspace-preferring ROI is shown (y-axis: betas; x-axis, conditions). (B) The best fitting PPA-pFS profile combination is shown. (C,D). The profiles for PPA and pFS are also shown. Table shows Pearson and Spearman correlations for all tested weightings of PPA and pFs with vROI.

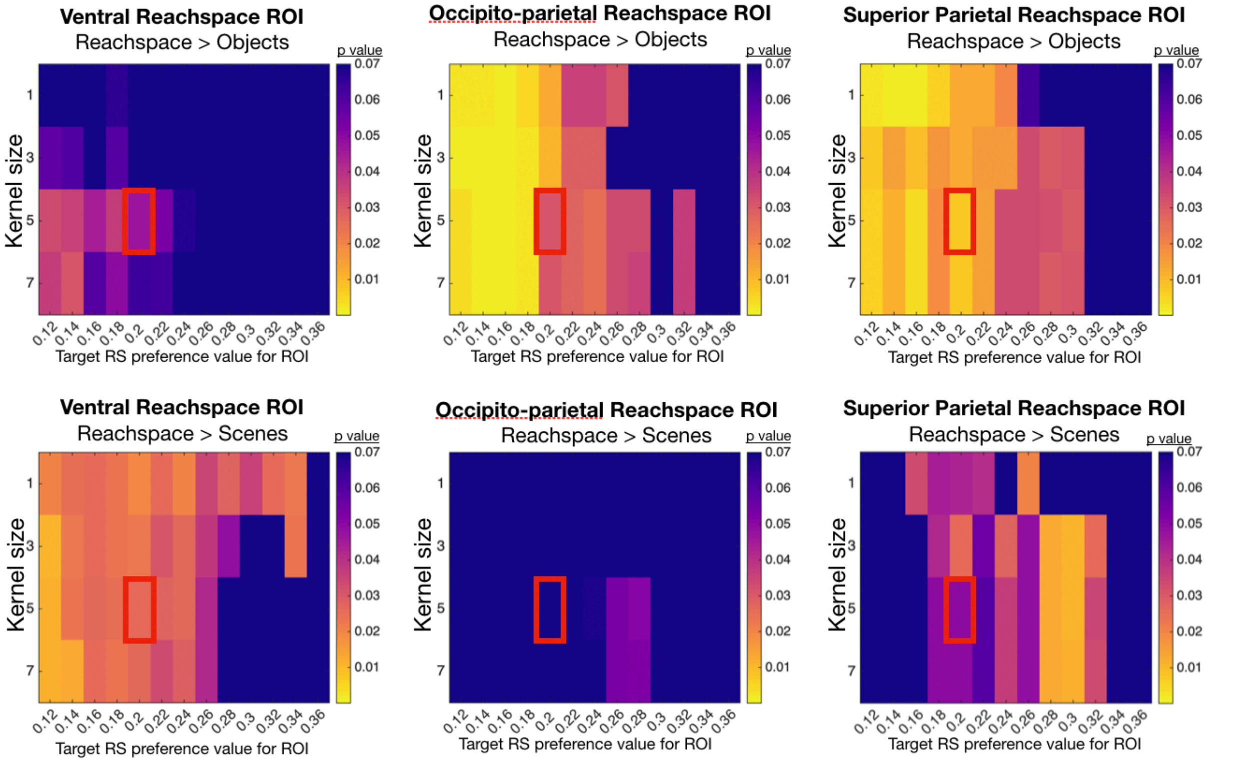


**Supplemental Figure 13:** Visualization of how Experiment 1 results vary as the automatic ROI-selection parameters are varied. For Experiment 2, reachspace-preferring ROIs were selecting using a semi-automatic procedure (see Methods). Parameters such as the size of the smoothing kernel and the reachspace-preference threshold value were determined a priori, based on analyses run in a separate set of data. Results in the main text were extracted from ROIs defined using a 5-voxel smoothing kernel and requiring a reachspace preference of 0.2 betas. Here, we display how the statistics significance of the preference for reachspaces over objects (top row) and scenes (bottom row) would have changed with different parameters. In each graph, the rows vary the size of the smoothing kernels applied to the statistical maps computed from the conjunction contrast RS>O & RS>S, from 1 to 7 voxels. The columns vary the threshold for the beta value of reachspace preference we required of the final ROI, from 0.12 to 0.36. The color in each cell shows the statistical significance of the comparison indicated in the title. The red square shows the cell corresponding to the parameters used in the main text.

#### Supplementary Tables

| ROI | Hemi | Exp 1 Set A | | | Exp 1 Set B | | | Exp2 Auto ROIs | | |
|---|---|---|---|---|---|---|---|---|---|---|
| | | *x* | *y* | *z* | *x* | *y* | *z* | *x* | *z* | *z* |
| vROI | LH | -28 | -69 | -9 | -24 | -70 | -11 | -28 | -67 | -12 |
| | RH | 26 | -71 | -10 | 20 | -71 | -11 | 24 | -71 | -14 |
| opROI | LH | -24 | -82 | 17 | -29 | -82 | 16 | -28 | -81 | 17 |
| | RH | 27 | -82 | 18 | 25 | -80 | 20 | 27 | -79 | 17 |
| spROI | LH | -23 | -65 | 48 | -21 | -65 | 51 | -21 | -66 | 48 |
| | RH | 17 | -69 | 52 | 15 | -66 | 57 | 14 | -67 | 53 |

**Supplementary Table 1:** Average TAL coordinates for reachspace-preferring ROIs in all experiments.

| | | Experiment 1 | | Experiment 2 |
|---|---|---|---|---|
| ROI | Comparison | ROIs defined in Set A | ROIs defined in Set B | Controlled images |
| VRSP | RS > S | *t(8) = 4.655, p = 0.001 | *t(10) = 4.940, p < 0.001 | *t(9) = 2.719, p = 0.012 |
| | RS > O | *t(8) = 5.326, p < 0.001 | *t(10) = 4.118, p < 0.001 | *t(9) = 2.082, p = 0.034 |
| ORSP | RS > S | *t(6) = 4.546, p = 0.002 | *t(10) = 4.802, p < 0.001 | t(5) = 0.785, p = 0.234 |
| | RS > O | *t(6) = 5.199, p = 0.001 | *t(10) = 5.239, p < 0.001 | *t(5) = 2.382, p = 0.032 |
| SPRSP | RS > S | *t(7) = 5.221, p = 0.001 | *t(7) = 5.984, p < 0.001 | t(5) = 2.021, p = 0.050 |
| | RS > O | *t(7) = 6.155, p < 0.001 | *t(7) = 5.587, p < 0.001 | *t(5) = 3.608, p = 0.008 |
| PPA | S > RS | *t(11) = 4.765, p < 0.001 | *t(11) = 2.572, p = 0.013 | *t(11) = 9.685, p < 0.001 |
| | RS > O | *t(11) = 11.288, p < 0.001 | *t(11) = 10.240, < 0.001 | *t(11) = 8.427, p < 0.001 |
| OPA | S > RS | *t(10) = 2.145, p = 0.029 | t(10) = 1.277, p = 0.115 | *t(10) = 4.254, p = 0.001 |
| | RS > O | *t(10) = 9.157, p < 0.001 | *t(10) = 10.236, < 0.001 | *t(10) = 9.316, p < 0.001 |
| RSC | S > RS | *t(11) = 6.958, p < 0.001 | *t(11) = 8.098, < 0.001 | *t(11) = 6.482, p < 0.001 |
| | RS > O | *t(11) = 9.154, p < 0.001 | *t(11) = 9.240, < 0.001 | *t(11) = 5.241, p < 0.001 |
| LO | O > RS | t(10) = 0.863, p = 0.204 | *t(10) = 6.570, < 0.001 | *t(11) = 11.197, p < 0.001 |
| | RS > S | *t(10) = 5.546, p < 0.001 | *t(10) = 3.934, p = 0.001 | *t(11) = 8.098, p < 0.001 |
| PFS | O > RS | t(10) = -0.122, p = 0.547 | t(10) = 0.643, p = 0.267 | *t(11) = 12.186, p < 0.001 |
| | RS > S | *t(10) = 4.855, p < 0.001 | *t(10) = 4.293, p = 0.001 | *t(11) = 6.037, p < 0.001 |

**Supplementary Table 2:** Statistical tests for all pairwise conditions comparisons in all ROIs in Experiments 1 and 2. In Experiment 1, we defined ROIs in Set A and extracted activations from them using Set B. Those statistical test are reported in the main text. Additionally, to confirm that the results were stable, we defined ROIs in Set B and extracted activations from Set B. These statistical results are reported here.

| ROI | Overall Activation Difference Between E1 and E2 | Difference in Magnitude of RS vs O Difference | Difference in Magnitude of RS vs S Difference |
|---|---|---|---|
| | *alpha = 0.0167* | *alpha = 0.0167* | *alpha = 0.0167* |
| vROI | t(55)=1.34, p=0.19 | * t(17) = 3.59, p<0.01 | t(17) = 1.23, p = 0.12 |
| opROI | t(37)=1.06, p=0.29 | t(11) = 1.68, p = 0.06 | t(11) = 2.00, p = 0.04 |
| spROI | t(40)=2.54, p=0.02 | t(12) = 1.82, p = 0.05 | * t(12) = 3.59, p<0.01 |
| | *alpha = 0.0167* | *alpha = 0.0167* | *alpha = 0.0167* |
| PPA | * t(70)=6.92, p<0.01 | * t(22)=4.91, p<0.01 | t(22)=1.70, p=0.05 |
| OPA | * t(64)=2.95, p<0.01 | t(20)=1.70, p=0.05 | t(20)=1.70, p=0.05 |
| RSC | * t(70)=2.53, p=0.01 | * t(22)=3.86, p<0.01 | t(22)=0.09, p=0.46 |
| | *alpha = 0.025* | *alpha = 0.025* | *alpha = 0.025* |
| LO | * t(67)=6.11, p<0.01 | * t(21)=5.65, p<0.01 | t(21)= 0.06, p=0.52 |
| pFs | * t(67)=5.98, p<0.01 | * t(21)=7.00, p<0.01 | t(21)= 1.34, p=0.90 |

**Supplementary Table 3:** Comparison between activations from original images (E1) vs controlled image sets (E2). **Rows**: reachspace-, scene- and object-preferring ROIs are shown in the three row sections. **First column**: The difference in overall magnitude between original and controlled images is report. These statistics were computed by averaging the overall responses to object, scenes and reachspace conditions, and then comparing these original and controlled image overall activation levels with a t-test. Controlled images elicited numerically smaller responses in all ROIs, though this difference was only significant (at the indicated post-hoc significance threshold) for scene and object ROIs. **Second column**: This column reportss whether the difference between between reachspaces and object conditions changed between original and controlled image sets. Betas for the object condition were subtracted from betas for the reachspace condition for both original (E1) and controlled images (E2), and the results were compared with a t-test. Overall, vROI, PPA and OPA saw a significant decrease in this difference for controlled images, while LO and pFs both saw a significant increase. **Third Column**. This column reports the same analysis as in the 2$^{nd}$ column, instead focusining on the difference between reachspace and scene activations. Overall, only spROI showed a significant difference, with a smaller difference between reachspaces and scenes in controlled images.

| ROIs to compare (ROI1, ROI2) | | Left Hemisphere | | | | | Right Hemisphere | | | |
|---|---|---|---|---|---|---|---|---|---|---|
| | | **ROI1 size** | **ROI2 size** | **overlap** | **overlap percent** | | **ROI1 size** | **ROI2 size** | **overlap** | **overlap percent** |
| PPA, VRS (SetA) | **Sub1** | 70 | - | - | - | **Sub1** | 68 | - | 0 | - |
| | **Sub2** | 114 | 57 | 0 | 0 | **Sub2** | 111 | 69 | 0 | 0 |
| | **Sub3** | 93 | 14 | 7 | 50 | **Sub3** | 64 | 28 | 3 | 10.7 |
| | **Sub4** | 56 | 16 | 0 | 0 | **Sub4** | 76 | 33 | 0 | 0 |
| | **Sub5** | 76 | 36 | 0 | 0 | **Sub5** | 50 | 41 | 0 | 0 |
| | **Sub6** | 97 | 32 | 0 | 0 | **Sub6** | 78 | - | - | - |
| | **Sub7** | 66 | 19 | 0 | 0 | **Sub7** | 89 | 24 | 0 | 0 |
| | **Sub8** | 115 | 32 | 0 | 0 | **Sub8** | 94 | 20 | 0 | 0 |
| | **Sub9** | 51 | 10 | 0 | 0 | **Sub9** | 93 | 24 | 0 | 0 |
| | **Sub10** | 46 | - | - | - | **Sub10** | 82 | - | - | - |
| | **Sub11** | 93 | 34 | 1 | 2.9 | **Sub11** | 80 | 34 | 5 | 14.7 |
| | **Sub12** | 88 | - | - | - | **Sub12** | 86 | - | - | - |
| | | | | **average % overlap:** | **5.88** | | | | **average % overlap:** | **3.18** |
| | | | | **sem** | **5.52** | | | | **sem** | **2.11** |
| PPA, VRS (SetB) | **Sub1** | 70 | 53 | 0 | 0 | **Sub1** | 68 | 41 | 0 | 0 |
| | **Sub2** | 114 | 17 | 0 | 0 | **Sub2** | 111 | 43 | 0 | 0 |
| | **Sub3** | 93 | - | - | - | **Sub3** | 64 | 35 | 10 | 28.6 |
| | **Sub4** | 56 | - | - | - | **Sub4** | 76 | - | - | - |
| | **Sub5** | 76 | 35 | 0 | 0 | **Sub5** | 50 | - | - | - |
| | **Sub6** | 97 | 34 | 0 | 0 | **Sub6** | 78 | 43 | 0 | 0 |
| | **Sub7** | 66 | 43 | 0 | 0 | **Sub7** | 89 | 37 | 0 | 0 |
| | **Sub8** | 115 | 22 | 0 | 0 | **Sub8** | 94 | 27 | 0 | 0 |
| | **Sub9** | 51 | 102 | 1 | 1.0 | **Sub9** | 93 | 32 | 5 | 15.6 |
| | **Sub10** | 46 | 37 | 0 | 0 | **Sub10** | 82 | - | - | - |
| | **Sub11** | 93 | 30 | 8 | 26.7 | **Sub11** | 80 | 37 | 3 | 8.1 |
| | **Sub12** | 88 | 98 | 0 | 0 | **Sub12** | 86 | 20 | 0 | 0 |
| | | | | **average % overlap:** | **2.77** | | | | **average % overlap:** | **5.81** |
| | | | | **sem** | **2.66** | | | | **sem** | **3.38** |
| FFA, VRS (SetA) | **Sub1** | 45 | - | - | - | **Sub1** | 43 | - | - | - |
| | **Sub2** | - | 57 | - | - | **Sub2** | 14 | 69 | 0 | 0 |
| | **Sub3** | 38 | 14 | 0 | 0 | **Sub3** | 42 | 28 | 0 | 0 |
| | **Sub4** | 38 | 16 | 0 | 0 | **Sub4** | 60 | 33 | 0 | 0 |
| | **Sub5** | 104 | 36 | 0 | 0 | **Sub5** | 84 | 41 | 0 | 0 |
| | **Sub6** | 94 | 32 | 0 | 0 | **Sub6** | 46 | - | - | - |
| | **Sub7** | 92 | 19 | 0 | 0 | **Sub7** | 58 | 24 | 0 | 0 |
| | **Sub8** | 59 | 32 | 0 | 0 | **Sub8** | 48 | 20 | 0 | 0 |
| | **Sub9** | 35 | 10 | 0 | 0 | **Sub9** | 95 | 24 | 0 | 0 |
| | **Sub10** | 22 | - | - | - | **Sub10** | 37 | - | - | - |
| | **Sub11** | 38 | 34 | 0 | 0 | **Sub11** | 102 | 34 | 0 | 0 |
| | **Sub12** | - | - | - | - | **Sub12** | 50 | - | - | - |
| | | | | **average % overlap:** | **0.00** | | | | **average % overlap:** | **0.00** |
| | | | | **sem** | **0.00** | | | | **+/-** | **0.00** |
| FFA, VRS (SetB) | **Sub1** | 45 | 53 | 0 | 0 | **Sub1** | 43 | 41 | 0 | 0 |
| | **Sub2** | - | 17 | - | - | **Sub2** | 14 | 43 | 0 | 0 |
| | **Sub3** | 38 | - | - | - | **Sub3** | 42 | 35 | 0 | 0 |
| | **Sub4** | 38 | - | - | - | **Sub4** | 60 | - | - | - |
| | **Sub5** | 104 | 35 | 0 | 0 | **Sub5** | 84 | - | - | - |
| | **Sub6** | 94 | 34 | 1 | 2.9 | **Sub6** | 46 | 43 | 0 | 0 |
| | **Sub7** | 92 | 43 | 0 | 0 | **Sub7** | 58 | 37 | 0 | 0 |
| | **Sub8** | 59 | 22 | 0 | 0 | **Sub8** | 48 | 27 | 0 | 0 |
| | **Sub9** | 35 | 102 | 0 | 0 | **Sub9** | 95 | 32 | 0 | 0 |
| | **Sub10** | 22 | 37 | 0 | 0 | **Sub10** | 37 | - | - | - |
| | **Sub11** | 38 | 30 | 0 | 0 | **Sub11** | 102 | 37 | 0 | 0 |
| | **Sub12** | - | 98 | - | - | **Sub12** | 50 | 20 | 0 | 0 |
| | | | | **average % overlap:** | **0.36** | | | | **average % overlap:** | **0.00** |
| | | | | **sem** | **0.36** | | | | **+/-** | **0.00** |

**Supplementary Table 4:** Analysis of voxel overlap between the ventral reachspace-preferring region and other classic ventral ROIs.

| | | | | | | | | | |
|---|---|---|---|---|---|---|---|---|---|
| pFs, VRS (SetA) | Sub1 | 91 | - | - | - | Sub1 | 69 | - | - | - |
| | Sub2 | 55 | 57 | 1 | 1.8 | Sub2 | 65 | 69 | 18 | 26.1 |
| | Sub3 | 49 | 14 | 6 | 42.9 | Sub3 | 27 | 28 | 8 | 28.6 |
| | Sub4 | 36 | 16 | 0 | 0 | Sub4 | 44 | 33 | 0 | 0 |
| | Sub5 | 68 | 36 | 0 | 0 | Sub5 | 88 | 41 | 0 | 0 |
| | Sub6 | 78 | 32 | 0 | 0 | Sub6 | 71 | - | - | - |
| | Sub7 | 75 | 19 | 0 | 0 | Sub7 | 102 | 24 | 0 | 0 |
| | Sub8 | 78 | 32 | 0 | 0 | Sub8 | 62 | 20 | 0 | 0 |
| | Sub9 | 51 | 10 | 0 | 0 | Sub9 | - | 24 | - | - |
| | Sub10 | 49 | - | - | - | Sub10 | 59 | - | - | - |
| | Sub11 | 75 | 34 | 0 | 0 | Sub11 | 69 | 34 | 0 | 0 |
| | Sub12 | - | - | - | - | Sub12 | - | - | - | - |
| | | | average % overlap: | | 4.97 | | | average % overlap: | | 7.81 |
| | | | | sem | 4.75 | | | | sem | 5.05 |

| | | | | | | | | | | |
|---|---|---|---|---|---|---|---|---|---|---|
| pFs, VRS (SetB) | Sub1 | 91 | 53 | 0 | 0 | Sub1 | 69 | 41 | 0 | 0 |
| | Sub2 | 55 | 17 | 0 | 0 | Sub2 | 65 | 43 | 0 | 0 |
| | Sub3 | 49 | - | - | - | Sub3 | 27 | 35 | 15 | 42.9 |
| | Sub4 | 36 | - | - | - | Sub4 | 44 | - | - | - |
| | Sub5 | 68 | 35 | 0 | 0 | Sub5 | 88 | - | - | - |
| | Sub6 | 78 | 34 | 0 | 0 | Sub6 | 71 | 43 | 0 | 0 |
| | Sub7 | 75 | 43 | 0 | 0 | Sub7 | 102 | 37 | 0 | 0 |
| | Sub8 | 78 | 22 | 0 | 0 | Sub8 | 62 | 27 | 0 | 0 |
| | Sub9 | 51 | 102 | 0 | 0 | Sub9 | - | 32 | - | - |
| | Sub10 | 49 | 37 | 0 | 0 | Sub10 | 59 | - | - | - |
| | Sub11 | 75 | 30 | 1 | 3.3 | Sub11 | 69 | 37 | 0 | 0 |
| | Sub12 | - | 98 | - | - | Sub12 | - | 20 | - | - |
| | | | average % overlap: | | 0.37 | | | average % overlap: | | 6.13 |
| | | | | sem | 0.37 | | | | sem | 6.13 |

**Supplementary Table 4 (continued):** Analysis of voxel overlap between the ventral reachspace-preferring region and other classic ventral ROIs.

| ROIs to compare (ROI1, RIO2) | | Left Hemisphere | | | | | Right Hemisphere | | | |
|---|---|---|---|---|---|---|---|---|---|---|
| | | ROI1 size | ROI2 size | overlap | overlap percent | | ROI1 size | ROI2 size | overlap | overlap percent |
| OPA, ORS (SetA) | **Sub1** | 94 | - | - | - | **Sub1** | 57 | 33 | 0 | 0 |
| | **Sub2** | 38 | 28 | 0 | 0 | **Sub2** | 31 | 20 | 0 | 0 |
| | **Sub3** | 80 | 60 | 3 | 5.0 | **Sub3** | 100 | 127 | 27 | 21.3 |
| | **Sub4** | - | - | - | - | **Sub4** | - | - | - | - |
| | **Sub5** | 184 | 32 | 2 | 6.3 | **Sub5** | 157 | 27 | 18 | 66.7 |
| | **Sub6** | 144 | - | - | - | **Sub6** | 106 | - | - | - |
| | **Sub7** | 37 | - | - | - | **Sub7** | 78 | - | - | - |
| | **Sub8** | 80 | - | - | - | **Sub8** | 89 | 75 | 9 | 12.0 |
| | **Sub9** | 87 | 40 | 6 | 15.0 | **Sub9** | 102 | 66 | 38 | 57.6 |
| | **Sub10** | 107 | - | - | - | **Sub10** | 94 | - | - | - |
| | **Sub11** | 89 | 17 | 7 | 41.2 | **Sub11** | 90 | 36 | 3 | 8.3 |
| | **Sub12** | 67 | - | - | - | **Sub12** | 72 | - | - | - |
| | | | **average % overlap:** | | **13.50** | | | **average % overlap:** | | **23.70** |
| | | | | **sem** | **7.33** | | | | **sem** | **10.35** |
| OPA, ORS (SetB) | **Sub1** | 94 | 28 | 2 | 7.1 | **Sub1** | 57 | 29 | 0 | 0 |
| | **Sub2** | 38 | 11 | 0 | 0 | **Sub2** | 31 | 22 | 0 | 0 |
| | **Sub3** | 80 | 10 | 0 | 0 | **Sub3** | 100 | 36 | 0 | 0 |
| | **Sub4** | - | - | - | - | **Sub4** | - | - | - | - |
| | **Sub5** | 184 | 66 | 29 | 43.9 | **Sub5** | 157 | 13 | 9 | 69.2 |
| | **Sub6** | 144 | - | - | - | **Sub6** | 106 | 11 | 0 | 0 |
| | **Sub7** | 37 | 31 | 0 | 0 | **Sub7** | 78 | - | - | - |
| | **Sub8** | 80 | 59 | 10 | 16.9 | **Sub8** | 89 | 26 | 0 | 0 |
| | **Sub9** | 87 | 47 | 9 | 19.1 | **Sub9** | 102 | 30 | 14 | 46.7 |
| | **Sub10** | 107 | 24 | 1 | 4.2 | **Sub10** | 94 | - | - | - |
| | **Sub11** | 89 | 14 | 1 | 7.1 | **Sub11** | 90 | 20 | 0 | 0 |
| | **Sub12** | 67 | 46 | 0 | 0 | **Sub12** | 72 | 42 | 0 | 0 |
| | | | **average % overlap:** | | **9.83** | | | **average % overlap:** | | **12.88** |
| | | | | **sem** | **4.38** | | | | **sem** | **8.72** |
| LO, ORS (SetA) | **Sub1** | 76 | 0 | 0 | 0 | **Sub1** | 97 | 33 | 0 | 0 |
| | **Sub2** | 78 | 28 | 0 | 0 | **Sub2** | 130 | 20 | 0 | 0 |
| | **Sub3** | 138 | 60 | 16 | 26.7 | **Sub3** | 121 | 127 | 0 | 0 |
| | **Sub4** | - | - | - | - | **Sub4** | - | - | - | - |
| | **Sub5** | 83 | 32 | 0 | 0 | **Sub5** | 54 | 27 | 0 | 0 |
| | **Sub6** | 128 | - | - | - | **Sub6** | - | - | - | - |
| | **Sub7** | 214 | - | - | - | **Sub7** | 187 | - | - | - |
| | **Sub8** | 96 | - | - | - | **Sub8** | 128 | 75 | 0 | 0 |
| | **Sub9** | 159 | 40 | 0 | 0 | **Sub9** | 177 | 66 | 0 | 0 |
| | **Sub10** | 84 | - | - | - | **Sub10** | 104 | - | - | - |
| | **Sub11** | 114 | 17 | 0 | 0 | **Sub11** | 79 | 36 | 0 | 0 |
| | **Sub12** | 177 | - | - | - | **Sub12** | 197 | - | - | - |
| | | | **average % overlap:** | | **4.45** | | | **average % overlap:** | | **0.00** |
| | | | | **sem** | **4.45** | | | | **+/-** | **0.00** |
| LO, ORS (SetB) | **Sub1** | 76 | 28 | 0 | 0 | **Sub1** | 97 | 29 | 0 | 0 |
| | **Sub2** | 78 | 11 | 0 | 0 | **Sub2** | 130 | 22 | 0 | 0 |
| | **Sub3** | 138 | 10 | 1 | 10.0 | **Sub3** | 121 | 36 | 0 | 0 |
| | **Sub4** | - | - | - | - | **Sub4** | - | - | - | - |
| | **Sub5** | 83 | 66 | 0 | 0 | **Sub5** | 54 | 13 | 0 | 0 |
| | **Sub6** | 128 | - | - | - | **Sub6** | - | 11 | - | - |
| | **Sub7** | 214 | 31 | 21 | 67.7 | **Sub7** | 187 | - | - | - |
| | **Sub8** | 96 | 59 | 0 | 0 | **Sub8** | 128 | 26 | 0 | 0 |
| | **Sub9** | 159 | 47 | 0 | 0 | **Sub9** | 177 | 30 | 0 | 0 |
| | **Sub10** | 84 | 24 | 0 | 0 | **Sub10** | 104 | - | - | - |
| | **Sub11** | 114 | 14 | 0 | 0 | **Sub11** | 79 | 20 | 0 | 0 |
| | **Sub12** | 177 | 46 | 0 | 0 | **Sub12** | 197 | 42 | 0 | 0 |
| | | | **average % overlap:** | | **7.77** | | | **average % overlap:** | | **0.00** |
| | | | | **sem** | **6.73** | | | | **sem** | **0.00** |

**Supplementary Table 5**: Analysis of voxel overlap between the occipital-partietal reachspace-preferring region and other classic lateral-dorsal ROIs. The superior parietal reachspace region is not shown, as it did not overlap with any ROIs.

| | | | | | | | | | | |
|---|---|---|---|---|---|---|---|---|---|---|
| Hand, ORS (SetA) | **Sub1** | 38 | - | - | - | **Sub1** | 67 | 33 | 0 | 0 |
| | **Sub2** | 122 | 28 | 0 | 0 | **Sub2** | 48 | 20 | 0 | 0 |
| | **Sub3** | 73 | 60 | 0 | 0 | **Sub3** | 113 | 127 | 2 | 1.6 |
| | **Sub4** | - | - | - | - | **Sub4** | - | - | - | - |
| | **Sub5** | 57 | 32 | 0 | 0 | **Sub5** | - | 27 | - | - |
| | **Sub6** | 54 | - | - | - | **Sub6** | 53 | - | - | - |
| | **Sub7** | 44 | - | - | - | **Sub7** | 74 | - | - | - |
| | **Sub8** | 118 | - | - | - | **Sub8** | 84 | 75 | 0 | 0 |
| | **Sub9** | 63 | 40 | 0 | 0 | **Sub9** | 114 | 66 | 0 | 0 |
| | **Sub10** | 91 | - | - | - | **Sub10** | 97 | - | - | - |
| | **Sub11** | 87 | 17 | 0 | 0 | **Sub11** | 114 | 36 | 0 | 0 |
| | **Sub12** | 51 | - | - | - | **Sub12** | 97 | - | - | - |
| | | | **average % overlap:** | | **0.00** | | | **average % overlap:** | | **0.27** |
| | | | | **sem** | **0.00** | | | | **sem** | **0.27** |

| | | | | | | | | | | |
|---|---|---|---|---|---|---|---|---|---|---|
| Hand, ORS (SetB) | **Sub1** | 38 | 28 | 0 | 0 | **Sub1** | 67 | 29 | 0 | 0 |
| | **Sub2** | 122 | 11 | 0 | 0 | **Sub2** | 48 | 22 | 0 | 0 |
| | **Sub3** | 73 | 10 | 0 | 0 | **Sub3** | 113 | 36 | 0 | 0 |
| | **Sub4** | - | - | - | - | **Sub4** | - | - | - | - |
| | **Sub5** | 57 | 66 | 0 | 0 | **Sub5** | - | 13 | - | - |
| | **Sub6** | 54 | - | - | - | **Sub6** | 53 | 11 | 0 | 0 |
| | **Sub7** | 44 | 31 | 0 | 0 | **Sub7** | 74 | - | - | - |
| | **Sub8** | 118 | 59 | 0 | 0 | **Sub8** | 84 | 26 | 0 | 0 |
| | **Sub9** | 63 | 47 | 0 | 0 | **Sub9** | 114 | 30 | 0 | 0 |
| | **Sub10** | 91 | 24 | 0 | 0 | **Sub10** | 97 | - | - | - |
| | **Sub11** | 87 | 14 | 0 | 0 | **Sub11** | 114 | 20 | 0 | 0 |
| | **Sub12** | 51 | 46 | 0 | 0 | **Sub12** | 97 | 42 | 0 | 0 |
| | | | **average % overlap:** | | **0.00** | | | **average % overlap:** | | **0.00** |
| | | | | **sem** | **0.00** | | | | **+/-** | **0.00** |

**Supplementary Table 5** (continued).
